# Brain-wide representations of olfactory behavior in *C. elegans*

**DOI:** 10.64898/2026.10.29.762753

**Authors:** Helena Casademunt, Hongruo Zhang, John Laver, Core Francisco Park, Sophia Kim, William Fredenberg, Wesley Hung, John Calarco, Mei Zhen, Aravinthan D.T. Samuel

## Abstract

Olfactory navigation emerges from dynamic interactions between sensory input, brain activity state and the animal’s own movements. In *C. elegans*, a compact nervous system enables brain-wide imaging in behaving animals exposed to controlled olfactory environments, with an undulatory locomotion that reduces navigation to transitions among forward runs, reversals, and turns. Here, we used tracking microscopy to record and compare brain-wide neural activity in crawling worms navigating spatial odor gradients or isotropic environments, and in immobilized worms responding to temporal odor pulses. Across conditions, we identified strongly correlated brain-wide activity patterns spanning olfactory sensory neurons, interneurons, and premotor interneurons. Comparisons between navigation in odor gradients and exploration in isotropic environments revealed a compact functional circuit linking olfactory sensory neurons, interneurons, and premotor neurons. Comparisons between different odors distinguished sensory activity relationships associated with odor identity from a shared sensorimotor circuit for navigational movement. Comparisons between crawling and immobilized animals revealed profound differences in motor-state transitions and brain-wide correlation structure, further demonstrating the dependence of sensory representations and motor decision-making on behavioral context. Our findings reveal a functional organization linking olfactory sensory activity to navigational movement and show how this organization depends on behavioral state and environmental context.

## Introduction

Animals navigate olfactory environments by generating movements oriented towards attractive odors and away from repellents. These navigational movements are both responses to odor gradients and active means of sampling them. Underlying neural circuits must both control navigational movement and evaluate movement-evoked sensory input along the animal’s trajectory.

Behavioral context – whether an animal is moving freely in an isotropic environment, moving toward a distant goal, or immobilized during sensory stimulation – may strongly influence sensorimotor processing in navigational circuits ^1^. From insects to mammals, the earliest stage of sensory processing can already be modified by locomotory state and recurrent neural activity. Such modulation has been proposed to help distinguish sensory changes generated by self-motion from those caused by the external environment ^2–4^. Consistently, behavioral context reshapes brain-wide functional organization, changing how activity is coordinated across neurons and brain regions. In mice, for example, neural activity becomes less correlated across neurons and brain regions as navigational tasks become more complex ^5^. In *Drosophila*, brain-wide activity is broadly reconfigured by behavioral states, including walking, resting, and grooming, as well as by external sensory input ^6,7^. In *C. elegans*, brain-wide activity is less tightly correlated in crawling animals than in immobilized or stationary animals ^8–10^.

Brain-wide recording offers a route to understanding how neural circuits integrate olfactory perception, decision-making, and navigational behavior through ongoing sensory and motor feedback. *C. elegans* is particularly well suited to this approach because it enables brain-wide recording with single-neuron resolution and comprehensive cellular identification ^11,12^, and because sensory input and motor output are only a few synapses apart in a compact sensorimotor network ^13,14^.

Previous recordings from selected candidate neurons in moving and immobilized *C. elegans* have revealed how individual subcircuits contribute to olfactory navigation ^15–17^. These subcircuits integrate signals from other sensory pathways that convey information about the animal’s movements and surroundings, as well as the recurrent motor feedback that modulates decision-making interneurons ^18,19^. How olfactory navigation emerges from a complete functional circuit – from sensory input to motor output embedded in the wider brain network – remains an open question.

Understanding olfactory processing requires brain-wide measurements of identified neurons throughout behavioral trajectories across diverse sensory environments and behavioral contexts. Recording many animals within each experimental condition is needed to distinguish differences in circuit dynamics across conditions ^20^. Here, we recorded brain-wide neural activity in *C. elegans* hermaphrodites as they crawled in spatial gradients of diacetyl (DA) or isoamyl alcohol (IAA), two well-studied, naturally occurring attractive odors ^21^, and in isotropic environments. We compared these recordings with brain-wide activity in immobilized worms exposed to temporal odor pulses using microfluidics ^22,23^. NeuroPAL ^12^ and complementary sparse multicolor markers allowed us to identify the same neurons across animals. Volumetric imaging ^24,25^, combined with computational tools for neuronal identification and tracking ^12,24,26,27^, enabled repeated measurements of the same set of neurons across experimental replicates. These measurements allowed us to establish patterns of circuit dynamics that reliably differentiate sensory environments and behavioral contexts.

We found that neural dynamics adapted to both sensory environments and behavioral contexts at multiple levels, from individual neurons to brain-wide activity patterns. Worms that navigated spatial odor gradients and immobilized worms that exhibited fictive responses to odor pulses showed distinct patterns of brain-wide correlations. As animals moved through odor gradients, olfactory neurons adapted to local odor conditions, allowing them to encode concentration changes. In crawling worms, comparisons of relationships among sensory neurons, interneurons, and motor neurons across sensory environments revealed a functional circuit whose dynamics distinguished chemotaxis from isotropic exploration. This olfactory navigational circuit integrates odor signals, motor feedback, and additional sensory information about the physical environment to guide odor-evoked decisions toward olfactory goals.

## Results

### Brain-wide imaging of the olfactory response

We recorded brain activity in adult hermaphrodites navigating spatial odor gradients and isotropic environments using a spinning-disc confocal microscope customized for multicolor, multi-neuronal volumetric imaging with semi-automated neural tracking (Fig. 1A) ^24,25^.

**Figure 1.**
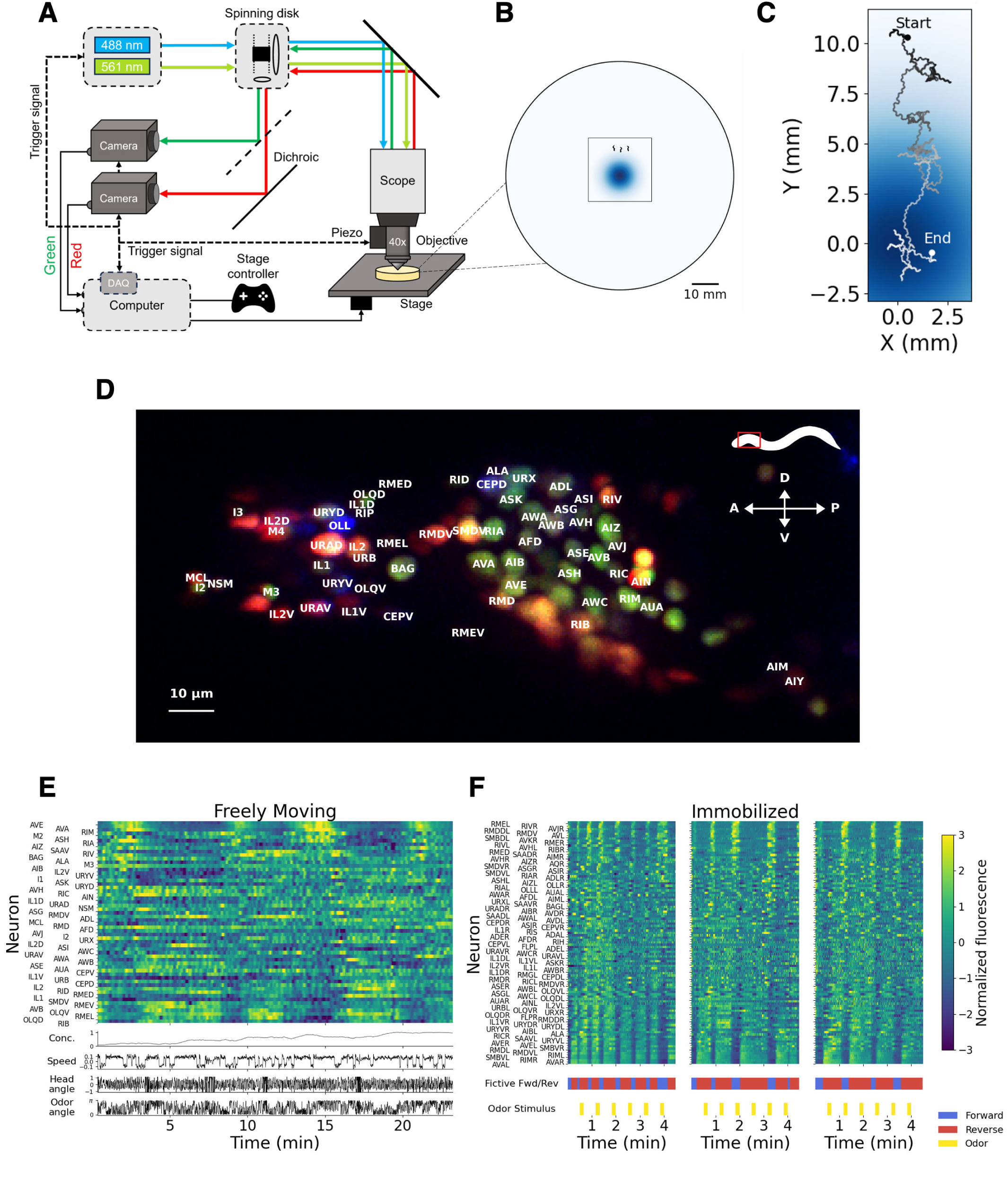
(**A**) A tracking microscope simultaneously records navigational behavior and neuron activity based on nuclear-localized calcium indicator (GCaMP6s) and multi-color neuronal markers to resolve the location and identity of nearly every neuron in the anterior brain. (**B**) Spatial gradient on the agar plate at the start of an experiment showing the odorant (diacetyl) concentration measured in arbitrary units; darker color indicates higher concentration. The square shows typical 25×25mm coverslip placement with respect to the plate and odor gradient. Worms are not drawn to scale. The experiment is the same as in (C) and (E). (**C**) Trajectory on a diacetyl gradient, for same worm plotted in (E). Darker color indicates higher concentration of odorant; colors match (B). Origin (0,0) corresponds to odor source position. (**D**) Maximum intensity projection image of the head of an ADS1046 worm with neuron labels. ADS1046 expresses three sparse markers in addition to pan-neuronal GCaMP6s and mNeptune: cyOFP1 (pseudocolored green), mScarlet (pseudocolored red), and BFP (pseudocolored blue). The image shows the sparse markers only, omitting GCaMP6s and mNeptune. Image taken with a 60x water immersion objective. A: anterior; P: posterior; D: dorsal; V: ventral. (**E**) Neuron activity heatmap for all annotated neurons in a single recording of an animal crawling towards diacetyl, corresponding to the trajectory in (C). Only neurons that were identified are shown. Concentration (a.u.), velocity (mm/s), head angle (a.u.), and angle to odor (rad) measurements are shown using the same time axis. Neuron activity is shown as z-score normalized fluorescence, with corresponding color bar shown in (F). (**F**) Neuron activity heatmap for all annotated neurons in an immobilized recording of an animal experiencing diacetyl pulses (10^−5^ dilution) in a microfluidic device. Neuron activity was recorded for three sequences of 6 pulses each. Odor stimulus pattern and fictive behavior annotations (see Microfluidics experimental design in Methods) are also shown. Neuron activity is shown as z-score normalized fluorescence.

Radial spatial odor gradients for freely navigating animals were generated on agar pads by allowing odors to diffuse from a point source using standard protocols ^28,29^. In brief, odors (diacetyl (DA) at 1% v/v or isoamyl alcohol (IAA) at 10% v/v) were applied to agar pads, and gradients formed by diffusion over ∼ 3 hours (Fig. S1C).

Animals were placed approximately 1 cm from the original odor source at the start of each experiment (Fig. 1B). Immediately before recording, the imaging area was covered with a coverslip. Brain activity and behavior were tracked continuously for 5-30 minutes as animals crawled toward the odor source (Fig. 1C). All recorded animals showed net movement toward higher odor concentrations (12/12 for DA and 16/16 for IAA).

For neuron identification, we adopted a simplified version of the combinatorial NeuroPAL color-code strategy (Fig. 1D). The original NeuroPAL transgene established a stereotyped multicolor fluorescence map for the entire neuronal population ^12^ but severely impaired the animal’s motility during olfactory navigation ^12^. We reduced marker expression in three overlapping subsets of neurons to preserve robust olfactory navigation (Fig. 1C; see also Figs S2, S3) while maintaining sufficient color diversity to identify nearly all neurons in the head.

In addition to recording brain activity, we extracted from the same recordings behavioral features relevant to olfactory navigation, including velocity, heading angle relative to the body centerline, distance and heading angle relative to the odor peak, and estimated local odor concentration (Fig. 1E). Positive and negative velocities corresponded to forward and backward movement, respectively.

We also recorded brain activity in *C. elegans* immobilized in microfluidic devices with pulses of DA delivered (Fig. S1A,B) ^23^. Fictive forward and backward motor states were inferred from neural activity of AVA, a premotor interneuron with higher activity associated with backward movement and lower activity with forward movement ^30^, similar to earlier studies (Fig. 1F) ^31–33^. We applied these datasets to assess effects of motor states on brain activities and to extract features of neuronal activity patterns associated with olfactory navigation.

### Olfactory navigation strategies revealed by movement statistics in spatial odor gradients

We first examined the behavioral strategy used for olfactory navigation by analyzing movement statistics in spatial odor gradients and in isotropic environments lacking odor.

In both olfactory gradients and isotropic environments, within each trajectory, worms alternated between forward and backward movement. Sustained forward runs were interrupted by turning maneuvers. These were either single turns, consisting of a brief reversal followed by forward movement, or sequences of two or more reversal-turn events in rapid succession called pirouettes (Fig. 2C, D). Pirouette length, measured by the number of reversal-turn events, varied in both environments (Fig. 2C).

**Figure 2.**
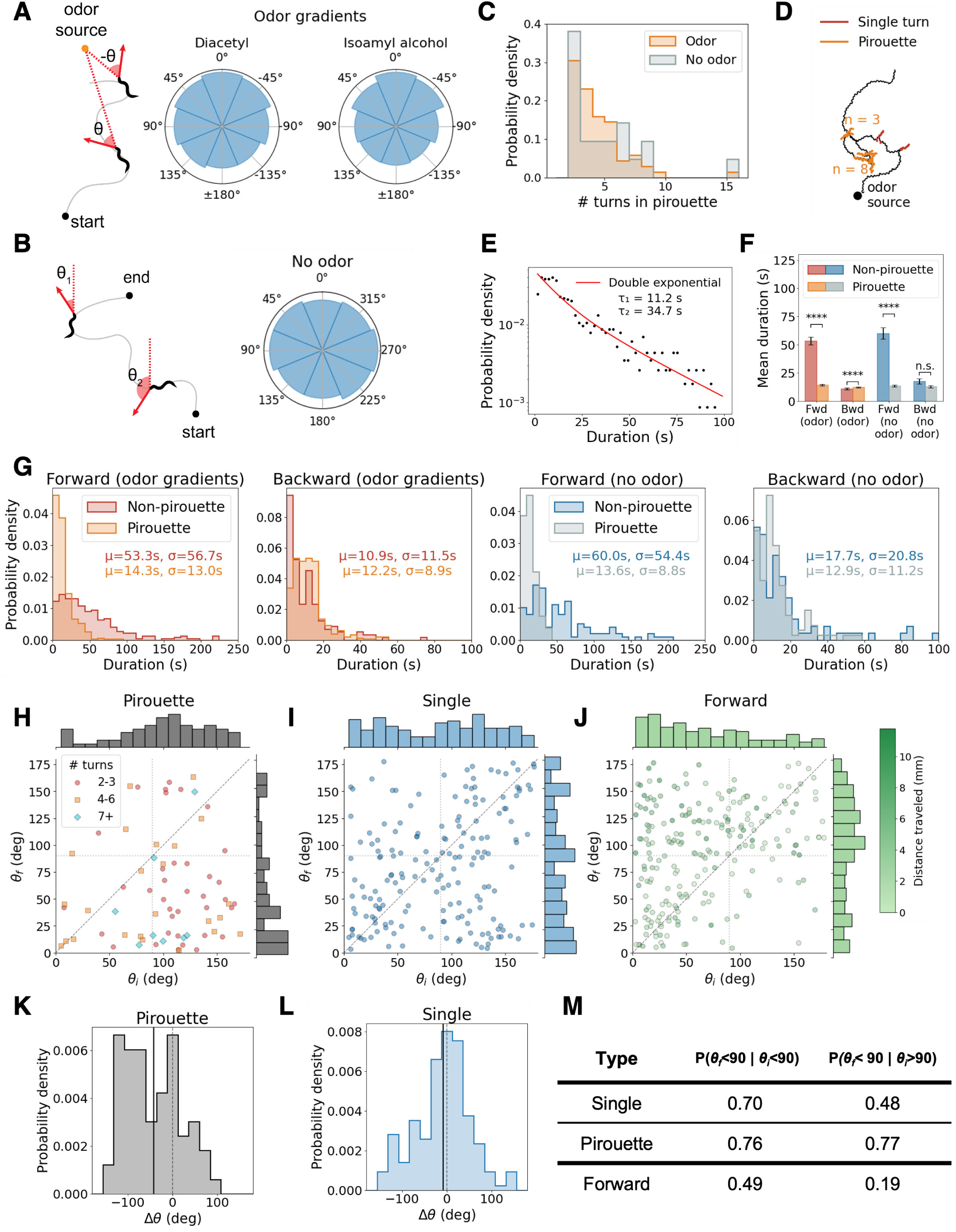
(**A**) Distribution of angles between animal head and odor source for worms navigating in odor gradients of diacetyl (N = 12) and isoamyl alcohol (N = 16). Schematic indicates how the angle is computed with respect to the odor source. The arrow indicates the direction the worm head points to. Angles of 0° indicate the animal head points directly towards the odor source and angles of 180° indicate the animal head points directly away. (**B**) Distribution of angles between animal head and lab reference frame for worms navigating in isotropic environments (N = 13). Schematic indicates how the angle is computed with respect to the lab frame. The arrow indicates the direction the worm head points to. (**C**) Distribution of number of turns in a pirouette for all animals navigating in odor gradients (69 pirouettes, 28 animals) and all animals in isotropic environments without odor (21 pirouettes, 13 animals). (**D**) Example trajectory of an animal navigating in an isoamyl alcohol gradient, showing the difference between single turns and pirouettes. n indicates the number of reversals in each pirouette. (**E**) Distribution of forward segment durations for all animals navigating in odor gradients (574 forward runs, 28 animals). Run durations are binned into 50 bins of 2-s width. Runs longer than 100 seconds were excluded. The red line shows a double exponential fit, with R^2^(log) = 0.907. (**F**) Mean duration of forward and backward segments inside or outside a pirouette, separately for worms navigating in odor gradients vs. in no-odor isotropic environments. Statistical comparisons between non-pirouette and pirouette segment lengths were performed using two-sided Mann-Whitney U tests. Forward segments were significantly shorter for pirouettes than for non-pirouettes both in odor gradients (p = 1.6 ×10^−43^, ****) and in isotropic environments (p = 2.8 ×10^−19^, ****). Backward segment lengths also differed significantly under odor (p = 8.2 ×10^−5^, ****), but were not significantly different in the no-odor condition (p = 0.40, n.s.). Significance is indicated as *p < 0.05, **p < 0.01, ***p < 0.001, ****p < 10^−4^; n.s., not significant. Odor gradients (28 animals): 605 forward segments (347 pirouette, 258 non-pirouette), 578 backward segments (418 pirouette, 160 non-pirouette). No odor (13 animals): 210 forward segments (93 pirouette, 117 non-pirouette), 198 backward segments (116 pirouette, 82 non-pirouette). (**G**) Distribution of forward and backward segment durations inside or outside a pirouette, separately for worms navigating odor gradients vs. in no-odor isotropic environments. Text shows the mean *µ* and standard deviation *σ* of the distributions. Segment counts are the same as in (F). (**H**) Angle *θ* between head and odor source before and after a pirouette for worms navigating in odor gradients (28 animals, 69 pirouettes). *θ_i_* is the mean angle to odor for the 5 seconds before the first reversal in a pirouette and *θ_f_* is the mean angle to odor in the 5 seconds after the end of the last reversal in the pirouette. Markers show number of turns in the pirouette. (**I**) Angle *θ* between head and odor source before and after a single reversal for worms navigating in odor gradients (28 animals, 176 turns). *θ_i_* is the mean angle to odor for the 5 seconds before the reversal and *θ_f_* is the mean angle to odor in the 5 seconds after the end of the reversal. (**J**) Angle *θ* between head and odor source at the start and end of a forward run (excluding forward segments within a pirouette) for worms in odor gradients (28 animals, 221 segments). *θ_i_* is the mean angle to odor in the first 5 seconds of the forward segments and *θ_f_* is the mean angle to odor in the last 5 seconds of the forward segment. We exclude any forward segments with duration less than 10 seconds. Marker color shows the distance traveled during the forward segment (mm). (**K**) Distribution of angle change during a pirouette for worms navigating in odor gradients. Angle change is measured as Δ*θ* = *θ_f_* − *θ_i_*. Plot uses the same data and definitions as in (H). Mean = −43.0° (69 pirouettes). (**L**) Distribution of angle change during single turns for worms navigating in odor gradients. Angle change is measured as Δ*θ* = *θ_f_* − *θ_i_*. Plot uses the same data and definitions as in (I). Mean = −8.6° (176 single turns). (**M**) Table showing fraction of pirouettes, single turns, or forward segments that end facing the odor direction, separated by whether they start facing the odor or away from it. Data combines all 28 animals in odor gradients, with 176 single turns, 69 pirouettes, and 221 forward segments (excluding segments within a pirouette). See (H), (I), (J), for angle definitions.

Forward movements during pirouettes were shorter than those outside pirouettes. Accordingly, the distribution of forward-movement durations in crawling worms was well fit by a double exponential, with the shorter time constant corresponding to forward segments within pirouettes (Fig. 2E). Pirouettes therefore constitute a behavioral state with shorter forward movements than the non-pirouette state, consistent with previous work ^28,34,35^. This pattern was also evident in both environments (Fig. 2F,G; Fig. S5A,B). By contrast, backward-movement statistics differed only weakly between pirouette and non-pirouette states in either condition.

When worms navigate olfactory gradients, consistent with previous studies of chemotactic navigation ^36–38^, forward runs are longer when initiated while the animal is oriented toward increasing odor concentrations (*θ_i_ <* 90^◦^) than toward decreasing odor concentrations (*θ_i_ >* 90^◦^) (Fig. S5C). This modulation reflects the biased-random-walk component of olfactory navigation, in which organisms prolong forward movements that carry them toward more favorable conditions ^28,39^.

During olfactory navigation in spatial gradients, worms spent more time with their heads oriented toward the odor source than in other directions (Fig. 2A; Fig. 2B). Pirouettes contributed to this directional bias by reorienting the head toward favorable directions (Fig. 2H-M). On spatial odor gradients, most pirouettes were initiated while the animal was heading toward decreasing odor concentration (*θ_i_ >* 90^◦^) and ended with headings toward increasing odor concentration (*θ_f_ <* 90^◦^), regardless of the number of reversal-turns they contained (Fig. 2H). The mean change in heading angle was similar for pirouettes of different lengths (Fig. S5D), while the change in heading angle per reversal-turn was variable and did not depend on the total number of reversal-turns within the pirouette (Fig. S5D). Pirouettes end with favorable headings regardless of whether they began with favorable or unfavorable headings (Fig. 2M).

The individual reversal-turn events within a pirouette appeared to be random exploratory steps. The first reversal-turn was usually initiated when the animal was heading in an unfavorable direction (*θ_i_ >* 90^◦^) and produced a variable change in heading angle (Fig. S5E,F). Additional reversal-turns also occurred across a broad range of heading angles, consistent with repeated sampling until exiting the pirouette (Fig. S5E,F). Consistent with this interpretation, the final reversal-turn tended to end with a favorable heading (Fig. S5E,F). As a result, the net change in heading angle across a pirouette was biased toward a reduction in heading angle relative to the gradient (Fig. 2K).

Single reversal-turn events were less effective than pirouettes at orienting the animal toward a favorable heading (Fig. 2I). This likely reflects angular persistence, because the net change in heading angle across a single turning maneuver was typically *<* 90^◦^ (Fig. 2L). Accordingly, when the animal began a single reversal-turn with a favorable heading (*θ_i_ <* 90^◦^), it usually maintained a favorable heading afterward (*P* (*θ_f_ <* 90^◦^ | *θ_i_ <* 90^◦^) = 0.70) (Fig. 2M) and single reversal-turn events that began with an unfavorable heading were not substantially more likely to end with a favorable heading (*P* (*θ_f_ <* 90^◦^ | *θ_i_ >* 90^◦^) = 0.48) (Fig. 2M).

Sustained forward runs usually began with the animal heading in a favorable direction (*θ_i_ <* 90^◦^), often because they followed pirouettes or single reversal-turn events, but they could end at any heading angle. Forward runs that began with an unfavorable heading (*θ_i_ >* 90^◦^) were not especially likely to end with a favorable heading, whereas shorter runs that began in unfavorable directions tended to remain pointed in unfavorable directions (Fig. 2J,M). Thus, we found no evidence for klinotaxis, or gradual steering during runs, in our experimental conditions ^36^.

When pirouettes, single reversal-turn events, and forward runs were analyzed separately during navigation in DA and IAA, we found no significant differences in their behavioral statistics (Fig. S6A–E). Thus, worms appeared to use essentially the same navigational strategy to ascend gradients of both odors in our experimental conditions.

### Fictive olfactory responses have both shared and divergent features compared with responses in behaving animals

In immobilized animals, we assessed responses to odor pulses by measuring transitions between inferred forward and backward motor states. By analogy with biased random walk navigation in freely moving animals, during pulses of attractive odors, odor onset would be expected to favor a transition from fictive reversal to fictive forward movement, whereas odor offset would be expected to favor the opposite transition.

This feature was only partially reflected by inferred motor-state transitions. We measured AVA activity in immobilized animals subjected to 10 s pulses of DA separated by 30 s intervals (Fig. 3A). When AVA was in a high-activity state, corresponding to fictive backward movement, animals transitioned to the fictive forward state during the first 5 seconds following odor onset in only 25% of cases, whereas odor offset was associated with this transition in 13% of cases. Similarly, when AVA was initially in a low-activity state, corresponding to fictive forward movement, animals transitioned to the fictive backward state within 5 seconds of odor offset in 26% of cases (Fig. 3B), whereas odor onset was associated with such transitions in 11% of cases.

**Figure 3.**
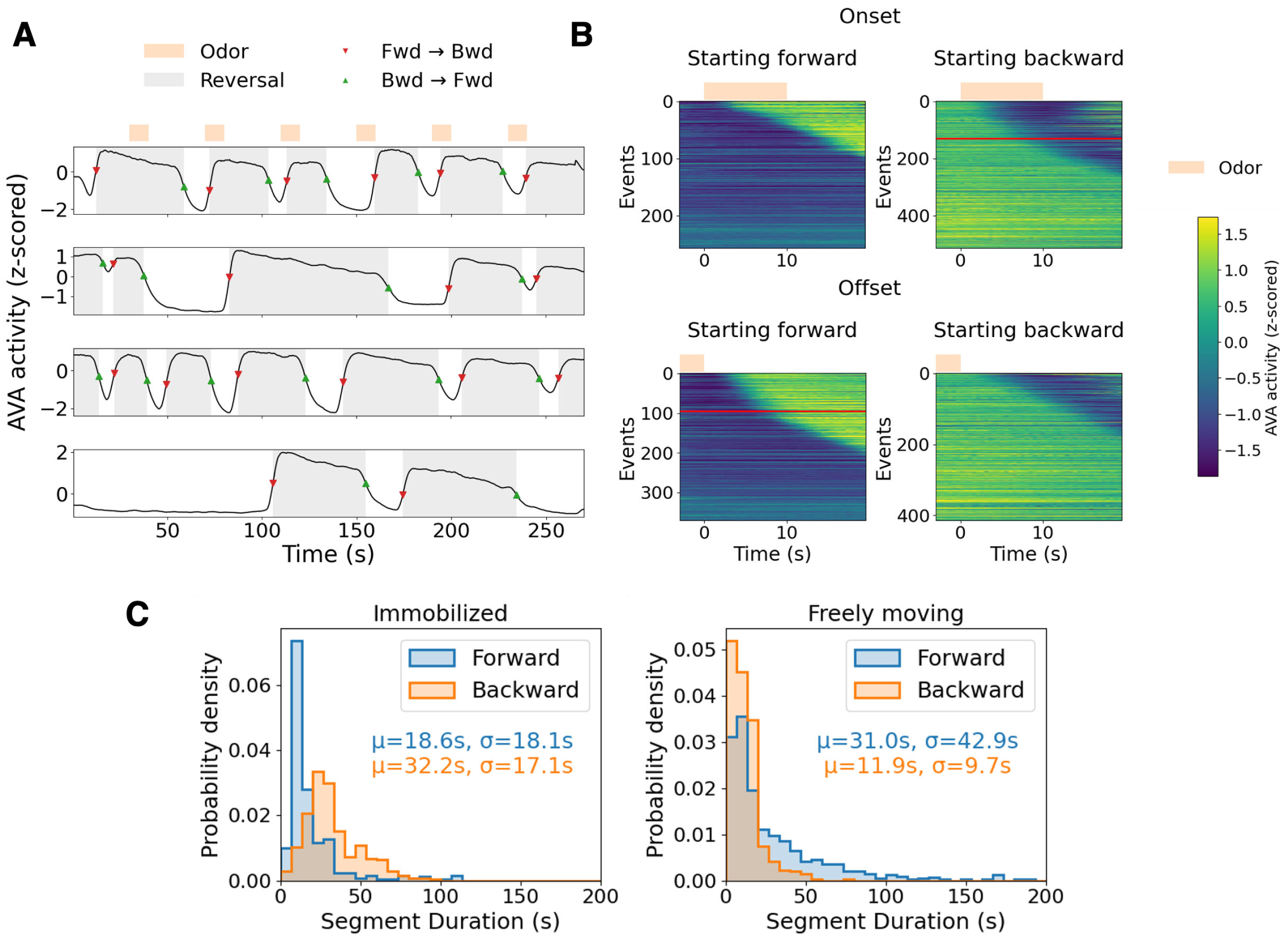
(**A**) AVA activity from four runs from different worms, showing how fictive reversals and forward (fwd) or backward (bwd) transitions are determined from neuron activity. AVA activity is z-score normalized fluorescence. Orange rectangles at the top of the plot show diacetyl pulse stimulus for all runs. (**B**) AVA activity aligned to odor onset or offset. Left plot shows activity for events where the worm is going forward before odor onset/offset, and right plot shows activity for events where the worm is going backward. The red line separates events that are classified as odor-evoked transitions, i.e. events that happen in the first 5 seconds after onset or offset. At odor onset, 257 events started forward and 514 started backward. 130/514 (25.3%) were classified as transitions from backward to forward. At odor offset, 370 events started forward and 414 started backward. 95/370 (25.7%) were classified as transitions from forward to backward. Data come from 153 6-pulse runs across 63 animals. (**C**) Forward and backward segment duration distribution in immobilized experiments (diacetyl pulses, 63 animals) and freely moving experiments (odor gradients only, 28 animals). Immobilized: 363 forward segments, 376 backward segments; freely moving: 605 forward runs, 578 reversals.

The mean durations of fictive forward and backward states in immobilized worms differed markedly from the durations exhibited by crawling animals (Fig. 3C,D). In crawling worms, forward runs tended to be longer than backward runs. In immobilized worms, by contrast, fictive backward states were much longer than fictive forward states. Previous imaging studies of AVA likewise reported sustained periods of fictive reversal activity ^32,33^. This pattern may reflect a response to immobilization itself, with animals confined in a microfluidic chip attempting prolonged backward wriggling expressed as sustained AVA activity.

### Brain-wide correlations reveal shared and context-dependent activity patterns

To assess neuronal interactions throughout the brain across different behavioral contexts, we computed neuron-neuron and neuron-behavior correlations for four conditions: microfluidic-immobilized animals exposed to DA pulses, animals crawling in DA gradients, animals crawling in IAA gradients, and crawling animals in isotropic environments. To identify groups of functionally related neurons, we constructed consensus correlation matrices by pooling data across experiments within each condition. Consistent with previous reports, hierarchical clustering of neuron-neuron correlations in immobilized worms revealed a block-diagonal structure with strongly anti-correlated modules corresponding to fictive forward and backward motor states (Fig. 4A). The fictive forward state was strongly associated with elevated activity in several neurons, including AVB, RME, RIB, and AIY ^32,33,40,41^. The fictive backward state was strongly associated with elevated activity of interneurons including RIM, AVE, and AIB ^16,32,33,42^. In immobilized worms, correlations between individual neuronal activity and fictive forward or backward state extended broadly throughout the brain, consistent with previously described global brain states.

**Figure 4.**
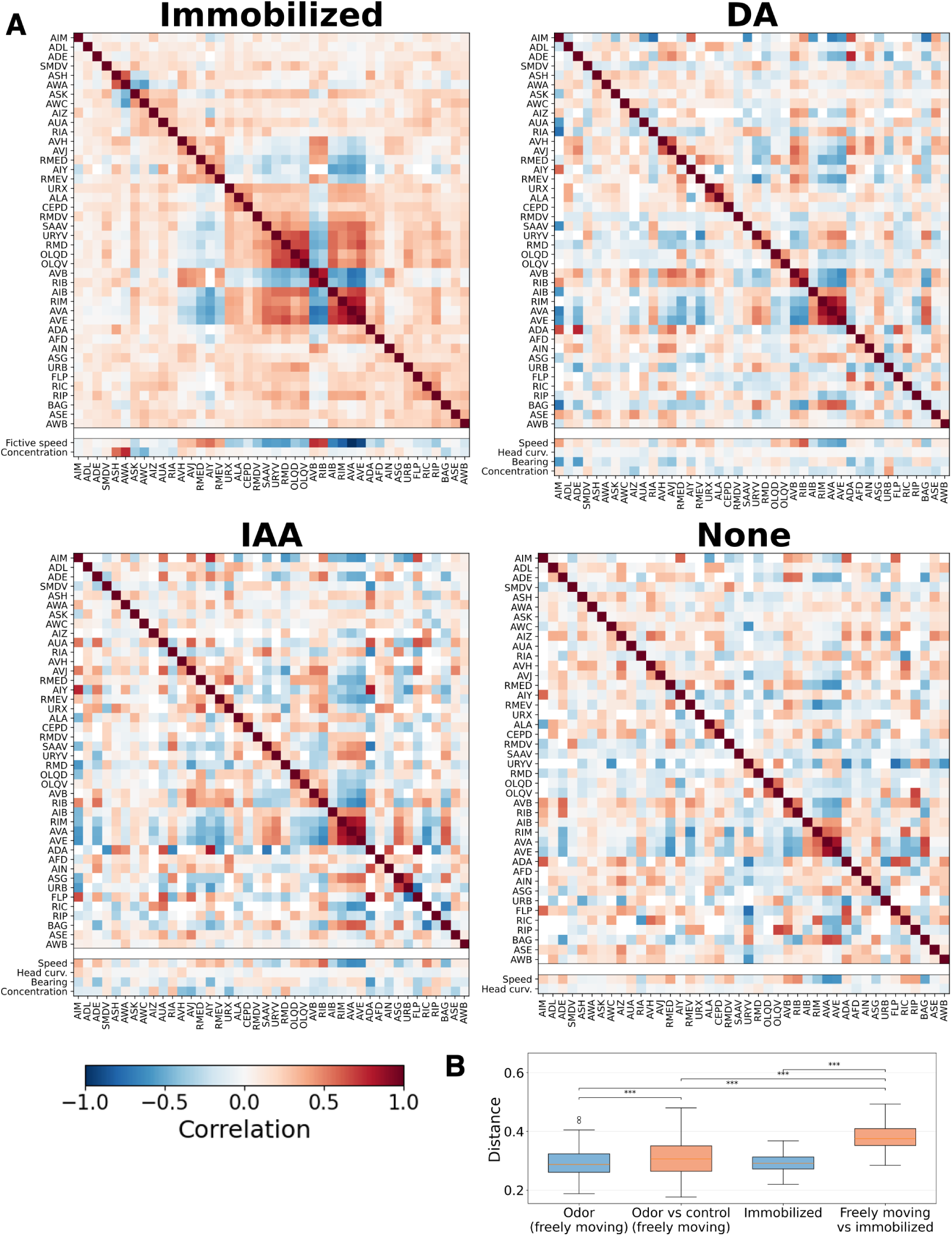
(**A**) Pairwise correlation matrices for recorded neurons and stimulus and behavioral variables, averaged across all worms experiencing each condition. Matrices for immobilized worms (8 animals), and crawling worms in gradients of diacetyl (DA; 12 animals), isoamyl alcohol (IAA; 16 animals), or no odorant (13 animals) are shown separately, sharing a color bar. Stimulus and behavioral variables shown vary by stimulus or experiment type. Matrices are sorted using hierarchical clustering of the immobilized worm matrix. Only neurons present in at least one dataset for each stimulus are included. For immobilized worms, left and right neurons are averaged before computing the correlation. (**B**) Euclidean distance distributions between individual neuron-to-neuron correlation matrices. Distance is the Frobenius norm of the difference between two matrices, normalized by the number of neurons. Distances are computed between different worms for the following conditions, in plotted order from left to right: (i) two freely moving (FM) worms in odor gradients (even if the odorant is different), (ii) two freely moving worms where one is in an odor gradient and the other is in a no-odor environment, (iii) two immobilized worms experiencing odor pulses, (iv) a freely moving worm in an odor gradient and an immobilized worm experiencing odor pulses. Statistical significance was assessed using two-sided Mann-Whitney U tests. FM vs. immobilized distances were significantly greater than all other distributions (p < 10^−14^ in all cases). Within-odor distances were significantly smaller than odor vs. no stimulus distances (p = 5.90 ×10^−5^). Within-odor and within-immobilized distances did not differ significantly (p = 0.80). *p < 0.05, **p < 0.01, ***p < 0.001.

When we examined cross-correlation matrices from crawling animals, with neurons ordered according to the hierarchical clustering obtained from immobilized worms, we observed a substantial reorganization of brain-wide activity patterns. Consistent with previous studies in moving worms, forward locomotion in crawling animals was strongly correlated with elevated activity in AVB and AIY ^32,41^, whereas backward locomotion was strongly correlated with AVA, RIM, AVE, and AIB ^32,41,42^. Correlations between chemosensory neurons like AWA, AWC, ASH also differed. Correlations were stronger for immobilized worms, where sensory neurons responded more uniformly to pulses than to actively sensed odor gradients in crawling worms. Across the rest of the brain, correlations with forward and backward movement were more heterogeneous and diverse.

Notably, cross-correlation matrices from crawling animals, whether in DA gradients, IAA gradients, or isotropic environments, were much more similar to one another than to those from immobilized worms. We quantified this internal consistency by calculating pairwise Euclidean distances between the consensus correlation matrices from each experimental condition (Fig. 4B). Distances were smaller among the three freely moving conditions and larger between freely moving conditions and the immobilized condition. By recomputing the hierarchical clustering from crawling animals and using that ordering to replot the cross-correlation matrix for immobilized worms (Fig. S7), we also observed distinct organization of brain-wide activity in crawling versus immobilized worms.

The similarity among cross-correlation matrices from animals navigating DA gradients, IAA gradients, and isotropic environments suggests that mobility versus immobilization is a dominant factor organizing brain-wide activity, whereas differences related to olfactory navigation, isotropic search, or odor identity are localized to relatively small numbers of neurons.

We also compared neuron-neuron and neuron-behavior correlations across individual datasets within each experimental condition. Within-condition Euclidean distances between pairs of datasets from the same crawling condition were distributed similarly to distances between different crawling conditions (Fig. S8A) and to distances among immobilized datasets (Fig. S8B). The statistically significant differences that remained could be attributed to small numbers of neurons, as described below (Fig. S8C). The overall similarity of cross-correlation matrices across datasets within a condition supports both the consistency of neuronal identity assignments and the high signal-to-noise ratio of the activity measurements.

Lastly, we used principal component analysis to compare how variance in brain-wide activity was distributed across collective activity patterns in immobilized and crawling animals. The cumulative variance explained by successive principal components was similar across conditions (Fig. S8D). Thus, the reorganization of neuron-neuron correlations was accompanied by a similar concentration of activity variance in the leading principal components. Hallinen et al. reported a similar qualitative difference in brain-wide activity patterns between crawling and immobilized recordings from the same animal ^8^, but without identifying individual neurons and therefore without the ability to compare homologous neurons across animals.

### Reconfiguration of brain-wide activity by immobilization

To identify specific cases of functional reconfiguration between identified neurons in crawling and immobilized animals, we computed the correlation between each neuron’s activity and forward locomotion, real for crawling animals and fictive for immobilized animals (Fig. 5A). As expected, AVA was the neuron most negatively correlated with forward locomotion in crawling animals, whereas AVB was among the neurons most positively correlated with forward locomotion in crawling animals and fictive forward locomotion in immobilized animals. Many neurons associated with forward and backward locomotion retained similar qualitative relationships to motor state across conditions. However, individual neurons differed between crawling and immobilized animals in the strength—and sometimes the sign—of their correlations with motor state (Fig. 5A).

**Figure 5.**
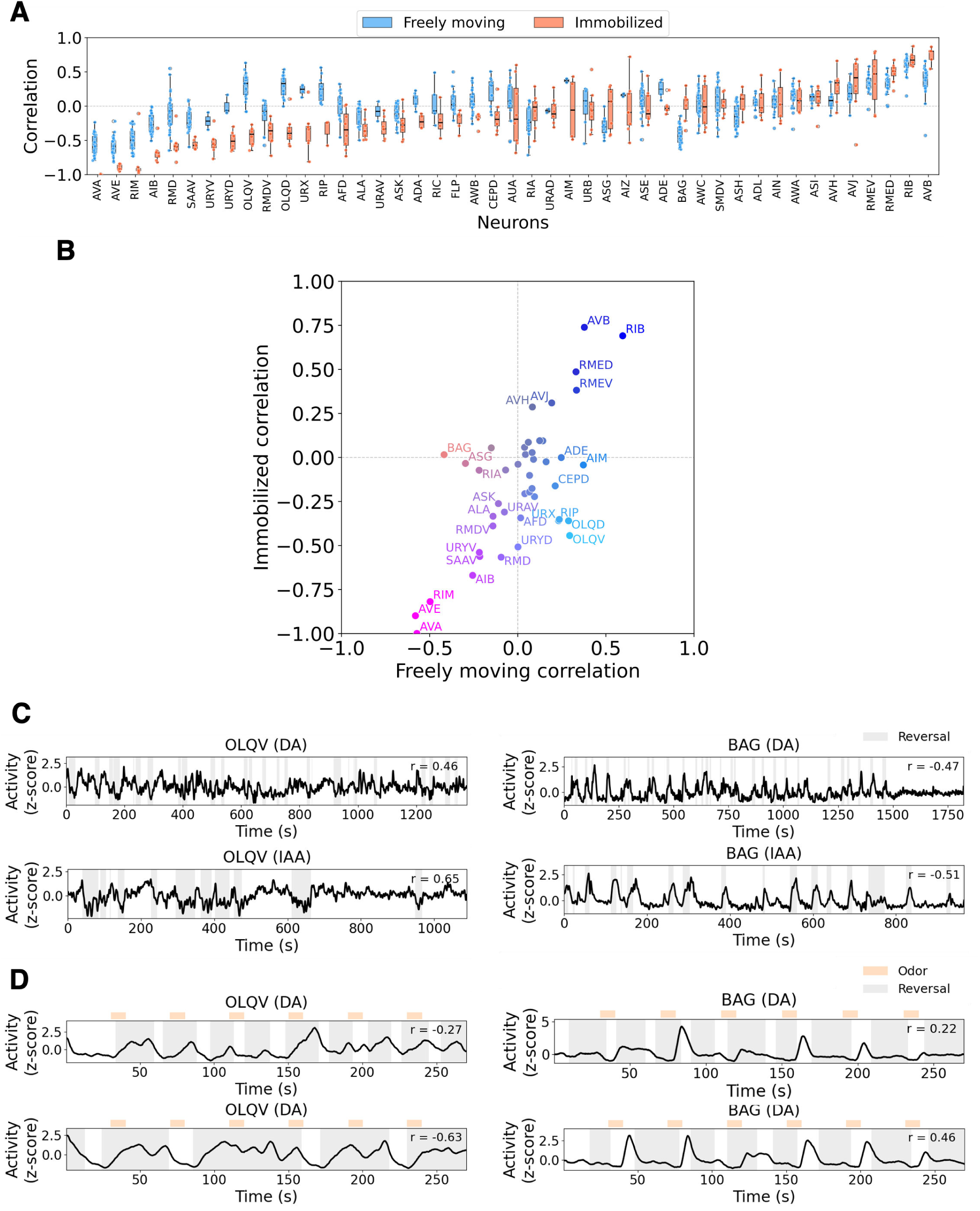
(**A**) Correlation of all neurons with velocity or fictive velocity, comparing distributions in freely moving (odor gradients only, 28 animals) and immobilized worms (8 animals). Fictive velocity is defined as the negative of AVA activity. For immobilized worms, left and right neurons are averaged before computing the correlation. Only neurons included in at least 2 datasets for each condition are shown. (**B**) Correlation of neuron activity with velocity in freely moving vs. fictive velocity in immobilized worms, showing the means for the correlation distributions in (A). Color is based on plotted correlation values and is used to help visualize correlation differences. Labels of neurons with small absolute correlation values are not shown for clarity. (**C**) Sample traces of OLQV and BAG in freely moving worms in odor gradients of diacetyl or isoamyl alcohol. Gray shading indicates reversals. Correlation *r* between neuron activity and velocity is shown for each trace. (**D**) Sample traces of OLQV and BAG in immobilized worms. Left and right neurons are averaged. Gray shaded area indicates fictive reversals. Orange regions show odor pulses. Correlation *r* between neuron activity and fictive velocity is shown for each trace.

A prominent example of this context-dependent reconfiguration involved the OLQ sensory neurons, which have been implicated in exploratory head movements during foraging and in nose-touch avoidance ^43–45^. OLQ exhibited one of the strongest sign reversals in correlation structure: its activity was positively correlated with forward locomotion in crawling animals but negatively correlated with forward locomotion in immobilized animals (Fig. 5B-D). Because OLQ also mediates TRPA1-dependent mechanosensation and nose-touch-evoked head withdrawal ^44^, this shift suggests that OLQ may contribute to the initiation or prolongation of backward bouts in immobilized animals by detecting mechanical constraint at the head.

A second example involved the aerotaxis circuit. The BAG sensory neuron detects CO_2_ ^46,47^, and in crawling animals its activity increases during reversals, likely reflecting detection of self-generated CO_2_ plumes during locomotion ^48^. Consistent with this idea, BAG activity was strongly correlated with backward movement in crawling animals (Fig. 5C). In immobilized animals, however, BAG no longer tracked forward-backward motor state and was instead associated with odor removal (Fig. 5D). Because BAG lacks sensory endings needed to directly detect external volatile odors, this shift suggests that BAG may be recruited into the olfactory circuit through synaptic or extrasynaptic signaling from primary olfactory receptor neurons ^49,50^.

Together, these results show that immobilization drives substantial reorganization of brain-wide activity. Neurons such as OLQ and BAG adopt distinct circuit roles in crawling and immobilized animals, with OLQ shifting its association with motor state and BAG functioning as a sensory neuron in the aerotaxis circuit but as an interneuron-like component of the olfactory circuit.

### Adaptive sensory thresholds encode odor changes during navigation

The olfactory sensory neurons AWA and AWC are broadly tuned to volatile odorants and play central roles in olfactory chemotaxis ^23,51,52^. Early ablation studies showed that chemo-taxis to DA is more sensitive to loss of AWA, whereas chemotaxis to IAA is more sensitive to loss of AWC ^51^. In our brain-wide analysis, one of the strongest spatially localized differences between animals navigating DA or IAA gradients and animals in isotropic environments was the relationship between AWA activity and forward locomotion. Previous studies in immobilized worms showed that AWA is activated by increases in DA or IAA concentration ^23,52,53^, whereas AWC is typically inhibited by increases in odor concentration, including DA and IAA ^15,23,52^.

Consistent with these earlier results, DA pulses in immobilized worms evoked increased AWA activity and decreased AWC activity (Fig. 6A) ^15^. Accordingly, and consistent with the biased random walk model, AWA activity in crawling animals was positively correlated with forward locomotion during gradient navigation but uncorrelated with locomotion in isotropic environments (Fig. 4A).

**Figure 6.**
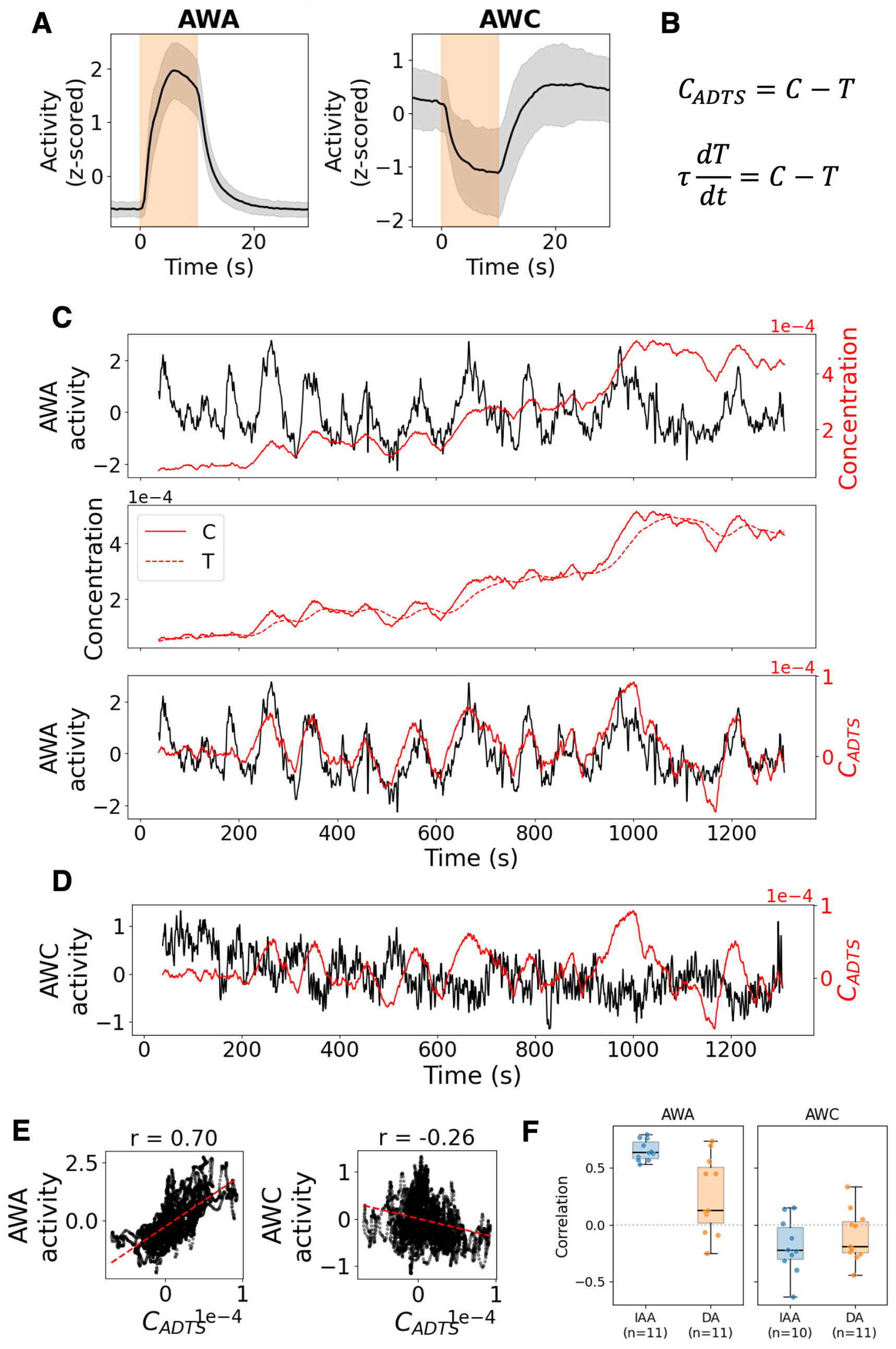
(**A**) Mean response to diacetyl pulse in microfluidics of two sensory neurons AWA and AWC. Gray shaded area shows the standard deviation. Orange background shows the odor pulse. Both neurons were recorded in 8 animals, for 144 total pulses. (**B**) Equations defining the ADTS model. *C* represents measured concentration, *T* is a slowly-relaxing threshold, *t* is time, and *τ* is a time constant that is fitted for each dataset. In subsequent analysis, we use the mean time constant for each odor (calculated after removing any datasets where the fit did not converge). We found a mean time constant of *τ* = 29.6 seconds for diacetyl and *τ* = 34.5 seconds for isoamyl alcohol. (**C**) (Top) AWA and raw concentration on the same plot for a worm navigating towards isoamyl alcohol. Every time concentration goes up, so does AWA, the magnitude of concentration changes grows as the worm approaches the source. (Middle) Relationship between *C* and *T* from the ADTS model. We show *T* using the mean time constant for isoamyl alcohol, *τ* = 34.5 seconds. (Bottom) AWA activity and *C_ADTS_* plotted together. Data are the same as in (E), showing correlation of 0.70 between AWA activity and *C_ADTS_*. (**D**) AWC activity plotted with *C_ADTS_*. They are anti-correlated, with a correlation of -0.26 (shown in (E)). (**E**) Scatter plots showing linear relation between *C_ADTS_* and AWA activity or AWC activity for a single worm, same as in (C) and (D). Correlation is positive for AWA and negative for AWC. Red dashed lines show a linear fit to the data. (**F**) Distribution of correlation coefficients between AWA or AWC activity and *C_ADTS_*, separated by odor. IAA datasets show a stronger relationship between neuron activity and *C_ADTS_*. Number of AWA and AWC measurements for each odor shown as *n*.

In our experiments, worms climbed spatial DA and IAA gradients spanning an estimated 10–100× range in odor concentration over a distance of about 1 cm. Olfactory sensory neurons must therefore detect the small concentration changes produced by self-motion while remaining sensitive across a broad dynamic range. Consistent with this requirement, aligning sensory activity to the time-varying local concentration experienced by crawling animals revealed transient, adaptive increases in AWA activity as animals moved up spatial gradients toward progressively higher concentrations of either odor (Fig. 6B; Fig. S9A).

To relate AWA activity to the time-varying concentration experienced by crawling animals, we adapted an “adaptive threshold model” ^54^. In our model, AWA activity is proportional to the difference between the instantaneous local odor concentration (*C*) and a stored threshold (*T*). The threshold itself relaxes toward the local concentration at a rate proportional to *C* − *T*, introducing a single free parameter, *τ*, that sets the relaxation timescale. Fitted to AWA activity, this Adaptive Dynamical Threshold of Sensitivity (ADTS) model captured the dynamics of the inferred threshold *T* in crawling animals (Fig. 6C,E; Fig. S9A,C). When we applied the ADTS model to AWC dynamics, we found that AWC activity was negatively correlated with *C* − *T*, consistent with AWC functioning as an “off”-type sensory neuron activated by odor removal and suppressed by odor addition (Fig. 6D,E; Fig. S9B,C).

Although early ablation studies showed that DA chemotaxis is more sensitive to loss of AWA, we observed somewhat stronger correlations with AWA activity during IAA chemotaxis than during DA chemotaxis (Fig. 6F) ^51^. This may be due to the higher concentration used in IAA experiments.

### Functional mapping of an olfactory navigation circuit

To define the sensorimotor circuit linking AWA activity to navigational behavior, we identified neurons whose cross-correlation with AWA differed significantly across crawling conditions (Fig. 7A).

**Figure 7.**
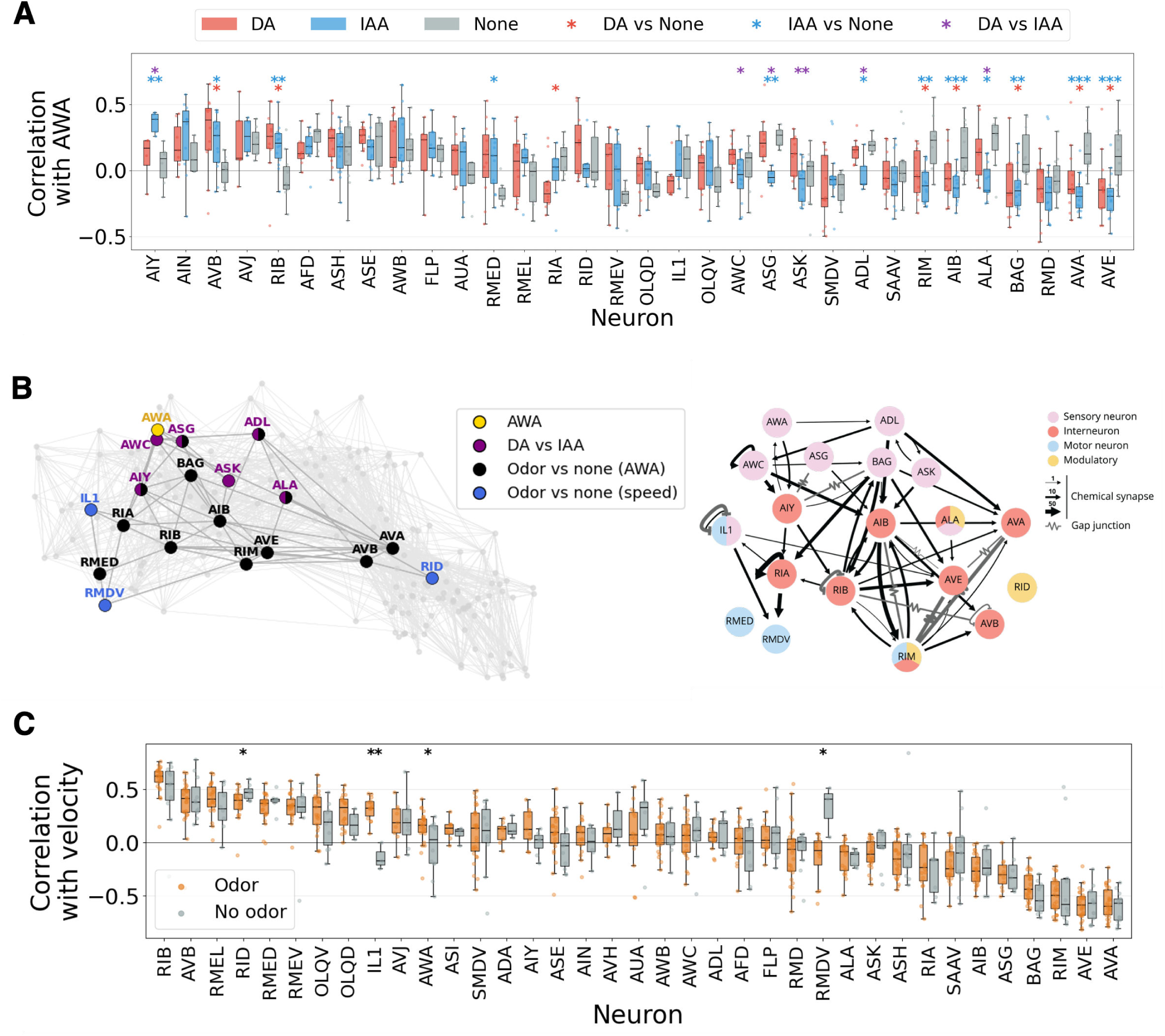
(**A**) Distribution of pairwise correlation between AWA and every other neuron in worms crawling in diacetyl gradients (12 animals), isoamyl alcohol gradients (16 animals), or isotropic environments (13 animals). Only neurons present in at least 3 datasets per stimulus are shown. Significance markings show results of two-sided Mann-Whitney U tests on the per-worm correlation values. Significance levels: *p<0.05, **p<0.01, ***p<0.001. (**B**) Wiring diagram of the neurons that show significant changes in correlation with AWA between different odors or between odor gradient and no-odor conditions, as well as neurons that show significant changes in correlation with velocity in odor vs. no-odor conditions. (Left) Neurons highlighted on the signal flow view of the connectome from Varshney et al. ^61^. (Right) Strength of connection diagrams generated at NemaNode.org, using the union of the two adult datasets from Witvliet et al. ^20^. (**C**) Distribution of pairwise correlation between velocity and every neuron in worms crawling in odor gradients (28 animals) or isotropic environments (13 animals). Only neurons present in at least 3 datasets per stimulus are shown. Significance markings show results of two-sided Mann-Whitney U tests on the per-worm correlation values. Significance levels: *p<0.05, **p<0.01, ***p<0.001.

Neurons that showed the largest differences in AWA correlation between DA and IAA navigation were predominantly other sensory neurons, including AWC, ASG, ASK, and ADL (Fig. 7A,B). This pattern suggests that odor identity is represented not by AWA alone, but by combinatorial activity distributed across a broader chemosensory ensemble ^23^. Such a coding scheme may be especially important in natural environments containing multiple odor cues, whereas our experiments quantified navigation in pure gradients of single odorants.

In contrast, differences in AWA correlations between olfactory gradients and isotropic environments revealed a compact sensorimotor pathway for navigation itself. This pathway extends from first-layer interneurons (AIB and AIY), through second-layer interneurons (RIA, RIB, and RIM), to premotor neurons for forward and backward locomotion (AVB and AVA) (Fig. 7A,B).

Thus, while differences between chemical gradients highlighted sensory-level divergence, differences between gradient navigation and isotropic exploration identified interneurons and premotor neurons that convert olfactory signals into locomotor output. The connectome supports this functional mapping ^20,55^. Many of these neurons were previously identified in systematic ablation studies, guided by the connectome, as components of the circuit that transforms olfactory sensory input into behavioral responses ^13,14^.

We expanded this functional mapping by identifying all neurons whose correlation with velocity differed between olfactory navigation and isotropic exploration (Fig. 7C). These neurons include IL1, which has an established mechanosensory role ^56,57^. IL1 activity was negatively correlated with forward speed, suggesting that olfactory navigation integrates odor information with mechanosensory feedback generated as the animal moves through its physical environment. IL1 and another identified neuron, RMD, also regulate spontaneous foraging head movements together with OLQ ^43,56,58^. The identified neurons also included RID, a neuroendocrine cell that promotes forward runs ^59^.

Together, these analyses define a compact functional pathway linking olfactory sensory activity to locomotor state during navigation.

### The activity of a first-layer olfactory interneuron depends on behavioral context

AIB, a first-layer interneuron, contributes to AWA-mediated olfactory chemotaxis through its joint representation of sensory input and locomotory state ^16,18^. AIB is thought to receive a corollary discharge associated with backward locomotion that reinforces sensory-evoked drive toward reversal. Accordingly, in immobilized animals, AIB activity is better predicted by models that combine incoming sensory signals, such as AWA activity, with a signal related to backward state, such as AVA activity.

To better understand how AIB contributes to olfactory chemotaxis, we compared its correlations with other neurons in animals navigating odor gradients and in isotropic environments (Fig. S10A) ^16,18^. AIB was strongly correlated with backward movement in both olfactory navigation and isotropic exploration (Fig. 7C).

However, several neurons predicted from the connectome to be synaptically connected to AIB showed context-dependent changes in their functional relationship with AIB. These included AWA and AIY, which became more negatively correlated with AIB during olfactory navigation, and SAA, a turning-associated head interneuron/motor neuron, which became more positively correlated with AIB during olfactory navigation (Fig. S10A) ^60^. By contrast, many other neurons, whether synaptically connected to AIB or not, showed little or no change in their functional correlation with AIB between olfactory navigation and isotropic exploration.

To further examine the joint encoding of sensory input and motor state by AIB, we compared immobilized worms responding to DA pulses with crawling worms navigating odor gradients and isotropic environments (Fig. S10B–G). Consistent with previous reports ^16,18^, AIB activity in immobilized animals was marginally better predicted by a joint encoding model that combined sensory input, represented by either AWA or AWC activity, with motor-state input, represented by AVA activity (Fig. S10F).

In immobilized worms, AIB activity was also well predicted by models using only anatomically defined presynaptic inputs, which include both olfactory sensory neurons and interneurons correlated with motor state, as well as by models incorporating all head neurons, which could capture indirect synaptic and nonsynaptic influences, including signals associated with AVA (Fig. S10F,G).

In crawling animals, by contrast, AIB activity during olfactory navigation was better predicted by AVA activity alone than in isotropic environments. Adding other neurons to the joint linear encoding model produced little or no improvement, suggesting that the determinants of AIB activity in crawling animals are either more distributed or more nonlinear (Fig. S10E,C).

Taken together, these results suggest that AIB undergoes a context-dependent reweighting of its relationship to sensory input, synaptic input, and motor state between immobilized and crawling conditions, and between olfactory navigation and isotropic exploration.

## Discussion

Animals navigate spatial environments by orienting self-motion toward distant goals. Because motor commands act through the biomechanics of the body and the surrounding environment, the relationship between neural commands and movement is inherently context dependent. Understanding how decision-making circuits convert sensory signals into navigational behavior therefore requires accounting not only for goal-related sensory input, but also for how the physical environment shapes the movements produced by motor commands. Those movements, in turn, generate sensory feedback that informs the animal about its surroundings and its progress toward distant goals.

Here, we examined brain-wide representations of olfactory sensorimotor responses in *C. elegans* evoked by diacetyl and isoamyl alcohol, two potent, naturally occurring attractive odorants sensed primarily by AWA and AWC olfactory neurons ^21,51^. As the worm moves through a spatial odor landscape, its movements sweep the sensory endings at its nose through the gradient, converting spatial differences in odor concentration into temporal sensory input.

Although many experimental setups have been developed to study olfactory navigation in *C. elegans*, physiological access to the underlying sensorimotor circuitry has often relied on constrained or immobilized preparations with externally imposed sensory input ^22,38,62–64^. We combined advances in multineuronal tracking, volumetric imaging, and neuronal identification to compare the activity of corresponding neurons across animals navigating odor gradients, exploring isotropic environments, and receiving odor pulses while being immobilized ^9,12,24,26,27,65,66^. These comparisons revealed a compact functional circuit linking olfactory sensory activity to navigational movement, with relationships among sensory neurons distinguishing odor identity and relationships with interneurons and premotor neurons distinguishing chemotaxis from isotropic exploration. They further showed how context-dependent neuronal interactions organize the olfactory circuit as animals respond to sensory input and move through their environment.

### Adaptive sensory coding

*C. elegans* uses an ensemble of chemosensory neurons to detect the concentration and identity of diverse odorant molecules ^51^. Within this ensemble, AWA and AWC are among the most broadly tuned olfactory neurons and are particularly important for chemotaxis to diacetyl and isoamyl alcohol ^23,67^. Previous work showed that AWC adapts to successive changes in background concentration by shifting its sensitivity threshold, thereby preserving responsiveness during long-range navigation ^54^. We captured this idea in an adaptive dynamical threshold (ADTS) model that describes AWA and AWC activity during chemotaxis to diacetyl and isoamyl alcohol. Although AWA responds to increasing odor concentration and AWC to decreasing concentration, the ADTS model captures their dynamics with opposite signs, suggesting a shared computational description of adaptation despite their opposing response polarities.

### A minimal navigational circuit

Brain-wide imaging reveals a compact sensorimotor circuit linking olfactory receptor neurons, interneurons, and head motor neurons, with distinct activity during olfactory navigation and isotropic exploration. This circuit extends from chemosensory neurons such as AWA and AWC, whose activity depends on odor identity, to head motor neurons such as RME and RMD. Differences in activity between odor gradients and isotropic environments also identify interneurons (AIB, AIY, RIA, RIB, RIM, AVA, AVB) that participate in the sensorimotor transformations regulating forward and backward movements.

These relationships begin to define a functional map of the core pathways underlying olfactory navigation. Many components have been characterized through targeted experiments, including AVA and AVB in backward and forward locomotion ^32,42^, RME and RMD in head movements associated with forward locomotion and foraging ^40,43,56,58^, and RID in sustaining the forward motor state ^59^. Brain-wide recording reveals how these many components are coordinated within a common dynamical framework during navigation.

### Behavioral context reshapes circuit function

Motor-related feedback influences first-layer olfactory interneurons, including AIY and AIB, that integrate incoming sensory information ^16,18,19^. In immobilized animals, AIB jointly represents olfactory input and behavioral state, with sensory responses gated by reversal state. AIB also contributes to reversal generation, placing it within a recurrent sensorimotor loop ^16,18^. Consistent with this view, we find that AIB is strongly recruited during freely moving navigation, where its activity becomes tightly coupled to backward locomotion in spatial odor gradients. The source of this motor-related signal remains unresolved, but it may arise through direct presynaptic input, recurrent signaling, electrical or metabolic coupling, or neuromodulation.

Brain-wide imaging also identifies sensory pathways that report environmental context. OLQ has been implicated in mechanosensation and in the rapid nose movements that accompany exploratory foraging in freely moving animals ^43,68^. Under immobilization, however, OLQ becomes strongly associated with fictive backward movement. Thus, the OLQ pathway may contribute to the abnormally long backward states observed in restrained worms.

The aerotaxis circuit shows a similar reconfiguration, in which the activity of the carbon dioxide-sensing BAG neurons tracked backward movement in navigating and exploring animals but was recruited by pulsed odor delivery when immobilized. When animals crawl beneath a coverslip, as required by this and other functional imaging setups, self-generated carbon dioxide can accumulate around the body, and reversals can carry the animal backward into a recently generated plume to activate BAG ^48^. Measurements in immobilized worms suggested that BAG can also participate in olfactory processing, effectively acting in an interneuron-like role within the olfactory circuit ^49,50^.

### Limitations and future directions

The circuit we identified likely represents a minimal map of the pathways that shape navigational behavior. Beyond the core circuitry linking olfactory input to forward and backward movement ^13,14^, our functional mapping includes neurons with mechanosensory roles, such as IL1^56,57^. Other mechanosensory neurons, including touch receptor neurons with cell bodies outside the head, were beyond our imaging coverage. Improved signal-to-noise ratios and broader volumetric coverage should extend brain-wide analysis to a wider range of sensory pathways, environmental contexts, and sensorimotor behaviors.

Improved methods for long-term brain-wide recording in individual animals will also be critical for characterizing persistent differences in neural dynamics among animals, even within isogenic populations. Such variability can arise from neuromodulation across development, which produces persistent individual differences in adult exploratory behavior ^69^, and from variation in circuit architecture itself, as reconstructions of multiple isogenic animals across development reveal widespread individual differences in synaptic wiring ^20^. Reliable brain-wide measurements in single animals, combined with quantitative analyses of behavior, gene expression, and connectivity, should help determine how these distinct sources of individuality are reflected in neural dynamics.

Another limitation of this study is that our measurements are confined to nuclear calcium dynamics. Our imaging approach uses nuclear-localized calcium indicators to isolate signals from individual neurons ^11,12^. While this strategy enables reliable cell identification and signal separation, it reduces sensitivity to intracellular dynamics elsewhere in the neuron. In *C. elegans*, calcium activity in certain neurites is only weakly correlated with somatic activity ^17^. Nuclear calcium measurements may therefore miss additional context-dependent dynamics within individual neurons.

### Conclusions

Effective navigation requires integrating information about the external environment with the animal’s own movements. In visually guided animals such as insects and vertebrates, neural representations of position and heading—including place cells, grid cells, and compass systems—relate self-motion to spatial goals ^70,71^. In non-visual animals such as *C. elegans*, multiple sensory modalities provide information about the environment and the animal’s movements within it. Mechanosensory and proprioceptive inputs report the mechanical structure of the environment, gas-sensing inputs report enclosure and movement-related context, and olfactory inputs coupled to self-motion report position relative to odor gradients and odor-emitting objects. Our findings suggest that brain-wide dynamics collectively represent the animal’s sensory environment and behavioral context, providing a distributed internal model that informs goal-directed navigation.

## Methods

Experiments were done using young adult worms reared at 23°C on NGM agar plates seeded with OP50 *E. coli*. We ensured worms were well-fed for at least two generations before experiments.

### Worm strains

#### Freely moving experiments

For brain-wide recordings, we used ADS1046, with genotype *lite-1(ce314)* X; *hpIs675* [P*rgef1*::GCaMP6s::3xNLS::mNeptune + *lin-15(+)*] *lin-15(n765)* X; *csbIs27* [P*ift-20*::BFP + P*eat-4*::cyOFP1 + P*unc-17*::mScarlet + P*acr-5*::BFP]. ADS1046 combines pan-neuronal calcium imaging with sparse multicolor markers for neuronal identification, following the strategy described by Yemini et al. ^12^. Because the positions and molecular features of *C. elegans* neurons are stereotyped, sparse fluorophore expression labels reproducible neuronal subsets across animals, allowing individual neurons to be distinguished.

The pan-neuronal imaging strain ZM9624 (*lin-15(n765)* X; *hpIs675* X) was crossed with the light-insensitive mutant MT21793 (*lite-1(ce314) gur-3(ok2245)* X). This cross produced ZM11178. Genotyping confirmed the *lite-1* mutation in ZM11178. We did not recover the *gur-3* mutation in this strain, likely because of its close proximity to the *hpIs675* transgene. We then crossed ZM11178 with the sparse-marker strain JAC766 carrying *csbIs27* [P*ift-20*::BFP + P*eat-4*::cyOFP1 + P*unc-17*::mScarlet + P*acr-5*::BFP]. The *csbIs27* array expresses three spectrally separable fluorophores—BFP, cyOFP1, and mScarlet—which we distinguished using combinations of excitation lasers and emission filters (see Image acquisition in freely moving animals). This cross produced ADS1046. We confirmed the presence of *hpIs675* and *csbIs27* by fluorescence, and the *lite-1* mutation by genotyping. We used the following PCR and sequencing primers:

- F_lite-1_ce314: TAAGCTGAACAGCAAGAGGAC
- R_lite-1_ce314: CTTTCAGGTGTTCTGAATGCG
- F_seq_lite-1_ce314: GCTTGATCGAACGACGGATG
- R_seq_lite-1_ce314: AAGTTGCCGGTTGCCTTAG

#### Immobilized experiments and consistency across NeuroPAL and ADS1046 strains

For immobilized microfluidic experiments, we primarily used OH16230 (*otIs672*[NeuroPAL; *rab-3*::NLS::GCaMP6s + *arrd-4*::NLS::GCaMP6s]), which expresses pan-neuronal GCaMP6s together with the NeuroPAL transgene for neuronal identification (see Yemini et al. ^12^ for the complete genotype). We developed ADS1046 for freely moving experiments because it supports robust locomotion and chemotaxis, whereas OH16230 moves more slowly and exhibits impaired chemotactic behavior ^12^. Conversely, the greater color diversity of NeuroPAL facilitates neuronal identification in microfluidic devices, where compression can distort head geometry and reduce the reliability of positional cues.

To assess whether strain choice affected the measured neural dynamics, we also recorded brain-wide activity from immobilized ADS1046 animals (*n* = 4) using the same experimental protocol and compared these recordings with those from immobilized OH16230 animals (*n* = 8). The two strains exhibited similar activity patterns in neurons associated with fictive forward and backward states and similar context-dependent relationships between neuronal activity and motor state (Fig. S4). Immobilized ADS1046 animals also reproduced the odor-evoked AWA and AWC responses measured in OH16230 animals (Fig. S4D; Fig. 6A). These comparisons support the consistency of the principal neural dynamics and neuronal identifications across the two strains (see Neuron ID annotation for ADS1046).

### Creating attractant gradients for navigation experiments

We studied navigation towards two attractive odorants: diacetyl (DA) and isoamyl alcohol (IAA). Radial spatial gradients of odorant were established following a procedure similar to the methods described in Pierce-Shimomura et al. ^28^. We placed a 5 *µ*L droplet of diluted odorant (10% v/v for IAA and 1% v/v for DA) at the center of a standard 10 cm Nematode Growth Medium (NGM) agar plate. We let the odorant diffuse through agar in the uncovered plate for 2–4 hours.

Diffusion was modeled using the solution to the diffusion equation for a point source in a cylindrical volume with the depth of the agar plate: ^28^

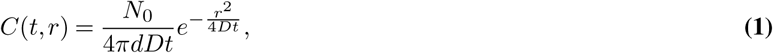

where *N*_0_ is the amount of attractant entering the agar, *d* is the agar depth, *r* is the distance from the odor source, and *t* is the time since placement of the odor droplet. Because the odorants are volatile, the fraction of the applied odorant that entered the agar was unknown, precluding an estimate of absolute concentration. We therefore used only the relative spatiotemporal concentration profile predicted by the model. The unknown multiplicative prefactor *N*_0_*/*(4*πdD*) was absorbed into an arbitrary normalization, and modeled concentrations were expressed as dimensionless relative values. Based on molecular size, we estimated the diffusion coefficient for both diacetyl and isoamyl alcohol to lie between those of biotin and fluorescein ^28^, and used *D* = 10^−3^ mm^2^ s^−1^. Varying *D* by an order of magnitude in either direction did not qualitatively alter the results.

For each recording, we placed three worms on a prepared agar plate 2–4.5 hours after odor placement. We selected worms directly from the plates where they were grown, choosing those outside the food lawn to reduce the chance of OP50 presence on the experimental odor plates. Worms were placed in a 10 *µ*L NGM droplet 1–1.5 cm away from the odor source. A No. 1 25×25 mm coverslip was placed on top, covering the imaging area. Since coverslip placement might move the worms away from the origin position or obstruct their movement, placing three worms increased the likelihood of having a viable worm for imaging on each plate.

### Microfluidics experimental design

We adapted the microfluidic design of Lin et al. ^23^ to deliver controlled odor pulses directly to the worm’s nose, with rapid switching between odor and buffer streams under controlled pressure ^22,72^. We modified the original device by reducing the width of the worm channel to 55 *µ*m.

Worms were washed in M9 buffer containing 3 mM tetramisole to induce paralysis and then loaded into the microfluidic device using a syringe. During each run, animals received six 10-second odor pulses separated by 30-second buffer periods, with an additional 30 seconds of buffer before the first pulse and after the final pulse, for a total run time of 4.5 minutes. For all experiments, we used diacetyl diluted to 0.001% v/v in CTX buffer (5 mM KH_2_PO_4_/K_2_HPO_4_ at pH 6, 1 mM CaCl_2_, 1 mM MgSO_4_, and 50 mM NaCl, adjusted to 350 mOsm/L with sorbitol).

### Image acquisition in freely moving animals

We acquired two types of volumetric images. We first acquired recordings of neuron activity while animals were navigating odor gradients. After each recording, we paralyzed the worms and obtained a static volume using multiple filter and laser combinations to separately record expression of every fluorophore, thus producing an ID image for neuron identification.

#### Brain-wide recordings of neuronal activity

Following Susoy et al. ^25^ and Venkatachalam et al. ^24^, we used a custom spinning-disc confocal microscope to record GCaMP6s and mNeptune fluorescence simultaneously. Emitted fluorescence was separated using an mCherry/GFP dichroic cube and projected onto two Andor Zyla 4.2 sCMOS cameras. The red channel was filtered using a Semrock BrightLine 617/73 nm filter, and the green channel using a Semrock BrightLine quad-band 440/521/607/700 nm filter. Volumetric stacks comprising 20 planes of 256 × 512 pixels were acquired at 5 volumes per second with an exposure time of 5 ms per plane. Successive planes were spaced 2 *µ*m apart. Imaging was performed with a 40×, 1.30-NA Nikon Plan Fluor oil-immersion objective. We manually tracked each animal to keep its head within the field of view. The imaging region was continuously illuminated with 488and 561-nm lasers. The 488-nm laser power was set to the highest level that did not elicit light-avoidance behavior.

Each animal was recorded for 5–30 minutes. Recordings were stopped when the animal reached the odor source, left the coverslip area, or showed signs of photobleaching. We recorded the position of the odor source relative to the animal and the elapsed time between odor placement and completion of the experiment.

Control experiments were performed identically on plates without added odorant.

#### ID image acquisition

After each navigation experiment, we immobilized the recorded animal by submerging it in a 5 *µ*L droplet of 25 mM tetramisole. After allowing up to 1 hour for complete immobilization, we acquired the ID image.

To separate the fluorescent markers expressed in the ADS1046 strain (GCaMP6s, mNeptune, mScarlet, cyOFP1, BFP), we used a combination of four laser frequencies and five emission filters. Table 1 shows the sequence of laser wavelengths and filters used for ID image acquisition.

**Table 1.** Sequence of laser and filter combinations used for ID image acquisition, in the same order used for acquisition. The ‘Red channel’ and ‘Green channel’ settings are designed to match those used for activity recording as closely as possible, and are included to permit matching annotations between activity and ID images. All filters are Semrock BrightLine.

|  | **Laser** | **Filter** | **Target fluorophore** |
|---|---|---|---|
| 1 | 640 nm | 731/137 nm | mNeptune |
| 2 | 561 nm | 625/90 nm | Red channel |
| 3 | 561 nm | 578/21 nm | mScarlet |
| 4 | 488 nm | 578/21 nm | cyOFP1 |
| 5 | 488 nm | Quad-band 440/521/607/700 nm | Green channel |
| 6 | 405 nm | 445/45 nm | BFP |

We recorded 41 animals using this setup: 12 in DA gradients, 16 in IAA gradients, and 13 in isotropic conditions. Complete activity heatmaps for all annotated neurons, together with concurrent sensory and behavioral measurements, are shown for each freely moving recording in Fig. S12, with one recording shown in Fig. 1E.

### Image acquisition in immobilized animals

Imaging was performed using a single-photon inverted spinning-disc confocal microscope. For each animal, we first acquired a static ID image for subsequent neuron identification, followed by 1–4 neural activity recordings during presentation of the stimulus sequence. Both image types were acquired with a 60×, 1.2-NA water-immersion objective. The recorded volume measured 112 *µ*m × 56 *µ*m × 30 *µ*m and encompassed the head region containing most neuronal nuclei. We acquired 21 planes spaced 1.5 *µ*m apart using a piezoelectric device.

For the ID image, we acquired volumes for each of the fluorophores expressed by OH16230 separately. The spatial resolution was 0.5 *µ*m by 0.5 *µ*m by 1.5 *µ*m per voxel. OH16230 contains three sparse markers—BFP, cyOFP1, and mNeptune—as well as pan-neuronal RFP and GCaMP6s. Table 2 shows the list of filters and lasers used to acquire each volume. Some laser/filter pairings differ between OH16230 and ADS1046 (see Image acquisition in freely moving animals) even when targeting the same fluorophore, because of different cross-talk constraints in each strain (e.g., mScarlet must be distinguished from cyOFP1 in ADS1046 but not in OH16230). When taking ID images of ADS1046 in the inverted microscope for ID atlas construction (see Neuron ID annotation for ADS1046), we used the list of filters and lasers in Table 3. Although some of the laser/filter pairings are not the same as in Table 1, the settings for the three sparse markers used for ID are matched.

**Table 2.** List of lasers and filters used for NeuroPAL ID image acquisition, shown in order of acquisition. Filters are Semrock BrightLine.

|  | Laser | Filter | Target fluorophore |
|---|---|---|---|
| 1 | 405 nm | 445/45 nm | BFP |
| 2 | 561 nm | R488/568 nm | RFP |
| 3 | 488 nm | R488/568 nm | GCaMP6s |
| 4 | 488 nm | 525/40 nm | cyOFP1 |
| 5 | 561 nm | 731/137 nm | mNeptune |

**Table 3.** List of lasers and filters used for ADS1046 ID image acquisition in microfluidics experiments using an inverted microscope, shown in order of acquisition. The ‘Red channel’ setting captures pan-neuronal red expression but is not used in practice. Filters are Semrock BrightLine.

|  | **Laser** | **Filter** | **Target fluorophore** |
|---|---|---|---|
| 1 | 488 nm | R488/568 nm | GCaMP6s |
| 2 | 561 nm | R488/568 nm | Red channel |
| 3 | 405 nm | 445/45 nm | BFP |
| 4 | 488 nm | 578/21 nm | cyOFP1 |
| 5 | 561 nm | 578/21 nm | mScarlet |
| 6 | 640 nm | 731/137 nm | mNeptune |

Neuron activity was acquired using a 488 nm laser at 2.51 volumes per second, with a BrightLine 525/40 nm emission filter. The spatial resolution was 1 *µ*m by 1 *µ*m by 1.5 *µ*m per voxel. Before the first run and between runs, the worm was allowed to rest for 1 minute without illumination.

We recorded eight animals for brain-wide activity analysis, with three runs per animal. We additionally recorded 63 animals for AVA-only analysis, with 1–4 runs per animal and 153 runs in total. Complete activity heatmaps for all annotated neurons, together with odor stimuli and fictive-state annotations, are shown for each brain-wide immobilized recording in Fig. S13, with one recording shown in Fig. 1F.

### Computational methods for neuronal activity extraction

To track neuronal positions despite deformation of the *C. elegans* head during undulatory movement, we adapted the point-tracking method developed by Park et al. ^26^.

#### Preprocessing

We first applied a difference-of-Gaussians (DoG) filter (*σ*_1_ = 1 pixel, *σ*_2_ = 5 pixels, 21-pixel kernel) to each *z* slice independently. Affine transformations to align the red and green channels were determined by manually annotating matching points in at least 3 frames. A custom-trained U-Net was used to segment the images and label each point as part of the front or back head ganglia. Using all the segmented points, we computed the center of mass to center the image. The head rotation was corrected using the eigenvector of the second-order image moment associated with the largest eigenvalue, which corresponds to the long axis of the head. The head orientation was determined using the front and back ganglia mask. If the mask did not distinguish two separate ganglia, the orientation was determined from the sign of the third moment along the rotation axis (computed after the initial rotation), which captures the skewness of the mass distribution. We manually corrected any frames that were oriented backward by an additional 180°. Finally, we cropped the images to the neuron area to reduce the memory required to train the neural network.

#### Neuron tracking

To generate the ground truth for the annotation neural network, we manually annotated 10 frames in each recording, labeling each neuron center with a number such that the same number was given to the same neuron in each frame. We labeled only the left or right half of the worm (whichever was closer to the objective in each recording), obtaining 80–137 neurons (median 92) across 41 recordings. The ground-truth frames were augmented using targeted augmentation ^26^ and used to train a convolutional neural network that predicted neuronal positions in the remaining frames. We manually reviewed the predictions. When additional correction was needed, we annotated one or two more frames and retrained the network.

#### Signal extraction

We extracted signals by integrating a Gaussian centered on each predicted point (standard deviation 1.5 pixels in *x* and *y*, 0.4 slices in *z*; 7×7×1 voxel kernel), separately for the red and green channels. The signals were corrected for differences in laser intensity depending on the position in the field of view by using a measured intensity field from a uniform fluorescent slide. Each channel was separately debleached by dividing by a fitted decaying exponential function. Time points at which the signal deviated by more than 1.5 standard deviations from the distribution of neighboring values within 50 frames on either side were treated as outliers and removed. Missing values were filled by one-dimensional linear interpolation along the time axis. To correct for motion artifacts, we used a Bayesian probabilistic model to separate the shared red/green motion-related signal from the green-only calcium signal, using the method from Creamer et al. ^73^. We observed that the first principal component of neural activity across all neurons in each animal was a global component likely reflecting residual imaging artifacts, and removed it to obtain the final neuronal activity traces. Finally, activity was smoothed with a Savitzky–Golay filter (order 1, window 11 frames), and the first and last 100 timepoints of each recording were discarded to avoid preprocessing and start- or end-of-experiment edge effects. For visualization of single-neuron traces only, we applied a second pass of the same Savitzky–Golay filter (order 1, window 11 frames); this additional smoothing was used solely for plotting and not for any quantitative analysis.

We further restricted neurons to those where the green fluorescence was above background levels. We estimated the background fluorescence distribution in each recording by sampling pixels from 100 random frames. For each frame, we sampled 100 random pixels, excluding any that were within 10 pixels from a tracked neuron position. At each pixel, a Gaussian of the same size as that used for signal extraction was used to determine the background value. A neuron was considered *recorded* if the mean green channel signal for the neuron exceeded the background mean by one standard deviation of the background distribution.

### Neuron ID annotation for ADS1046

We assigned identities to 47–70 neurons per recording (median 64) across 41 experiments, all on the same side of the animal.

Beyond pan-neuronal GCaMP6s and mNeptune, ADS1046 expresses three sparse fluorophores (BFP, cyOFP1, mScarlet) driven by promoters that label distinct neuronal subsets. Their combined expression produces a stereotyped color pattern across animals, which we used together with neuron position to assign identities.

We built a neuron identification atlas in two stages. First, we matched neurons across animals: we found correspondences between different ID images based on neuron position and fluorescence levels, using characteristic activity patterns from the freely moving recordings (e.g., correlations with forward or backward motion) as an additional cue when needed. Once neurons could be matched across worms, we assigned each cross-animal cluster a known identity.

Identities were assigned using the published expression patterns of the four sparse promoters. *eat-4* marks glutamatergic and *unc-17* cholinergic neurons, ^74,75^ so we used the neurotransmitter maps defined in Serrano-Saiz et al. ^76^ and Pereira et al. ^77^ to identify these subsets. *ift-20* marks ciliated sensory neurons, ^78^ and *acr-5* marks nerve-ring neurons and a subset of head neurons (Table 4). ^79^

**Table 4.** List of neurons that express *ift-20* or *acr-5*. ^79^

| **Gene** | **Neurons** |
|---|---|
| *ift-20* | IL1, IL2, OLL, BAG, ASK, ADL, ASI, AFD, AWA, AWB, ASG, ADF, ASH, ASE, AWC, ASJ, FLP |
| *acr-5* | AVG, URA, URY, ALA, ventral cord neurons |

As an additional validation, we recorded ADS1046 in the microfluidic device under diacetyl pulses (see Microfluidics experimental design) and compared the resulting response patterns to those obtained from the NeuroPAL strain in matched experiments.

### Neuron activity extraction in immobilized animal recordings

#### Brain-wide activity experiments

We annotated neurons using the NeuroPAL CELL-ID software GUI (https://github.com/ `amin-nejat/CELL_ID`). ^12^ We analyzed brain-wide activity from eight animals, tracking 133–163 neurons per animal (median 146) and assigning identities to 84–114 neurons per animal (median 102) across both sides of the head.

For each animal, we transferred the annotated neuronal positions from the ID image to the first frame of each activity recording using an adaptation of NeuroPAL-traces (https://github.com/venkatachalamlab/NeuroPAL-traces) ^24^. We manually proofread these annotations using a custom GUI and then tracked neuronal positions through the remaining frames using ZephIR ^27^. We extracted neural activity by weighting the voxels surrounding each tracked position with a Gaussian having a standard deviation of 1.5 pixels in *x* and *y* and 0.4 slices in *z*We corrected each neural activity trace for photobleaching by fitting an exponential with an offset and dividing the raw trace by the fit. If the fit failed or did not describe a decay because it had a negative amplitude or a positive growth rate, we instead divided the trace by its mean. We then z-score normalized the resulting traces.

#### AVA-only experiments

We collected data for 63 worms, for a total of 153 runs. We used these data to determine forward and backward segment lengths and response rates in Fig. 3.

We annotated AVA neurons using the NeuroPAL CELL-ID software GUI (https://github.com/amin-nejat/ `CELL_ID`). ^12^ Neuron positions were tracked and activity was extracted using NeuroPAL-traces (https://github.com/ `venkatachalamlab/NeuroPAL-traces`). ^24^ We then excluded the first and last frames of each trace and flagged a frame as an outlier when its differences from the preceding and following frames had opposite signs and both magnitudes exceeded the mean plus 10 standard deviations of the activity values in that trace. To correct for bleaching, we fit a decaying exponential to a median-filtered version of each trace (25-frame window) and subtracted it from the raw trace when the fit was decaying. Finally, we normalized each trace by anchoring its 5th and 95th percentiles (computed after excluding the first 25 frames) to 0 and 1.

Fictive forward and backward transitions were inferred from AVA activity as described in Fictive behavior measurements.

### Behavior and concentration measurement extraction

To quantify behavior in freely moving recordings, we used neuronal positions in each frame. We defined the animal’s instantaneous position using RMDV, which was identified in every recording, and linearly interpolated its position across missing frames. We estimated the relative odor concentration experienced by the animal from its position and the elapsed time since odor placement using Equation 1.

Additionally, we computed velocity, angle of the head with respect to the body, and angle to odor (except in no-odor control worms):

- **Velocity.** We computed velocity from per-frame displacement, assigning a negative sign when the displacement vector formed an angle greater than 90^◦^ with the body axis (i.e., when the worm moved backward). The body axis was determined using the rotation of the frames during preprocessing (see Computational methods for neuronal activity extraction).
- **Head angle.** We computed head angle as the angle between the body axis and the vector connecting two anterior reference neurons. The posterior reference neuron was usually M3, and the anterior reference neuron was MCL, NSM, or I2, in that order of preference. When these neurons were not annotated, which occurred in four datasets, we manually selected annotated neurons as close as possible to the default reference neurons.
- **Angle to odor.** Angle to odor was computed by finding the angle between the worm’s absolute head angle (the sum of the head angle and the body axis) and the vector that points from the worm’s position to the odor source. An angle of 0^◦^ indicates the worm is headed directly towards the odor and an angle of ±180^◦^ indicates the worm is headed directly away from it.

For all behavioral and concentration time series, we excluded values that deviated by more than 1.5 standard deviations from the distribution of neighboring values within 50 frames on either side. We then linearly interpolated the missing values.

#### Pirouette and reversal identification

Using the sign of the worm velocity, we identified segments of each trajectory corresponding to forward and backward movement and manually proofread them by examining the trajectories. We classified each backward segment as either a single turn or part of a pirouette. Pirouettes are bouts of turns in which forward and backward segments have similar lengths and the animal remains within a localized area. We manually identified the end of a pirouette when the animal resumed a sustained forward run that was longer than the forward segments within the pirouette. See Fig. 2 for examples of single turns and pirouettes annotated on a trajectory.

### Fictive behavior measurements

To infer fictive behavior, we used the activity of the neuron AVA, which is associated with reversals. In freely moving worms, this neuron is active during backward motion and quiet during forward motion. We observed this in our freely moving recordings and it is well established in the literature. ^31,32,80,81^

We manually proofread AVA activity to ensure the recordings were high-quality enough to infer fictive behavior from them. When both AVAL and AVAR were available, we used their average to infer behavior. All recordings had at least one AVA neuron. We filtered the resulting activity trace using a Savitzky-Golay filter with window size 11 frames and computed its derivative. We automatically detected derivative peaks and troughs and manually proofread them to ensure quality. We omitted the first 5 seconds of the recording to avoid artifacts. A positive peak in the AVA activity derivative indicated a transition from fictive forward to fictive backward state, whereas a trough indicated the reverse transition. We defined the negative of AVA activity as fictive velocity.

### Data analysis

#### Sensory neuron activity model details

The ADTS model definition from Fig. 6B was discretized as

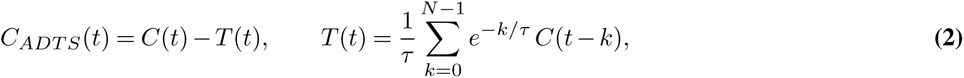

where we used *N* = 500 frames. For *t* ≥ *N* −1, an additional boundary correction *e*^−^*^N/τ^ C*(*t* −*N* +1) was added to *T* (*t*), account-ing for the contribution from times before the convolution window under the assumption that *C* was held constant before *t* − *N* + 1. For *t < N* − 1, where the full *N* -point window is not yet available, *T* (*t*) was computed as a weighted average 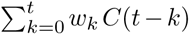 with weights *w_k_* ∝ *e*^−^*^k/τ^* normalized to sum to 1, plus an analogous boundary term *C*(0) *e*^−^*^t/τ^* to account for the assumed constant pre-recording concentration *C*(0). The first 350 frames were excluded from fitting and plotting to avoid this initialization regime. Before fitting, the concentration was 1-D-interpolated to remove any missing values. AWA activity was low-pass filtered with a 4th-order Butterworth filter at 0.07 Hz, applied prior to the Savitzky–Golay step described in Computational methods for neuronal activity extraction. We fit AWA activity *AWA*(*t*) = *a C_ADTS_* + *b*, where *a*, *b*, and *τ* are free parameters, independently for each worm. We determined the best *τ* for each odorant by averaging the fitted values for all datasets where the fit converged. The mean *τ* for diacetyl was *τ* = 29.6 seconds and for isoamyl alcohol it was *τ* = 34.5 seconds. All datasets were refit for *a* and *b* using the mean *τ* values for their corresponding odorant. The fit correlation values were computed using the AWA activity obtained with standard preprocessing.

#### AIB model detail

We compared linear models that predicted AIB activity from different sets of neurons. For freely moving recordings, we held out the middle quarter of each AIB activity trace as the test set. For immobilized recordings, we averaged AIBL and AIBR when both were available and held out the middle of the three runs as the test set.

For single-neuron (AVA) or neuron-pair (AVA + AWA or AVA + AWC) models, coefficients were estimated by ordinary least squares with an intercept (scikit-learn’s LinearRegression ^82^).

We also fit models using either all neurons presynaptic to AIB or all recorded neurons. Presynaptic inputs were defined as neurons with chemical synapses onto AIB in either of the two adult connectomes reported by Witvliet et al. ^20^. For these models, we used elastic-net regression with hyperparameters selected by cross-validation on the training set using scikit-learn’s ElasticNetCV ^82^.

The overall penalty strength *α* and the L1/L2 mixing ratio *ρ* were chosen by five-fold cross-validation on the training set, without access to the held-out test segment. We searched a 50×50 grid of (*ρ, α*) values, with *ρ* ranging from 0.001 to 1 and *α* logarithmically spaced between 10^−5^ and 10. For each combination we computed the mean five-fold cross-validated mean-squared error on the training data, selected the pair that minimized this error, and refit the model on the full training set using those hyperparameters.

We used the fitted coefficients to predict AIB activity in the held-out test segment. For cross-animal prediction, we retained only predictors identified in both the training and test animals and applied the corresponding coefficients from the trained model. Because potentially informative neurons absent from the test recording were omitted, this procedure may underestimate transfer performance. We quantified within-animal and cross-animal prediction performance using the Pearson correlation between measured and predicted AIB activity in the test set.

#### Cumulative variance explained

For each dataset separately, we performed PCA on the preprocessed, z-score-normalized neuronal activity traces using scikit-learn ^82^. For immobilized animals, we averaged bilateral neuron pairs before z-score normalization. We then calculated the fraction of variance explained by each principal component. We averaged these fractions across freely moving animals recorded in odor gradients, excluding isotropic controls, and separately across all brain-wide recordings from immobilized animals.

#### Measuring odor-evoked transition rates

To quantify odor-evoked fictive transitions, we required each animal to remain in the same fictive forward or backward state for the 5 seconds preceding stimulus onset or offset. Because changes in neural activity typically began approximately 3 seconds before the annotated state transition, we imposed an additional 3-second delay after stimulus onset or offset and required the initial state to persist throughout this delay. We then counted transitions occurring during the subsequent 5 seconds.

### Additional resources

Recorded brain volumes, neuronal identities, activity traces, and concurrent sensory and behavioral variables are available through an interactive web-based browser: https://helenacasademunt.github.io/worm-viewer/

## AUTHOR CONTRIBUTIONS

A.S. and M.Z. developed the strategy for brain-wide imaging. H.C. and A.S. designed experiments. H.C. did the experiments and data analysis. S.K. and W.F. helped with data analysis. W. H, J.L., H.Z., J.C., and M.Z. designed and built the transgenic *C. elegans* strains. C.P. wrote the software to extract neural traces. H.C. and A.S wrote the manuscript.

## ACKNOWLEDGEMENTS

We thank Albert Lin for help with microfluidics. We thank members of the Samuel and Zhen labs for useful conversations. Microfluidics devices were manufactured using the Soft Materials Cleanroom facility of the Harvard MRSEC (DMR-1420570).

## Supplement

**Figure S1.**
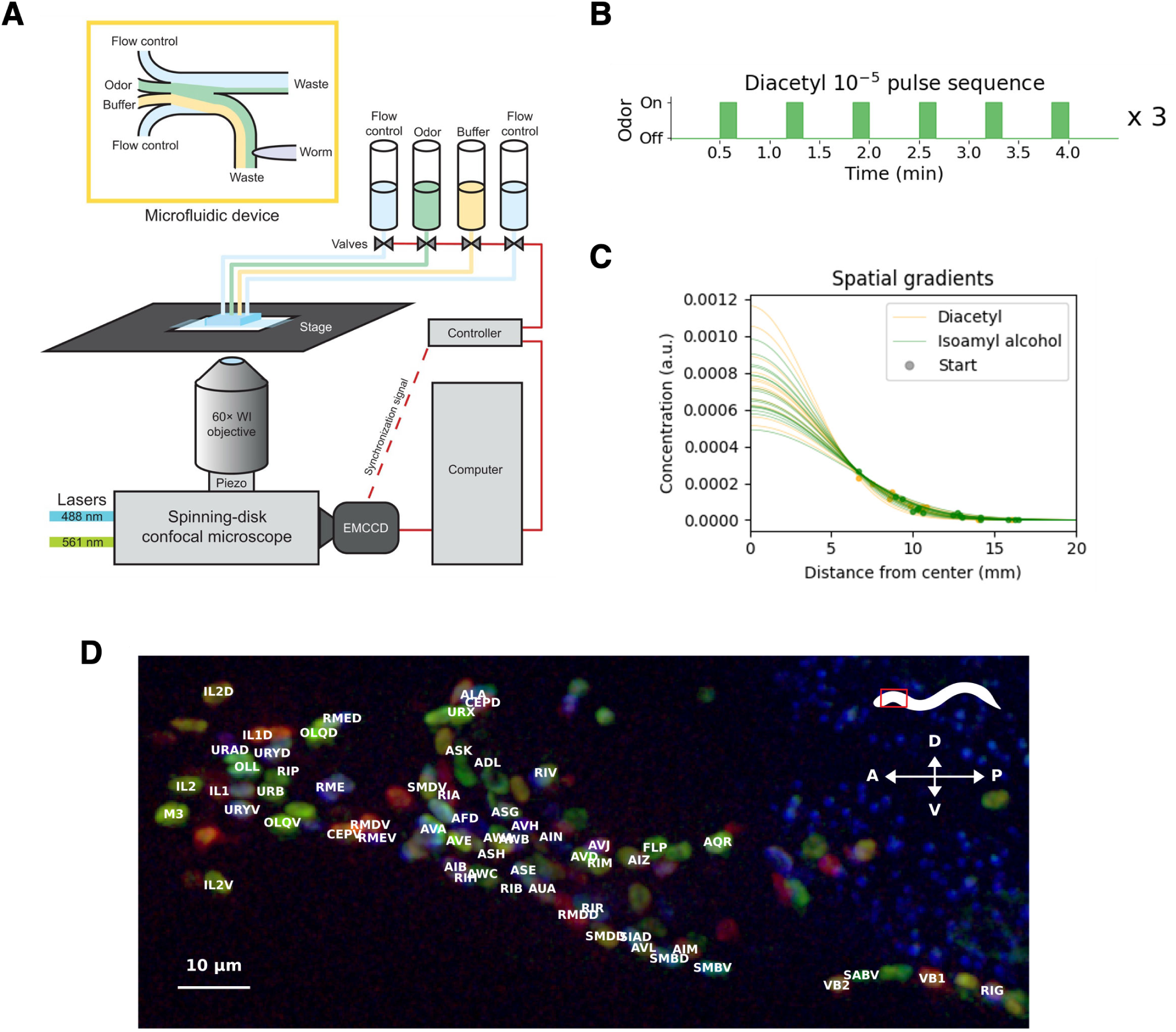
(**A**) Schematic of inverted microscope setup and chip used in microfluidics experiments. Figure adapted from Lin et al. ^23^. (**B**) Stimulus sequence for worms immobilized in microfluidic chips, consisting in 6 10-s diacetyl (10^−5^ dilution) separated by 30-s buffer intervals. The sequence is repeated for three runs per worm. (**C**) Concentration profiles for all worms in the same (arbitrary) concentration units. Markers indicate worm placement on the profile. (**D**) Maximum intensity projection image of the head of a NeuroPAL worm with neuron labels. Only labels for neurons on the right size are shown; left side is omitted for clarity. The image shows the NeuroPAL sparse markers only, omitting panneuronal markers. Image taken with a 60x water immersion objective. A: anterior; P: posterior; D: dorsal; V: ventral.

**Figure S2.**
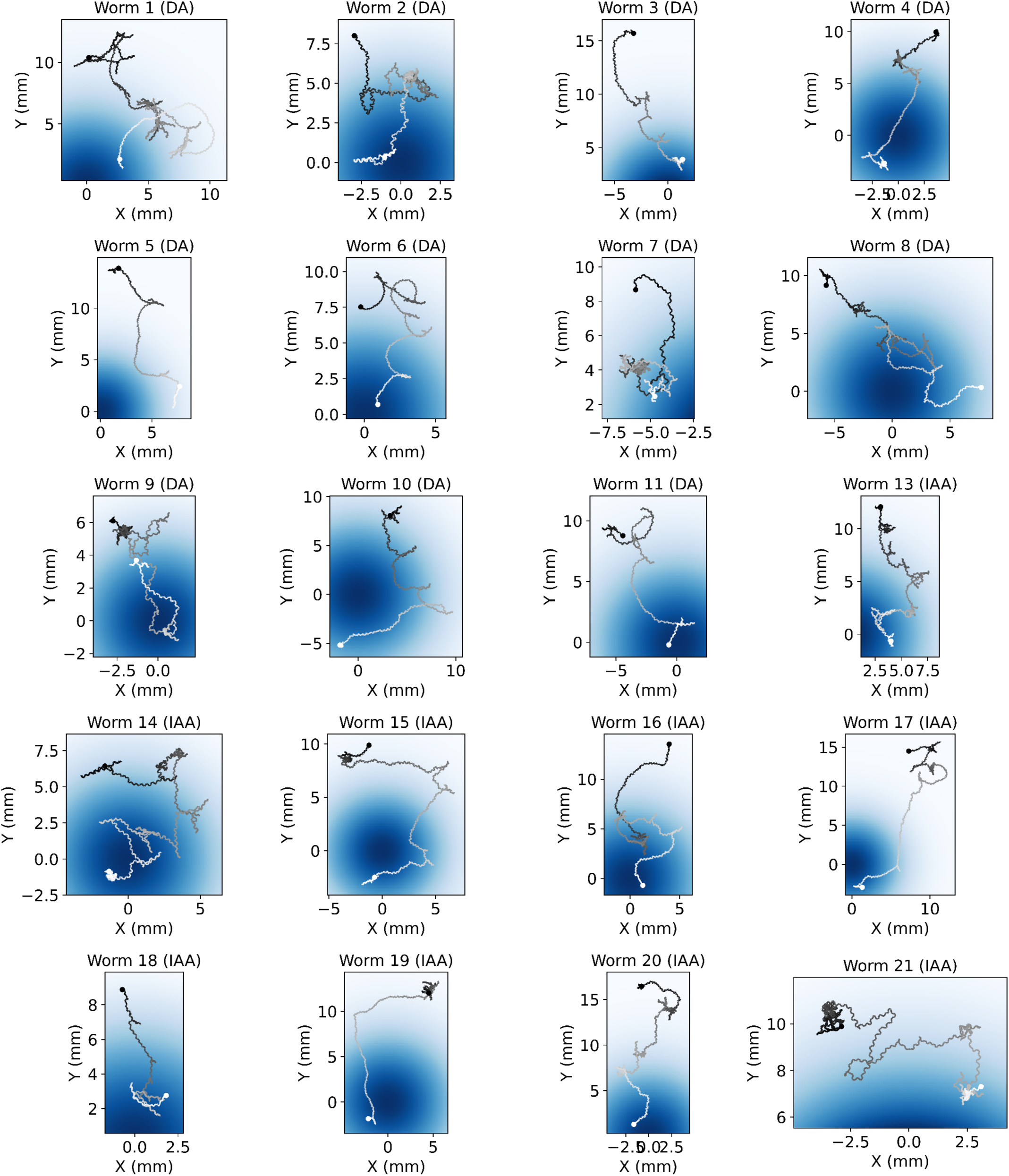
Individual trajectories

**Figure S3.**
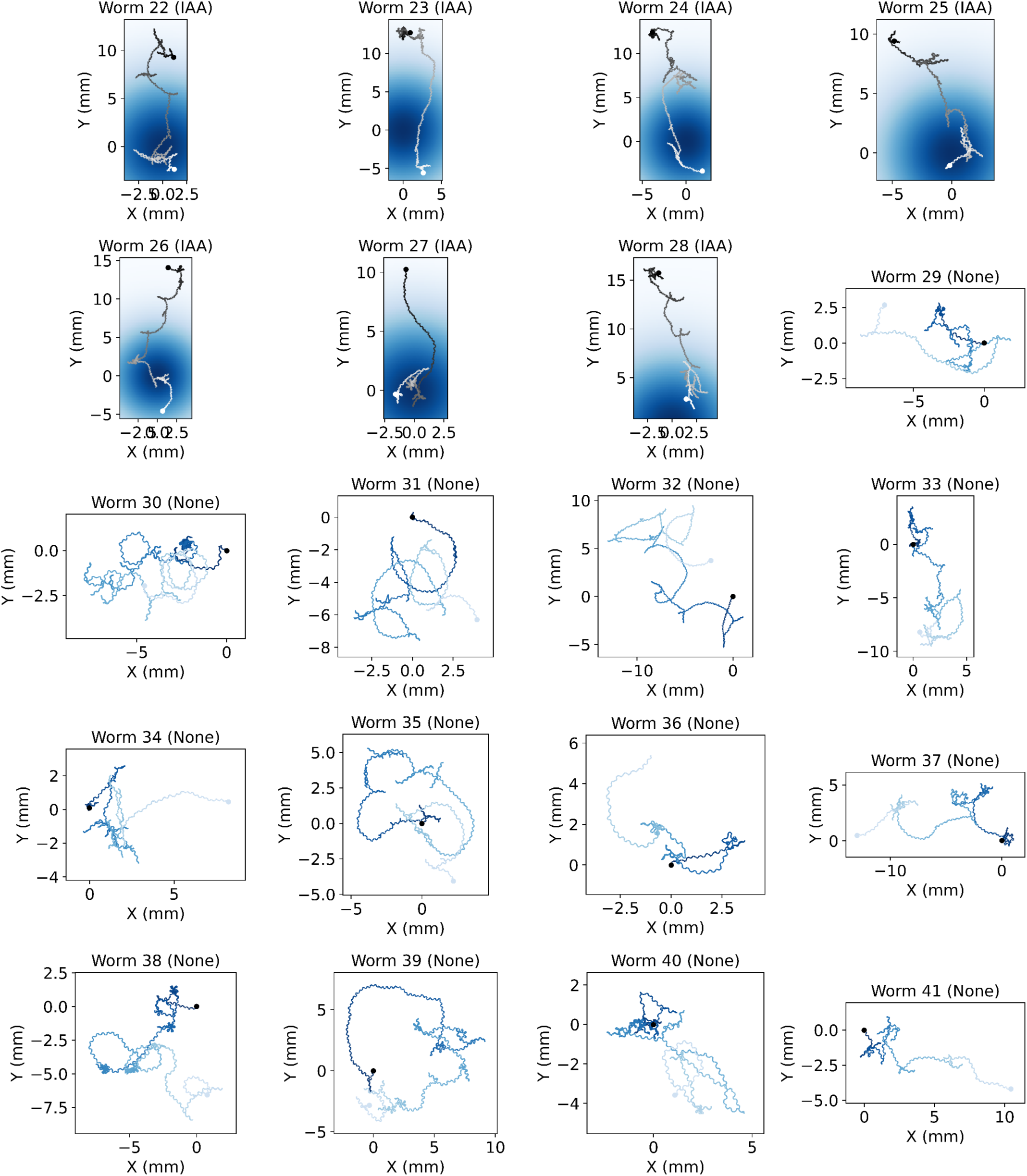
Individual trajectories

**Figure S4.**
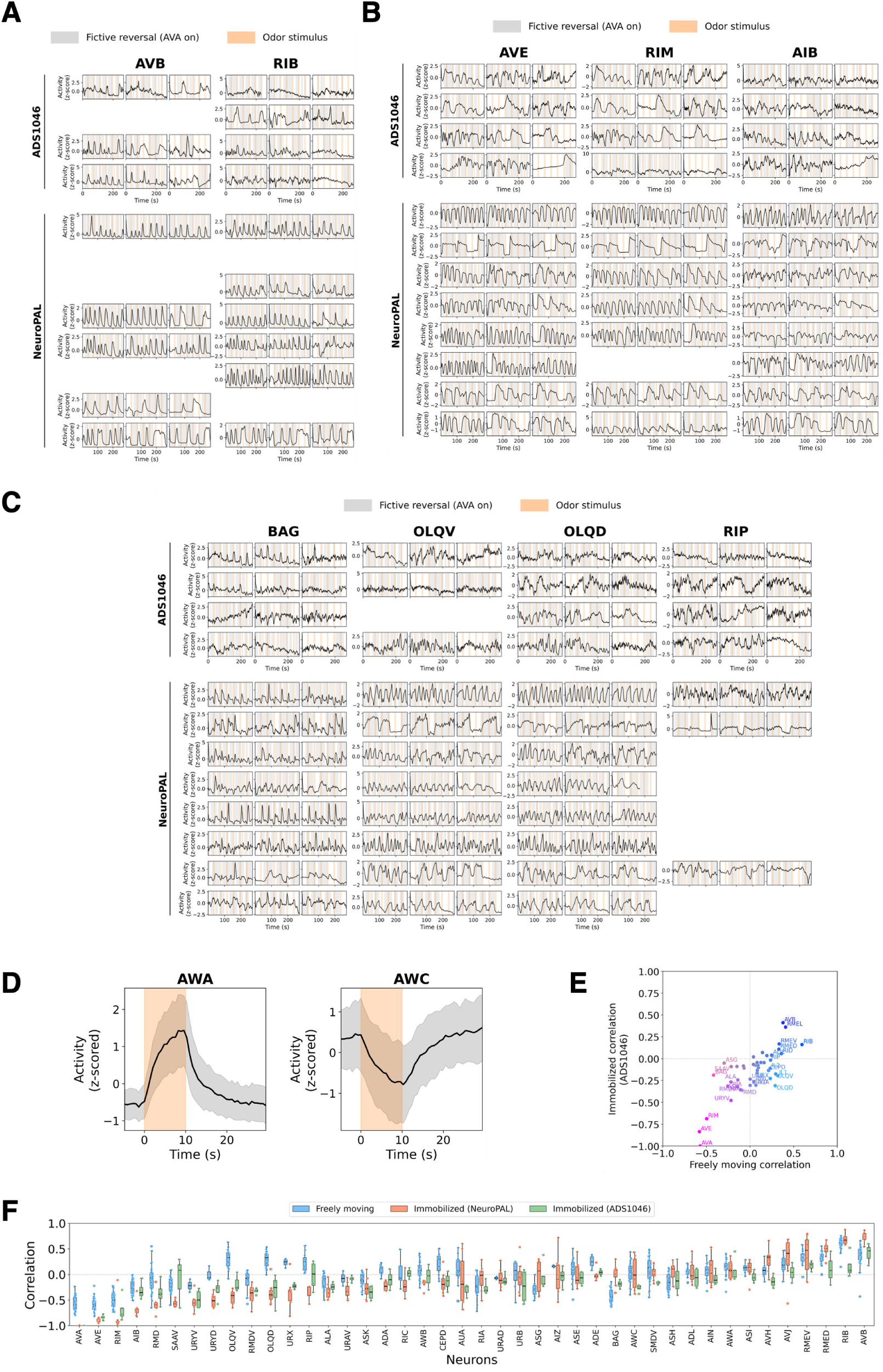
(**A**) Comparison of all immobilized recordings for ADS1046 and NeuroPAL for neurons with highest correlation with forward motion. (**B**) Comparison of all immobilized recordings for ADS1046 and NeuroPAL for neurons with highest correlation with backward motion. (**C**) Comparison of all immobilized recordings for ADS1046 and NeuroPAL for neurons with different correlation with (fictive) speed in immobilized vs. freely moving recordings. (**D**) Mean response to diacetyl pulse in microfluidics of AWA and AWC in ADS1046. Gray shaded area shows the standard deviation. Orange background shows the odor pulse. Both neurons were recorded in 4 animals. (**E**) Correlation of neuron activity with velocity in freely moving vs. fictive velocity in immobilized ADS1046, as in Fig. 5B. (**F**) Correlation of all neurons with velocity or fictive velocity, comparing distributions in freely moving (odor gradients only, 28 animals), immobilized NeuroPAL worms (8 animals), and ADS1046 worms (4 animals). NeuroPAL and freely moving data are the same as in Fig. 5A.

**Figure S5.**
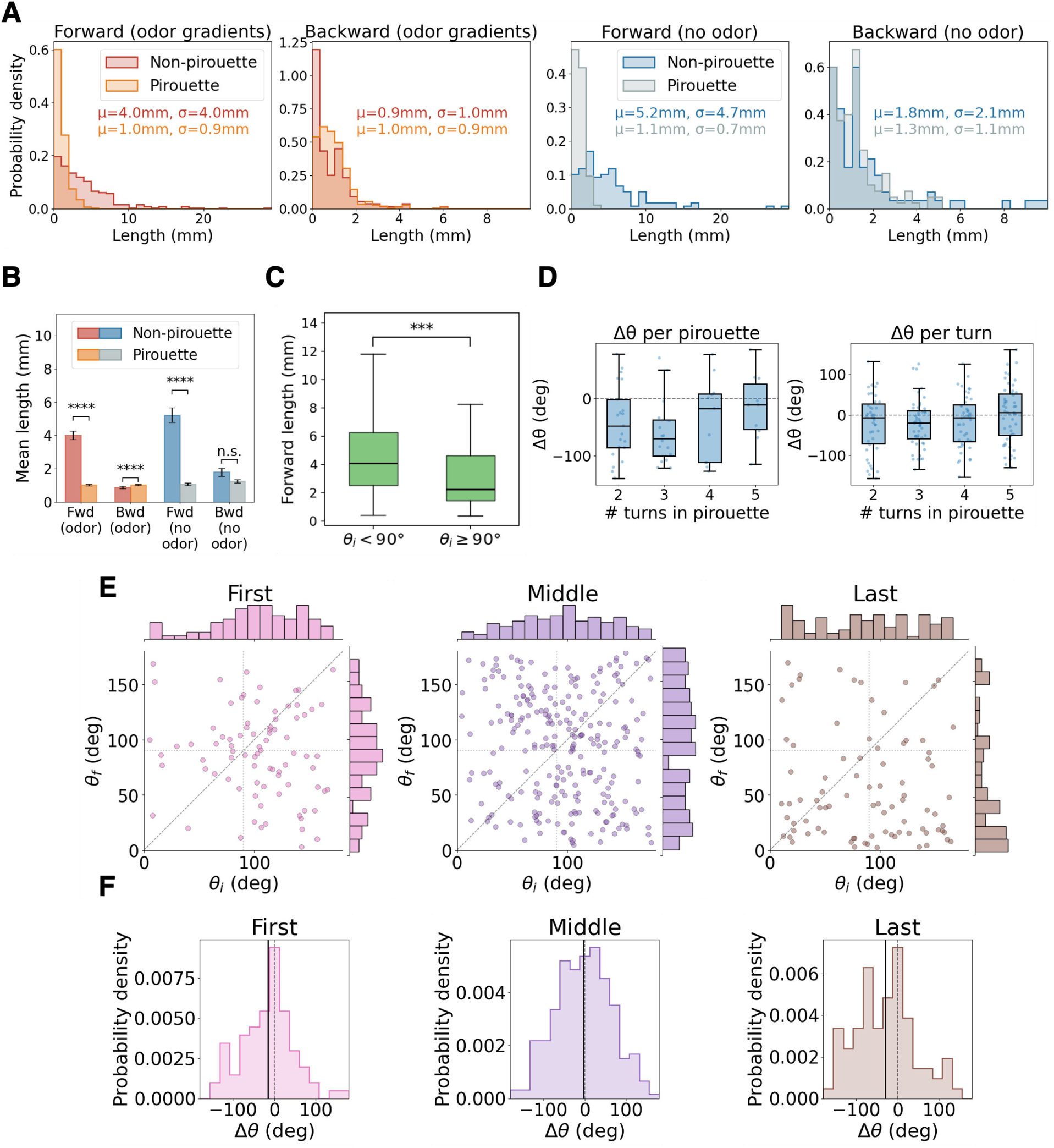
(**A**) Distribution of forward and backward segment lengths, measured in distance traveled, inside or outside a pirouette, separately for worms navigating odor gradients vs. in no-odor isotropic environments. Text shows the mean *µ* and standard deviation *σ* of the distributions. Odor gradients (28 animals): 605 forward segments (347 pirouette, 258 non-pirouette), 578 backward segments (418 pirouette, 160 non-pirouette). No odor (13 animals): 210 forward segments (93 pirouette, 117 non-pirouette), 198 backward segments (116 pirouette, 82 non-pirouette). (**B**) Mean length of forward and backward segments inside or outside a pirouette, separately for worms navigating in odor gradients vs. in no-odor isotropic environments. Statistical comparisons between non-pirouette and pirouette segment lengths were performed using two-sided Mann-Whitney U tests. Forward lengths were significantly shorter for pirouettes than for non-pirouettes both in odor gradients (p = 2.2 ×10^−40^, ****) and in isotropic environments (p = 4.1 ×10^−21^, ****). Backward lengths also differed significantly under odor (p = 6.2 ×10^−5^, ****), but were not significantly different in the no-odor condition (p = 0.21, n.s.). Significance is indicated as *p < 0.05, **p < 0.01, ***p < 0.001, ****p < 10^−4^; n.s., not significant. Animal and segment counts are the same as in (A). (**C**) Distribution of forward segment length for runs starting towards the odor source (*θ_i_ <* 90) or away from it (*θ_i_ >* 90). Forward runs exclude forward segments within a pirouette. Runs starting toward the odor were significantly longer than those starting away (two-sided Mann-Whitney U test; p = 3.13 ×10^−4^, ***). Mean distances: 5.05 mm (*θ_i_ <* 90°, n=152); 3.62 mm (*θ_i_ >* 90°, n=69). (**D**) Angle change Δ*θ* = *θ_f_* − *θ_i_* per pirouette (left) and per turn in a pirouette (right) as a function of pirouette length for animals in odor gradients (28 animals, 69 pirouettes, 271 reversals inside pirouettes). Angle change per pirouette is measured between the mean angle in the 5 seconds after the last reversal and in the 5 seconds before the first reversal. Angle change per reversal is measured between the 5 seconds before the reversal and the 5 seconds after the reversal. Statistical comparisons between all pairs were performed using two-sided Mann-Whitney U tests; all pairs were above significance threshold of p < 0.05. (**E**) Angle at the start and end of first (n=84), middle (n=248), or last reversals in a pirouette (n=86). Counts differ for first and last since the recording might start or end in the middle of a pirouette. (**F**) Distribution of angle differences Δ*θ* at the start and end of individual reversals in a pirouette, separating first, last, and middle reversals. Mean angle differences: -15.6° (first, n=84), -2.6° (middle, n=248), and -29.5° (last, n=86).

**Figure S6.**
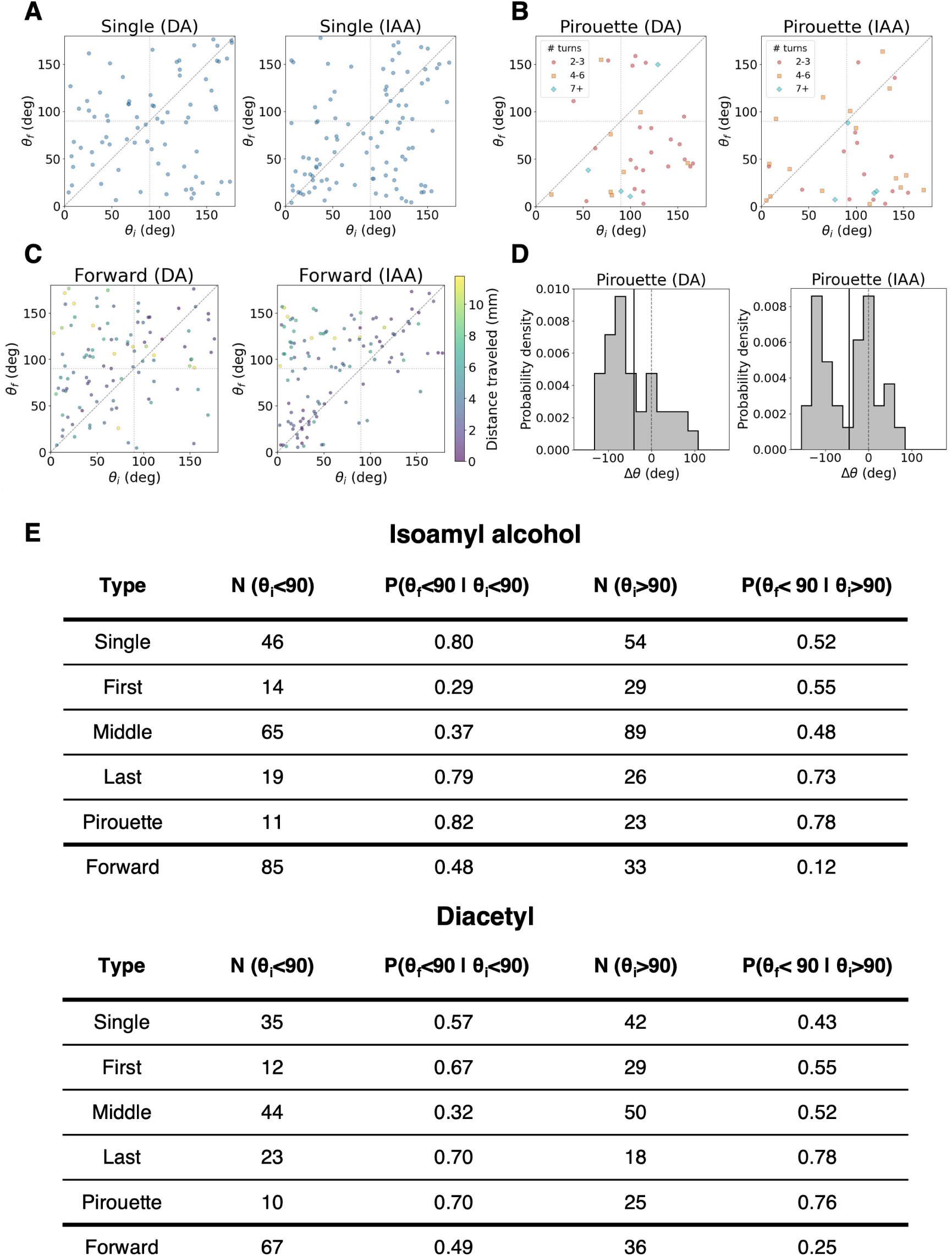
(**A**) Angle to odor at the start and end of single turns, separated by odor (DA: diacetyl, 76 turns; IAA: isoamyl alcohol, 100 turns). *θ_i_* is the mean angle to odor for the 5 seconds before the reversal and *θ_f_* is the mean angle to odor in the 5 seconds after the end of the reversal. (**B**) Angle to odor at the start and end of pirouettes, separated by odor (DA: 35 pirouettes; IAA: 34 pirouettes). Markers show the number of turns in the pirouette. *θ_i_* is the mean angle to odor for the 5 seconds before the first reversal in a pirouette and *θ_f_* is the mean angle to odor in the 5 seconds after the end of the last reversal in the pirouette. (**C**) Angle to odor at the start and end of forward segments, separated by odor (DA: 103 segments; IAA: 118 segments). *θ_i_* is the mean angle to odor in the first 5 seconds of the forward segments and *θ_f_* is the mean angle to odor in the last 5 seconds of the forward segment. Marker color shows the distance traveled during the forward segment (mm). (**D**) Histogram of angle change Δ*θ* = *θ_f_* − *θ_i_* for pirouettes, separated by odor. Mean angle difference: -41.1° (n=35) for DA, -45.1° (n=34) for IAA. (**E**) Tables showing the number N of each type of turn and of forward segments, and the fraction of each one that end towards the odor, depending on whether they started towards or away from the odor. Forward runs exclude segments within a pirouette. Total pirouette numbers differ from first reversal and last reversal numbers because incomplete pirouettes at the start or end of an experiment are excluded from pirouette count. For these, we still count the first reversal (if pirouette is at the end of the experiment) or last reversal (if pirouette is at the start of the experiment).

**Figure S7.**
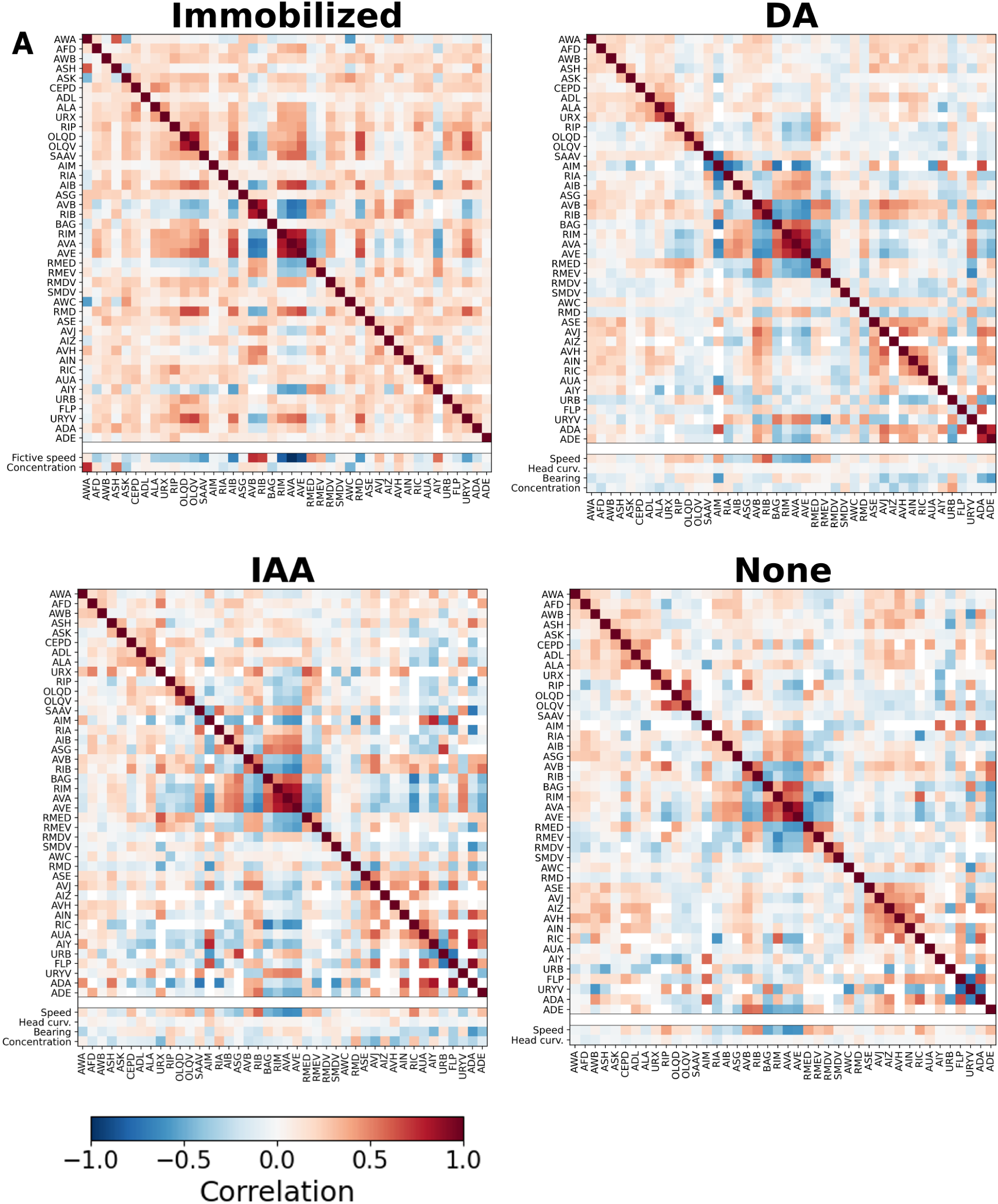
Pairwise correlation matrices for recorded neurons and stimulus and behavioral variables, averaged over worms experiencing the same condition. Matrices for immobilized worms, and crawling worms in gradients of diacetyl (DA), isoamyl alcohol (IAA), or no odorant are shown separately, sharing a color bar. Stimulus and behavioral variables shown vary by stimulus or experiment type. Same data as in Fig. 4A but sorted differently, using hierarchical clustering of the average of all freely moving worm matrices instead of immobilized worm matrices.

**Figure S8.**
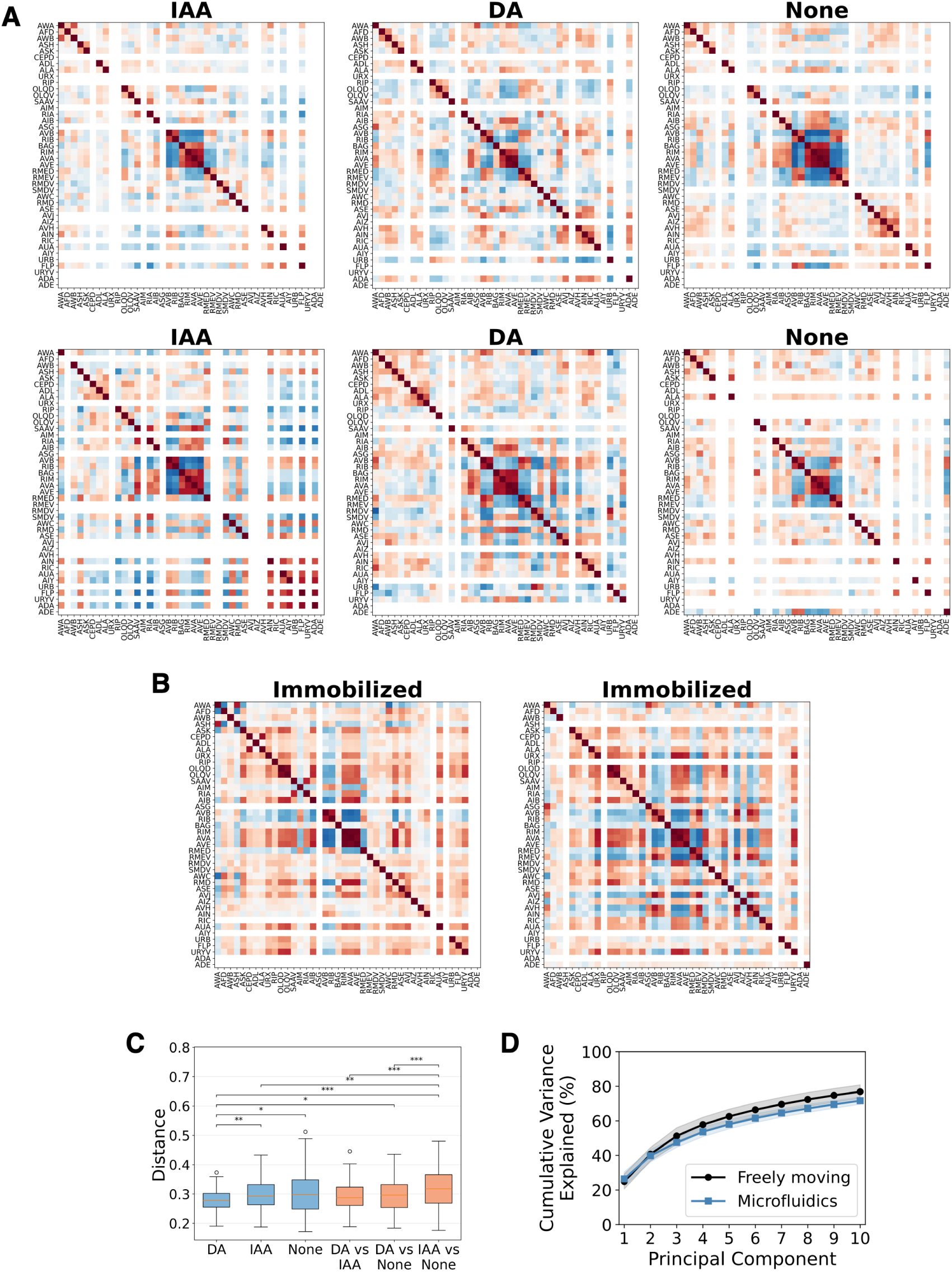
(**A**) Examples of correlation matrices corresponding to single datasets for worms crawling in odor gradients or isotropic environments. Set of neurons shown is the same as in Fig. 4A and Fig. S7. (**B**) Examples of correlation matrices corresponding to single datasets for immobilized worms, with the same set of neurons as in (A). (**C**) Distance between individual correlation matrices for freely moving worms separated by stimulus, measured as the Frobenius norm of the difference between individual matrices normalized by the number of neurons included in the matrix. Blue box plots show differences between freely moving worms experiencing the same stimulus condition (diacetyl gradients, isoamyl alcohol gradients, or no gradient). Orange box plots show differences between freely moving worms experiencing different stimuli. Statistical comparisons between distributions of pairwise correlation matrix distances were performed using a two-sided Mann-Whitney U test. Among within-stimulus comparisons, DA differed significantly from IAA (p = 4.32 ×10^−3^) and from no stimulus (p = 2.62 ×10^−2^), while IAA and no stimulus did not differ significantly (p = 0.74). Several between-stimulus distributions also differed from within-stimulus distributions: notably, the IAA vs. None distances were significantly larger than the DA within-stimulus distances (p = 1.39 ×10^−6^), the IAA within-stimulus distances (p = 3.14 ×10^−3^), the DA vs. IAA between-stimulus distances (p = 9.62 ×10^−6^), and the DA vs. None between-stimulus distances (p = 8.16 ×10^−4^). The DA within-stimulus distances also differed from the DA vs. None distances ( p = 2.18 ×10^−2^). All remaining pairwise comparisons were not statistically significant (p > 0.05). Significance levels: *p < 0.05, **p < 0.01, ***p < 0.001. (**D**) Cumulative variance explained by first 10 principal components in freely moving and immobilized data. Cumulative variance explained is averaged for all the freely moving datasets in odor gradients (28 animals) or all the immobilized datasets experiencing diacetyl pulses (8 animals). Shaded region corresponds to standard deviation. Principal component analysis is done after preprocessing; for freely moving worms, one of the preprocessing steps is removing the first principal component, since we find that it corresponds to motion artifacts.

**Figure S9.**
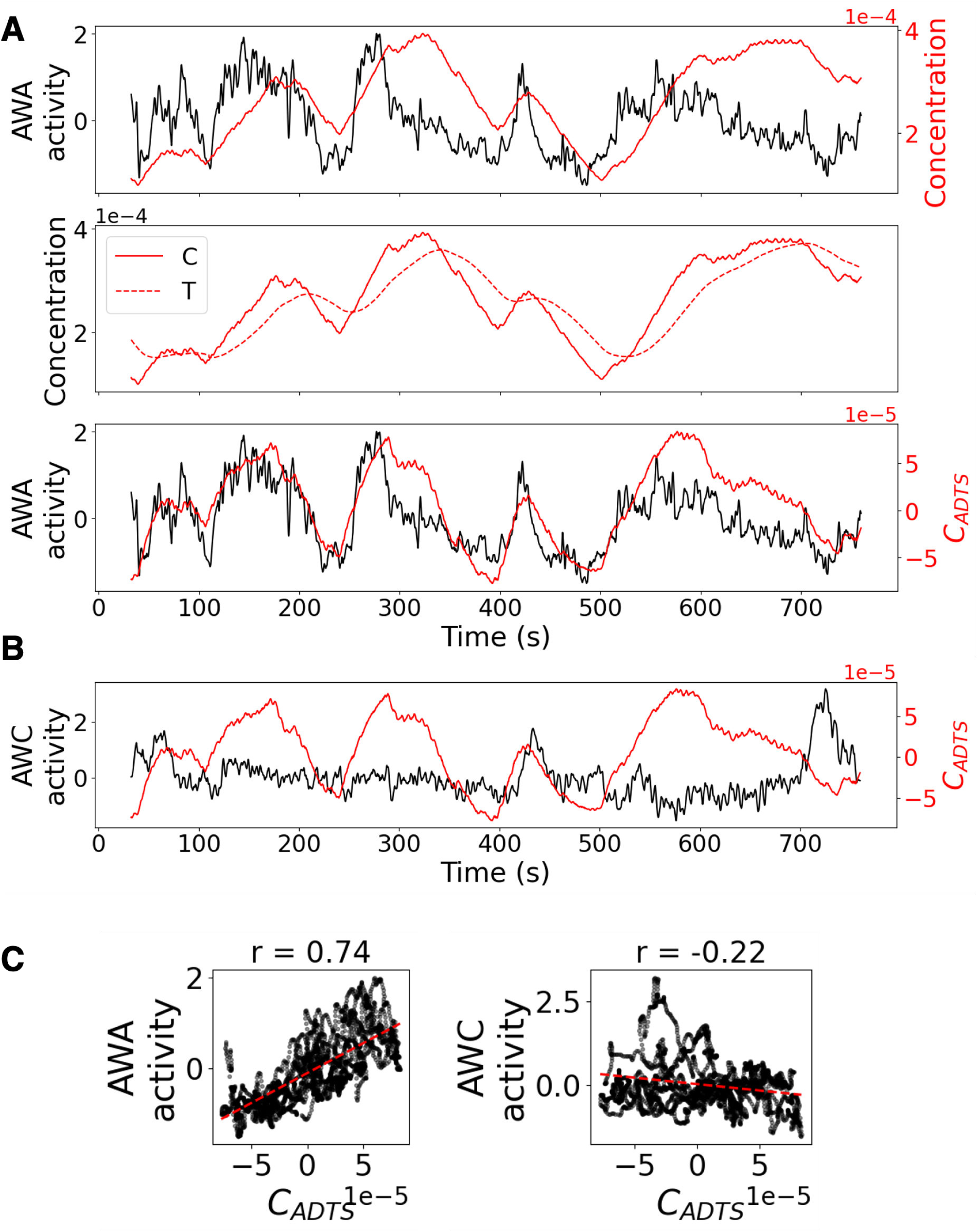
Mirroring Fig. 6 C-E for a different animal in a diacetyl gradient. (**A**) (Top) AWA and raw concentration on the same plot for a worm navigating towards diacetyl. (Middle) Relationship between *C* and *T* from the ADTS model. We show *T* using the mean time constant for diacetyl, *τ* = 29.6 seconds. (Bottom) AWA activity and *C_ADTS_* plotted together. Data are the same as in (C), showing correlation of 0.74 between AWA activity and *C_ADTS_*. (**B**) AWC activity plotted with *C_ADTS_*. They are slightly anti-correlated, with a correlation of −0.22 (shown in (C)). (**C**) Scatter plots showing linear relation between *C_ADTS_* and AWA activity or AWC activity. Correlation is positive for AWA and negative for AWC. Red dashed lines show a linear fit to the data.

**Figure S10.**
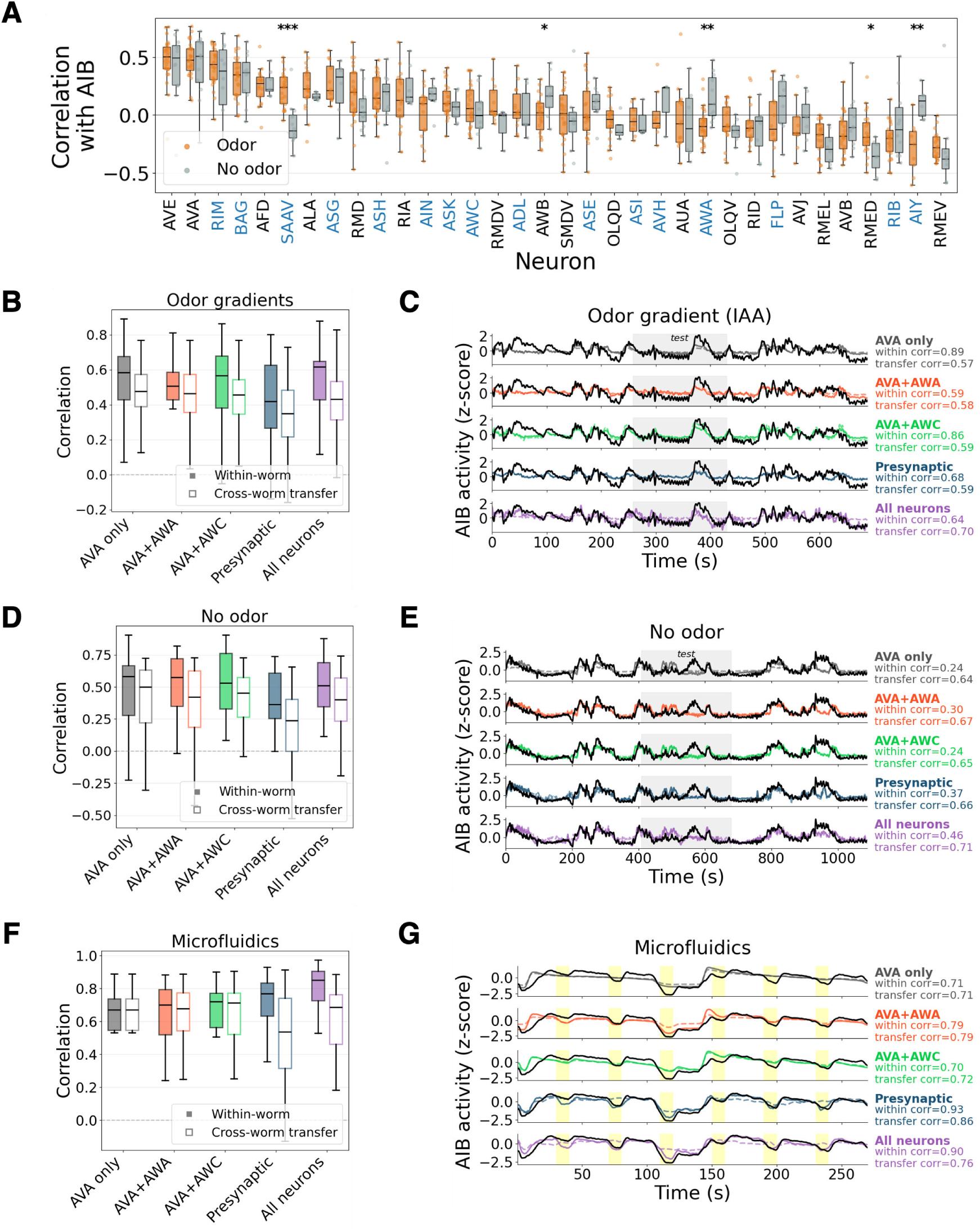
(**A**) Distribution of pairwise correlation between AIB and every other neuron in worms crawling in diacetyl gradients (12 animals), isoamyl alcohol gradients (16 animals), or isotropic environments (13 animals). Only neurons present in at least 3 datasets per stimulus are shown, sorted by median correlation in IAA datasets. Significance markings show results of two-sided Mann-Whitney U tests on the per-worm correlation values. Significance levels: *p<0.05, **p<0.01, ***p<0.001. Neurons presynaptic to AIB are shown in blue, using chemical synapse annotations in the union of the two adult connectomes in Witvliet et al. ^20^. (**B**) Correlation between linear model prediction and ground truth AIB activity in worms crawling in odor gradients, for models using different sets of neurons to fit activity (AVA only, AVA and AWA, AWA and AWC, neurons presynaptic to AIB, or all neurons recorded; see Methods). Within-worm distributions are computed for test data in the same animal used for training and cross-worm transfer distributions are computed for test data in other animals. (**C**) Example AIB activity trace and predictions from all models in (B) on a worm crawling in an odor gradient. Correlation values are computed on the test data. The test region is indicated with gray shading. Transfer shows the best prediction across all other worms used for training. (**D**) Correlation between linear model prediction and ground truth AIB activity in worms crawling in isotropic environments, for models using different sets of neurons to fit activity (AVA only, AVA and AWA, AWA and AWC, neurons presynaptic to AIB, or all neurons recorded; see Methods). Within-worm distributions are computed for test data in the same animal used for training and cross-worm transfer distributions are computed for test data in other animals. (**E**) Example AIB activity trace and predictions from all models in (D) on a worm crawling in an isotropic environment. Correlation values are computed on the test data. The test region is indicated with gray shading. Transfer shows the best prediction across all other worms used for training. (**F**) Correlation between linear model prediction and ground truth AIB activity in worms immobilized in microfluidic devices, for models using different sets of neurons to fit activity (see Methods). Within-worm distributions are computed for test data in the same animal used for training and cross-worm transfer distributions are computed for test data in other animals. (**G**) Example AIB activity trace and predictions from all models in (F) on a worm immobilized in a microfluidic device. Correlation values are computed on the test set (run 2). Yellow shading shows the odor pulses. Transfer shows the best prediction across all other worms used for training.

**Figure S11.**
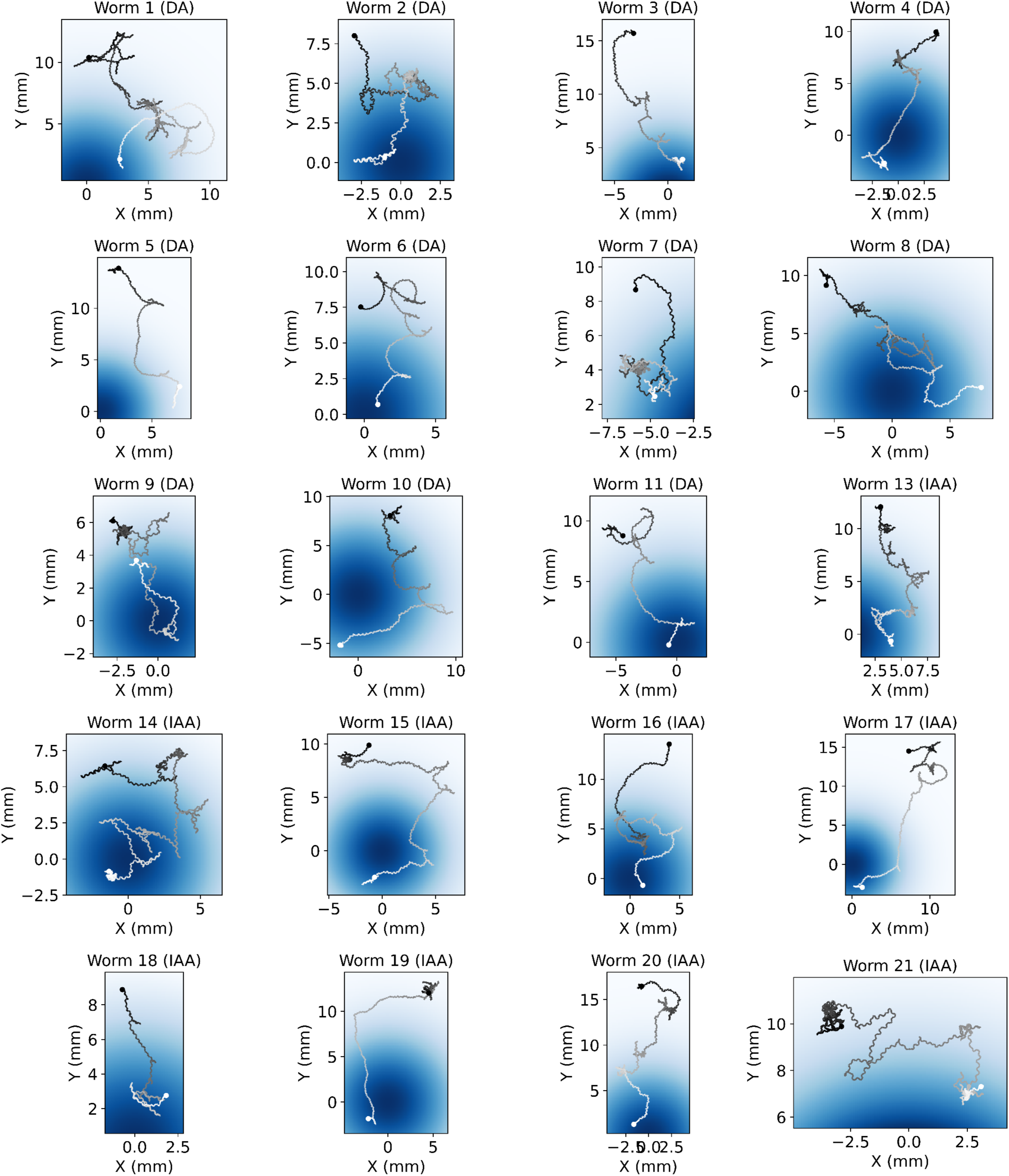
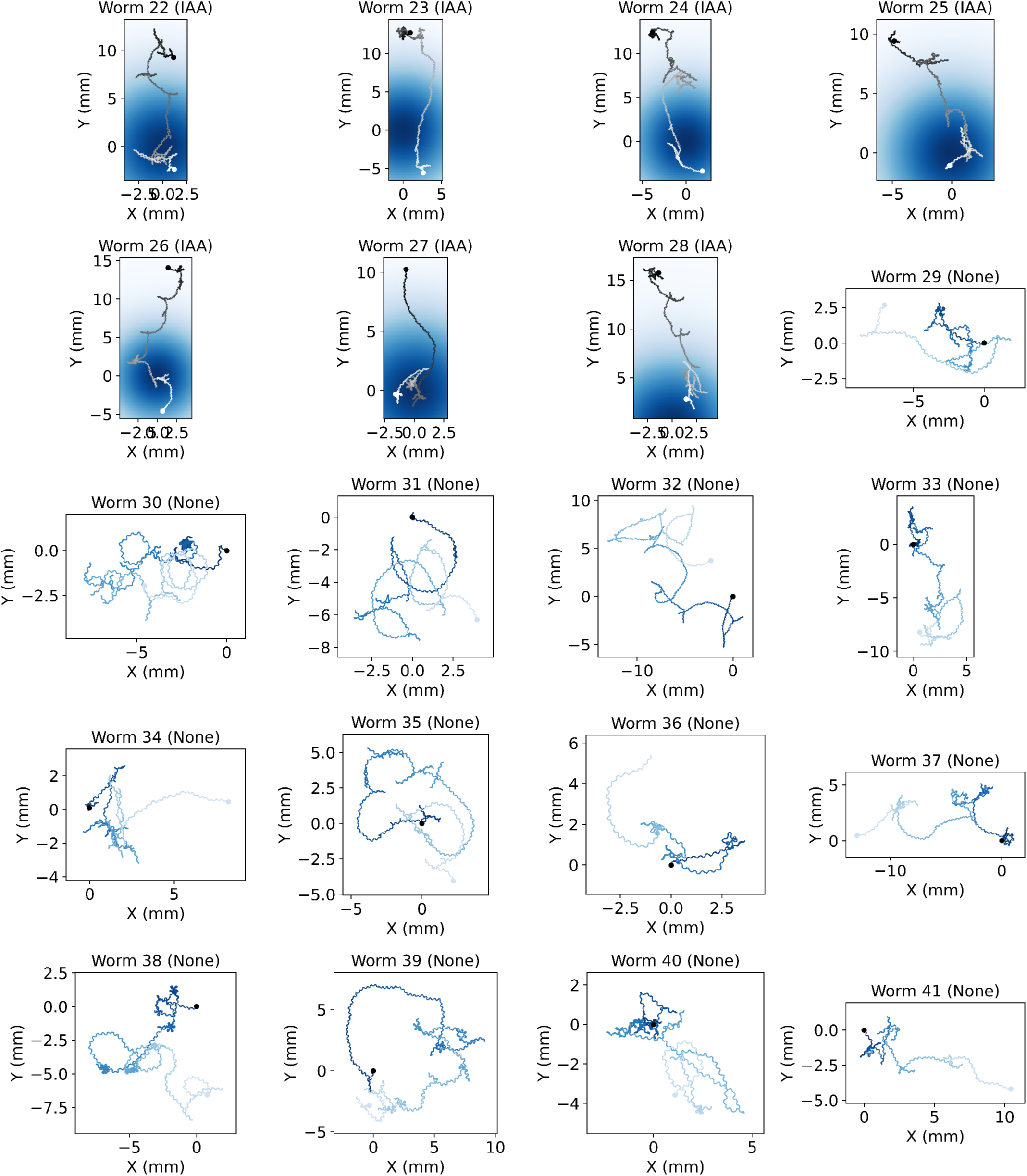
Trajectories on spatial odor gradients or isotropic environments. Dark dot marks the start and light dot marks the end. Background color shows the odor concentration (in arbitrary units). Darker colors represent higher concentrations. Worm numbers are the same as in Fig. S12. Worm 12 not shown; see main Fig. 1C instead.

**Figure S12.**
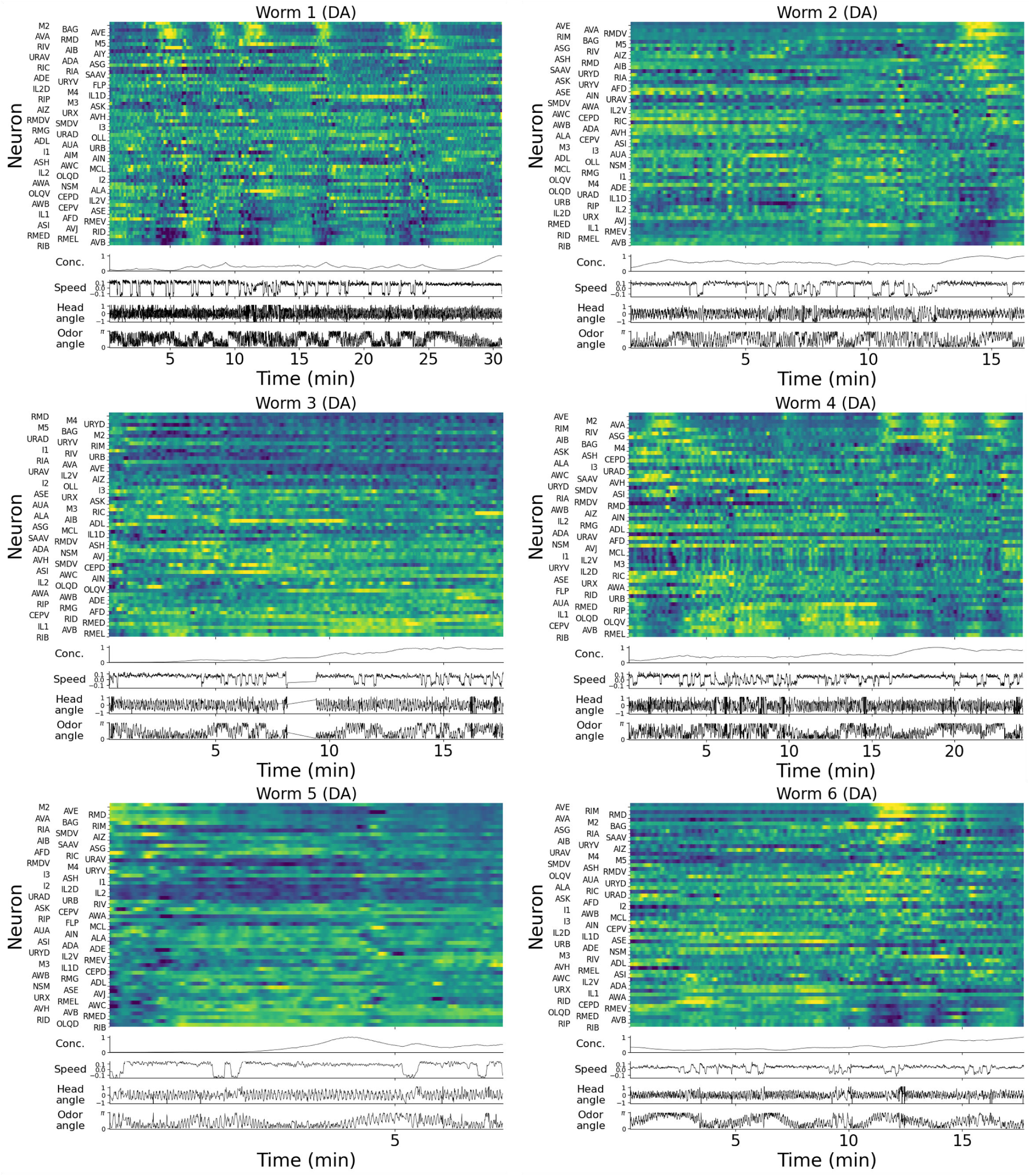
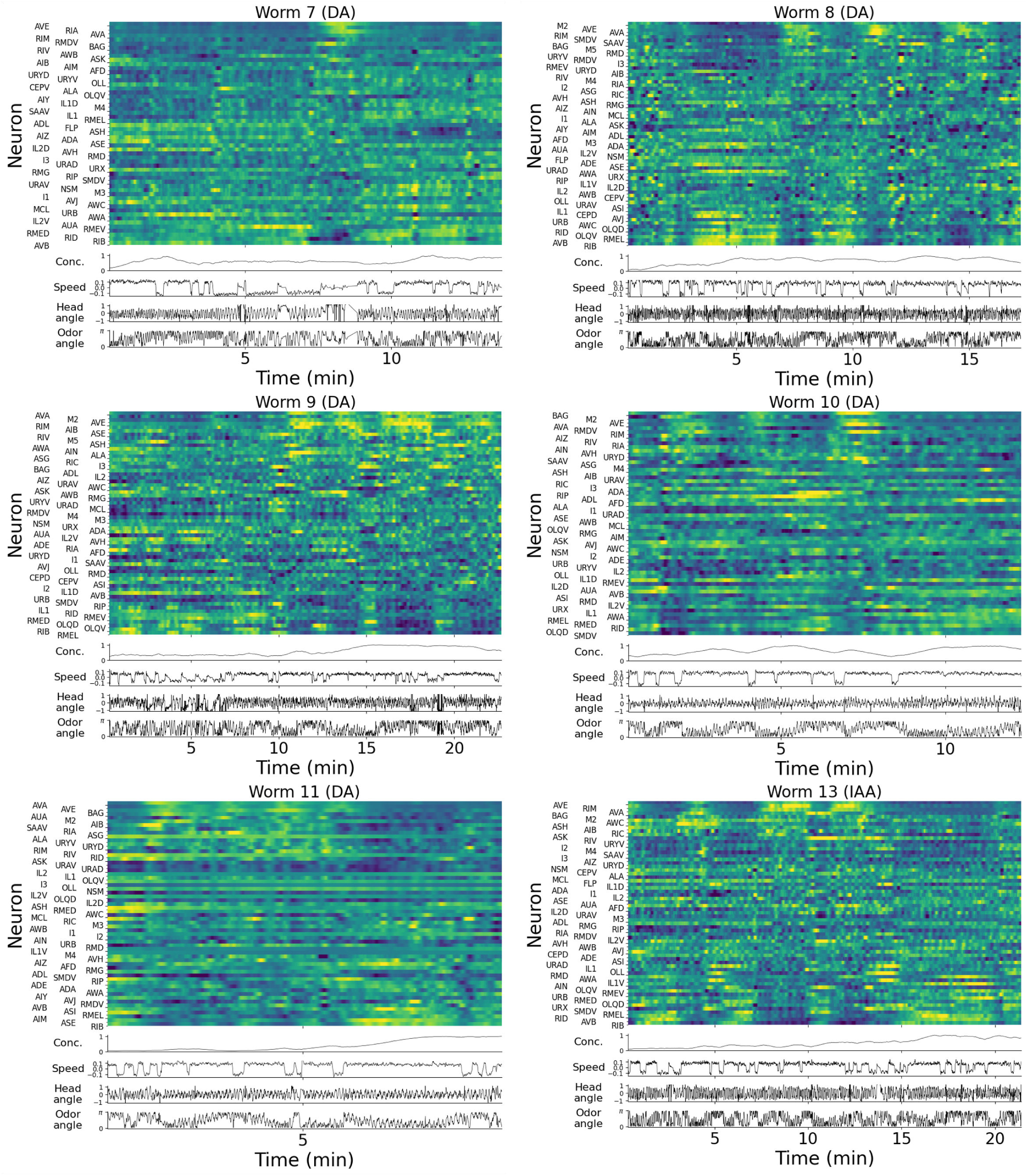
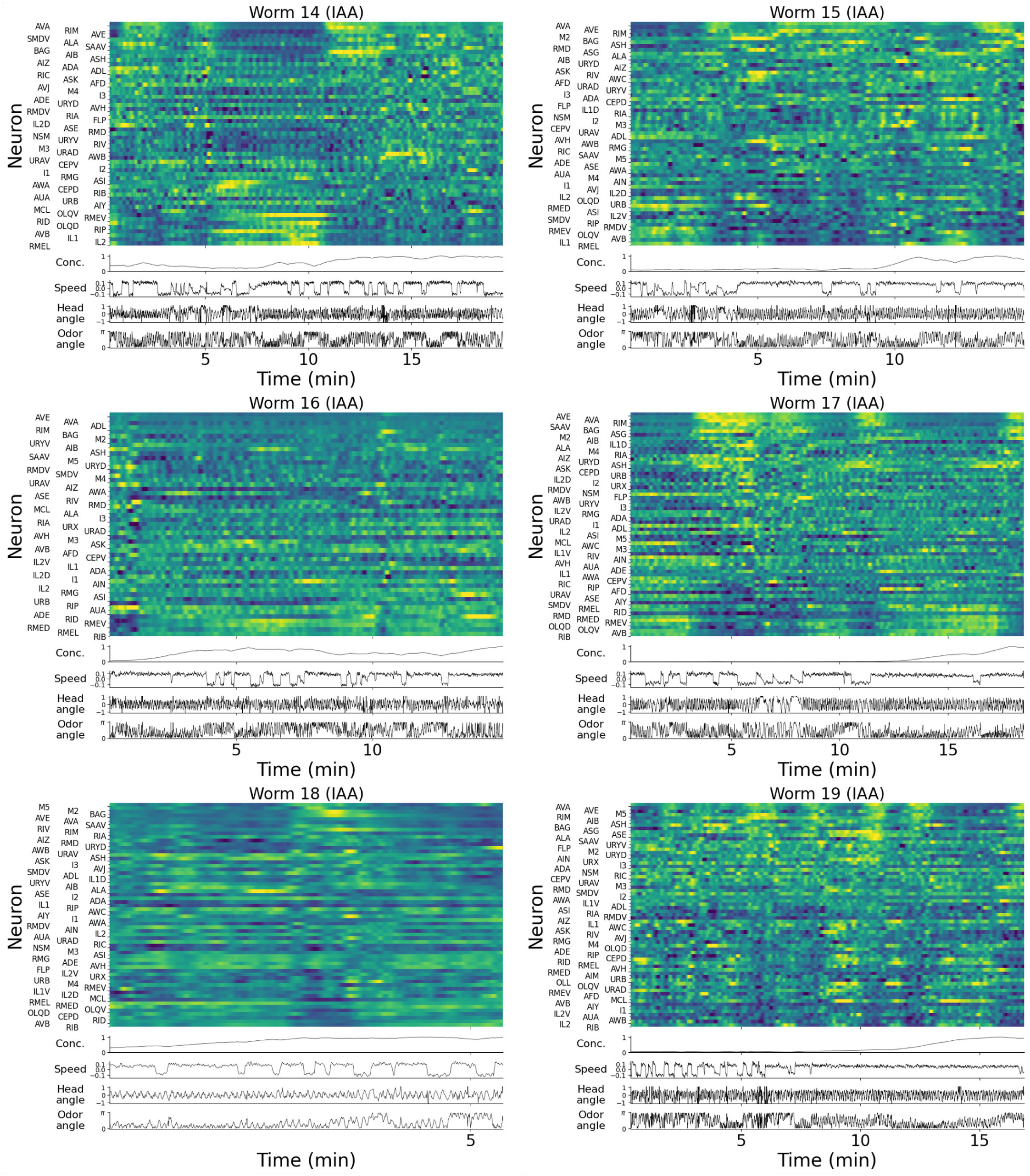
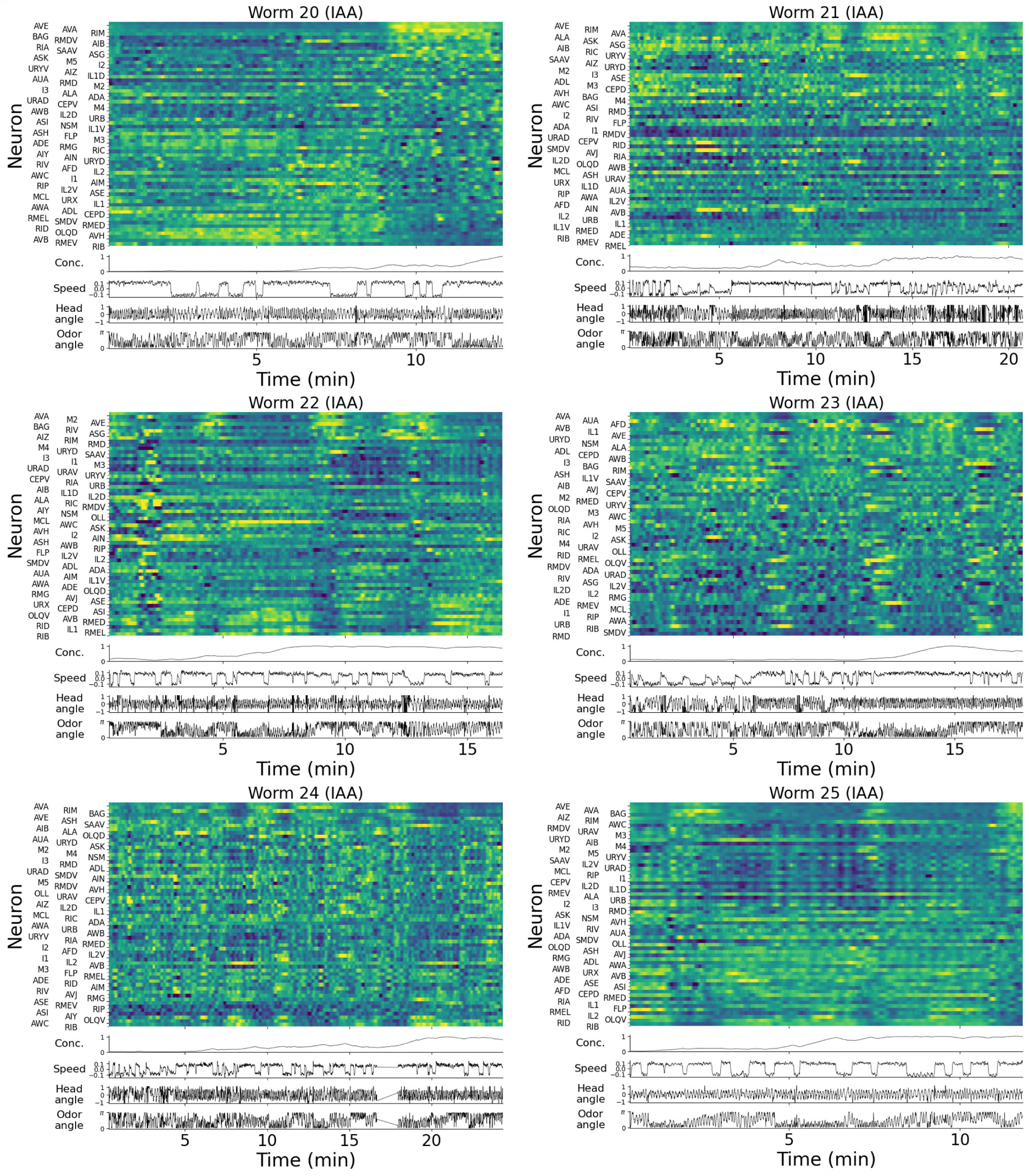
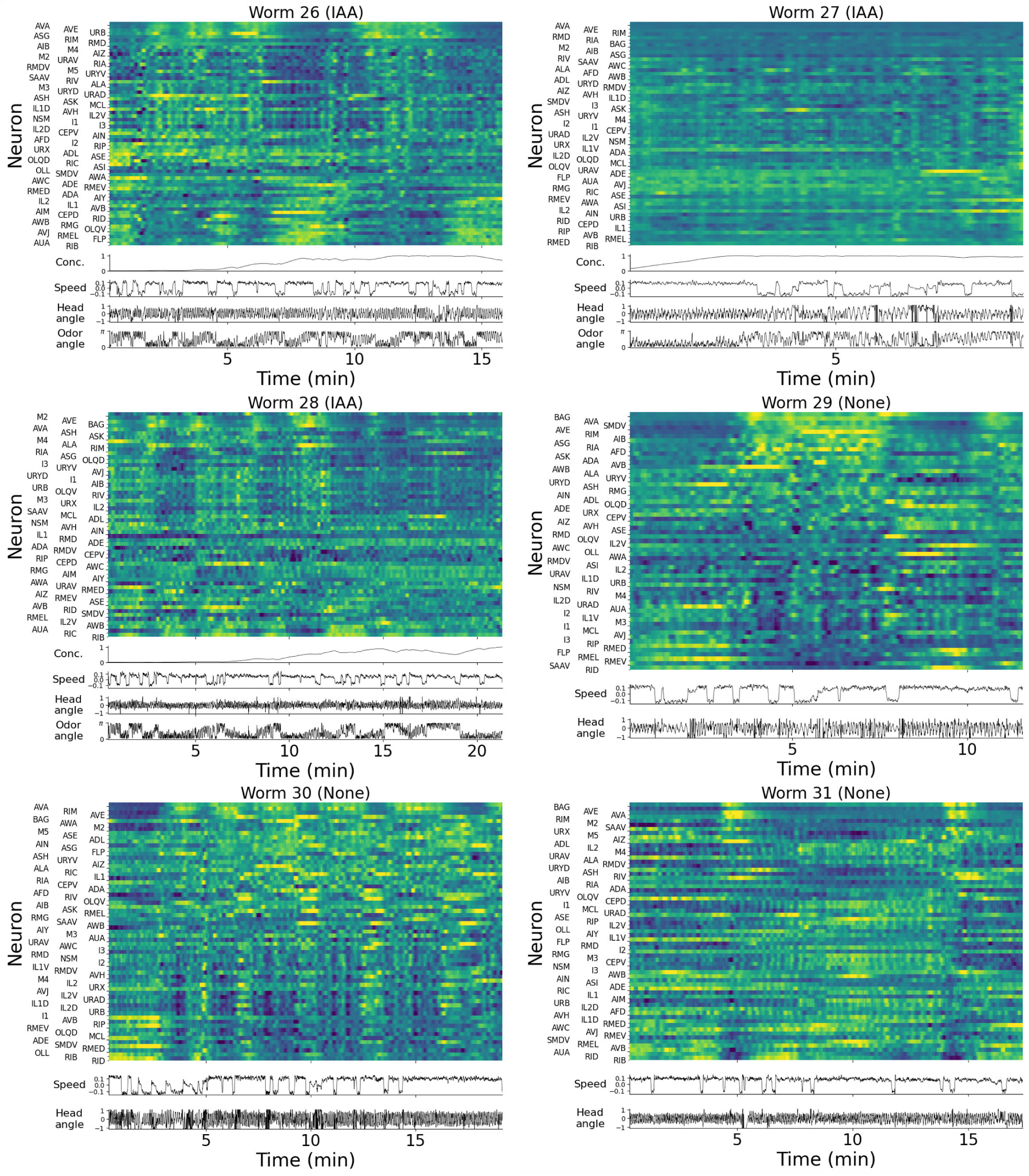
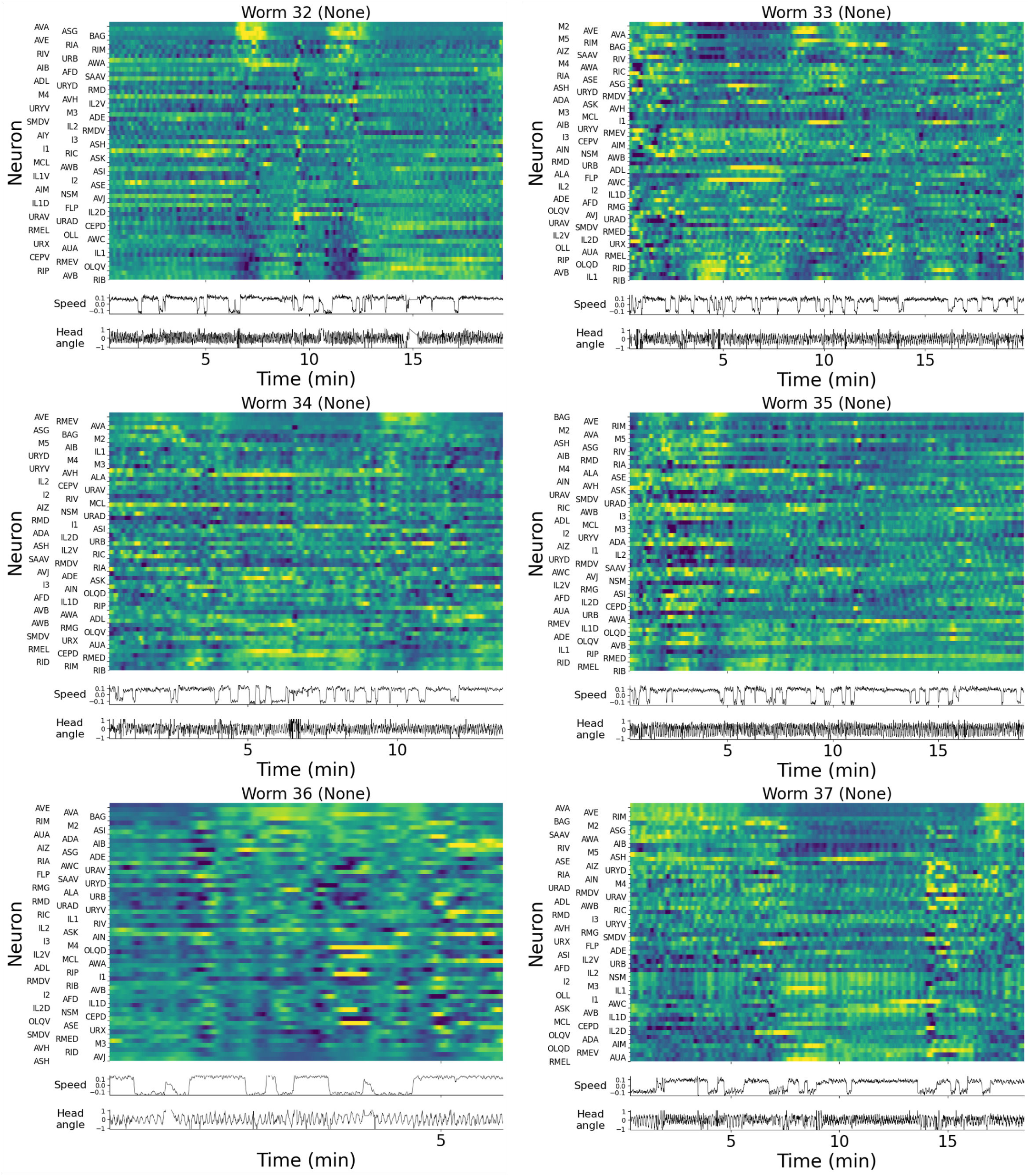
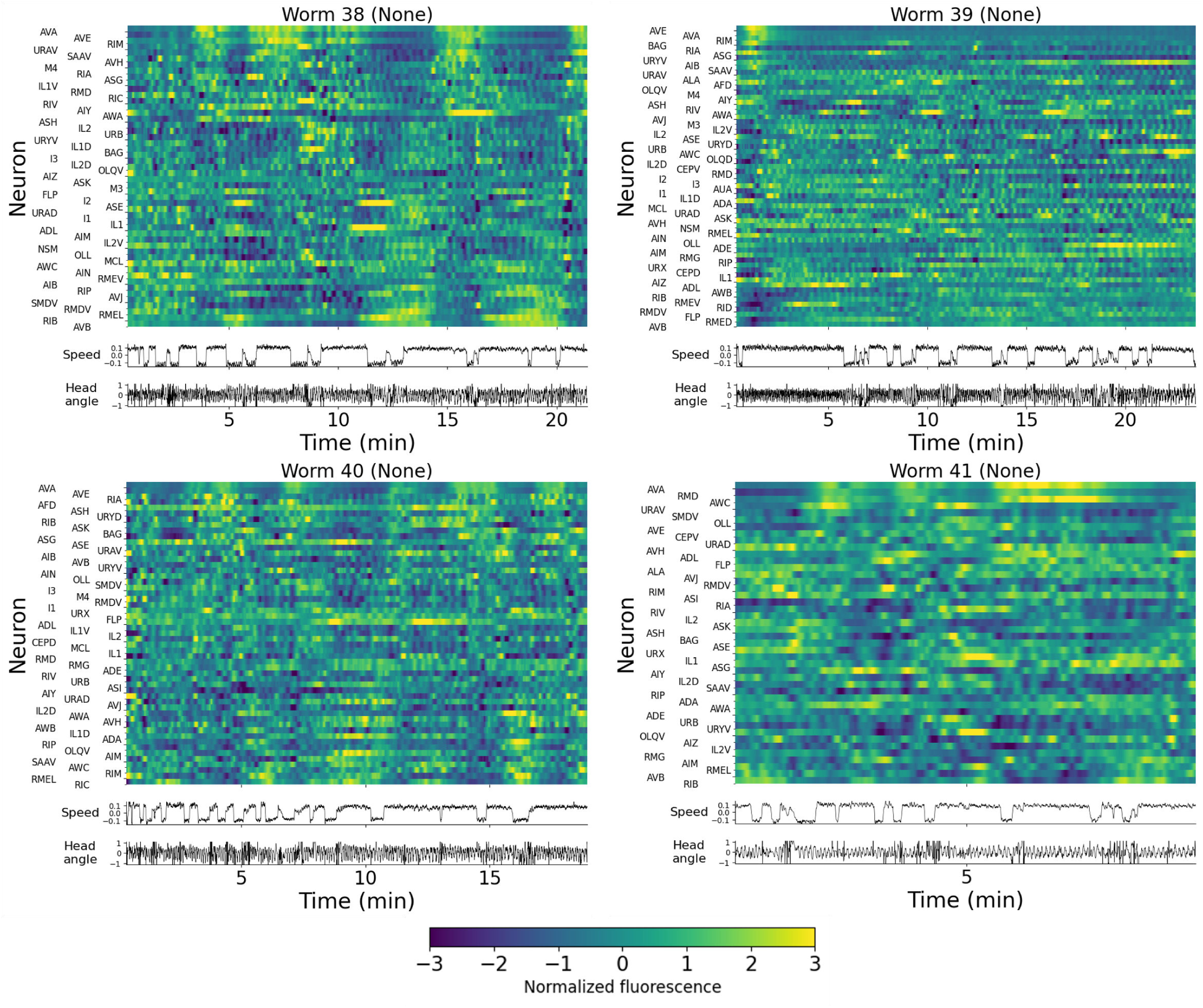
Activity heatmaps of all annotated neurons in each animal crawling in an odor gradient or isotropic environment. Only neurons that were identified are shown. Concentration (a.u., only odor gradient worms), velocity (mm/s), head angle (a.u.), and angle to odor (rad, only odor gradient worms) measurements are shown using the same time axis. Neuron activity is shown as z-score normalized fluorescence, with a color bar shared across all plots. Worm numbers are the same as in Fig. S11. Worm 12 not shown; see main Fig. 1E instead.

**Figure S13.**
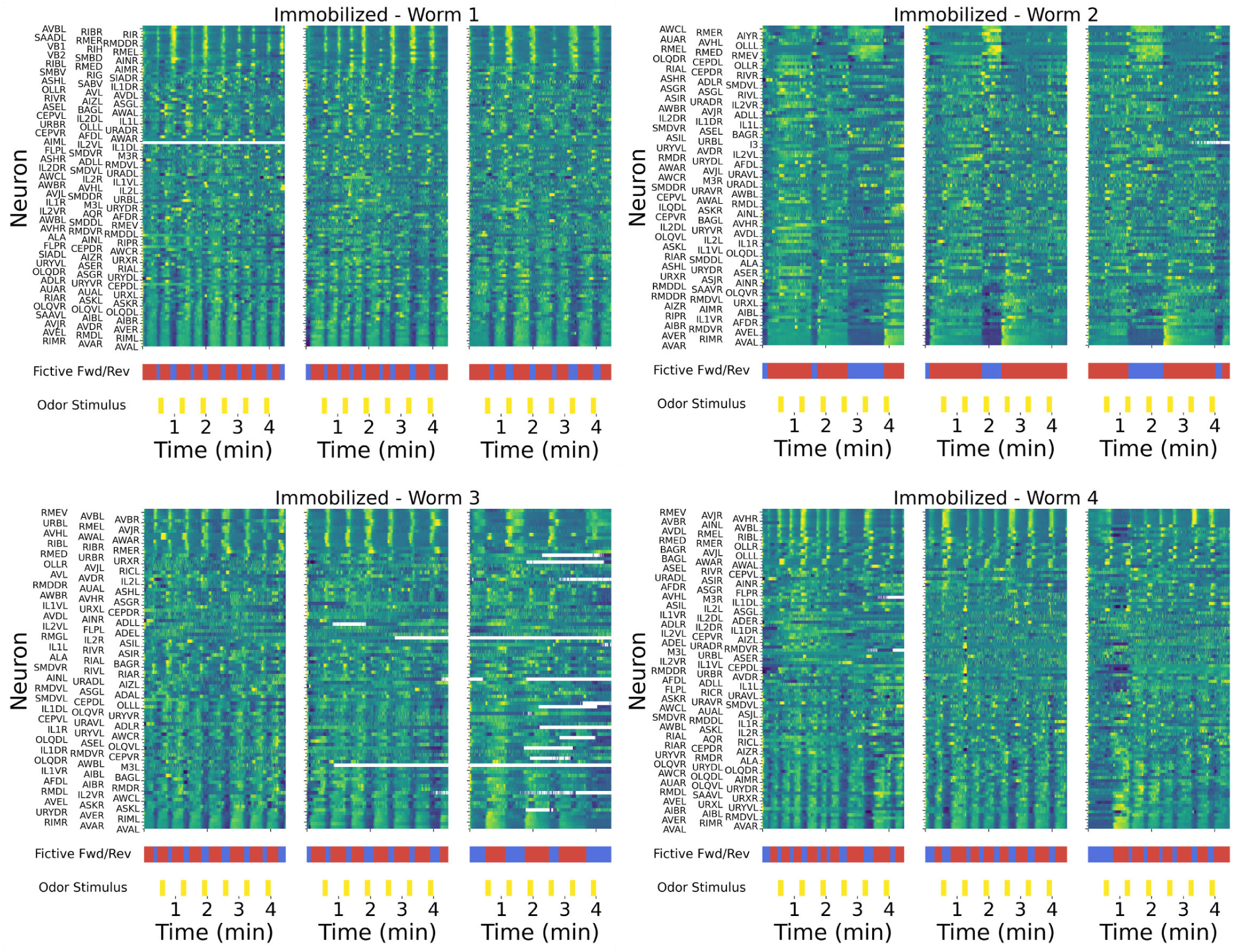
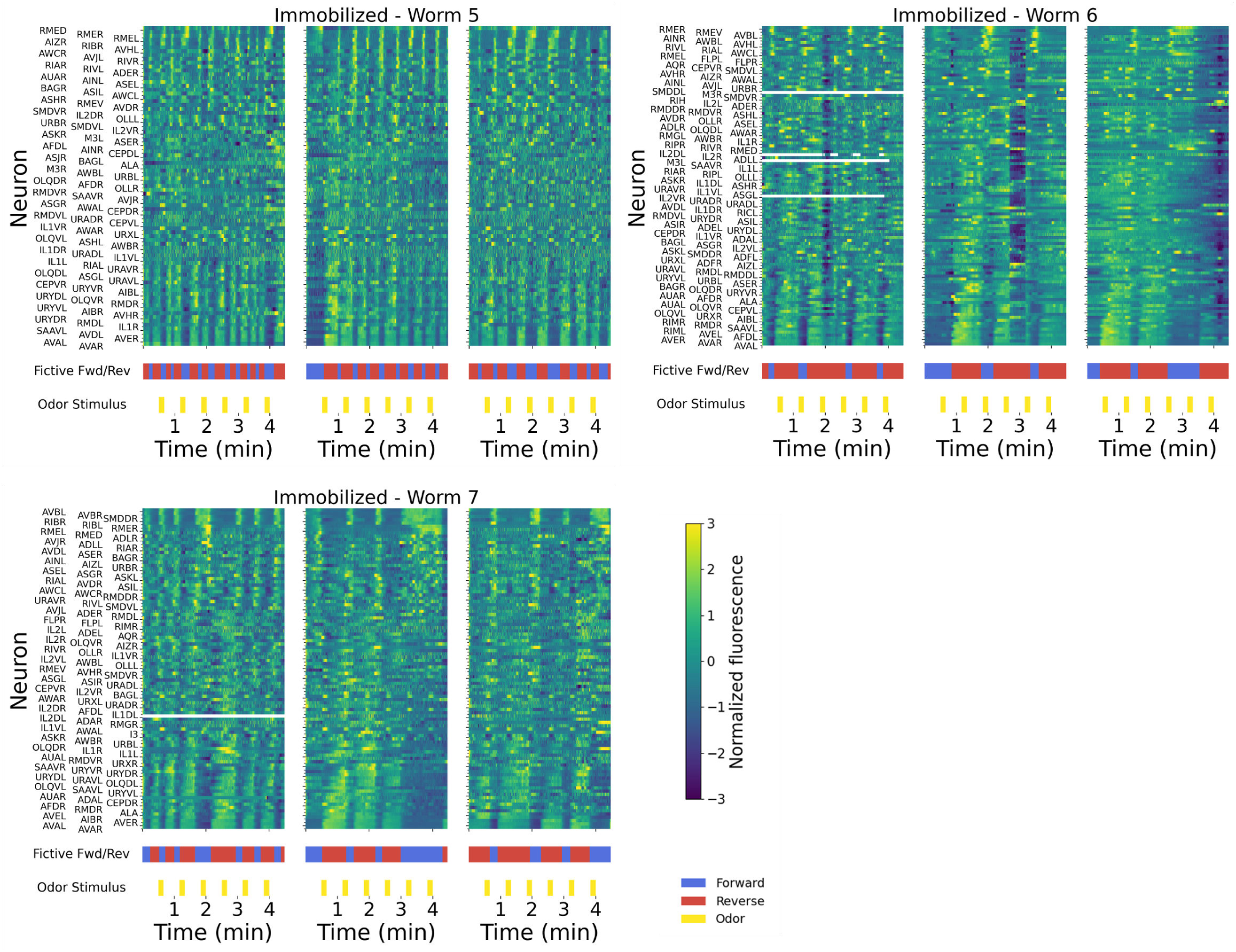
Neuron activity heatmap for all annotated neurons in each immobilized animal, showing 3 sequences of 6 diacetyl pulses each. Odor stimulus pattern and fictive behavior annotations are also shown. Neuron activity is shown as z-score normalized fluorescence, with a color bar shared across all plots. Worm 8 not shown; see main Fig. 1F instead.

## Notes

### Competing Interest Statement

The authors have declared no competing interest.

### Funding

- U.S. National Science Foundation: IOS-1452593, 1806818, IOS-1555914
- National Institutes of Health: 1UF1NS111697-01, U01NS132158
- Canadian Institutes of Health Research: 154274

## References

1. K. L. Briggman and W. B. Kristan. Multifunctional pattern-generating circuits. Annual Review of Neuroscience, 31(1):271–294, 2008.

2. Misha B. Ahrens, Jennifer M. Li, Michael B. Orger, Drew N. Robson, Alexander F. Schier, Florian Engert, and Ruben Portugues. Brain-wide neuronal dynamics during motor adaptation in zebrafish. Nature, 485(7399):471–477, 2012.

3. Georg B. Keller, Tobias Bonhoeffer, and Mark H”ubener. Sensorimotor mismatch signals in primary visual cortex of the behaving mouse. Neuron, 74(5):809–815, 2012.

4. Anmo J. Kim, Jamie K. Fitzgerald, and Gaby Maimon. Cellular evidence for efference copy in drosophila visuomotor processing. Nature Neuroscience, 18(9):1247–1255, 2015.

5. Lucas Pinto, Kanaka Rajan, Brian DePasquale, Stephan Y. Thiberge, David W. Tank, and Carlos D. Brody. Task-dependent changes in the large-scale dynamics and necessity of cortical regions. Neuron, 104(4):810–824.e9, 2019.

6. Sophie Aimon, Takeo Katsuki, Tongqiu Jia, Logan Grosenick, Michael Broxton, Karl Deisseroth, Terrence J. Sejnowski, and Ralph J. Greenspan. Fast near-whole-brain imaging in adult drosophila during responses to stimuli and behavior. PLoS Biology, 17(2):e2006732, 2019. ISSN 1545-7885. doi: 10.1371/journal.pbio.2006732.

7. Kevin Mann, Stephane Deny, Surya Ganguli, and Thomas R. Clandinin. Coupling of activity, metabolism and behaviour across the drosophila brain. Nature, 593(7858):244–248, 2021. ISSN 1476-4687. doi: 10.1038/s41586-021-03497-0.

8. Kelsey M. Hallinen, Ross Dempsey, Monika Scholz, Xinwei Yu, Ashley Linder, Francesco Randi, Anuj K. Sharma, Joshua W. Shaevitz, and Andrew M. Leifer. Decoding locomotion from population neural activity in moving *C. elegans*. eLife, 10:e66135, 2021. ISSN 2050-084X.

9. Adam A. Atanas, Jungsoo Kim, Ziyu Wang, Eric Bueno, McCoy Becker, Di Kang, Jungyeon Park, Talya S. Kramer, Flossie K. Wan, Saba Baskoylu, Ugur Dag, Elpiniki Kalogeropoulou, Matthew A. Gomes, Cassi Estrem, Netta Cohen, Vikash K. Mansinghka, and Steven W. Flavell. Brain-wide representations of behavior spanning multiple timescales and states in *C. elegans*. Cell, 186(19):4134–4151.e31, 2023. ISSN 0092-8674.

10. Charles Fieseler, Itamar Lev, Ulises Rey, Lukas Hille, Hannah Brenner, and Manuel Zimmer. An intrinsic neuronal manifold underlies brain-wide hierarchical organization of behavior in *C. elegans*. bioRxiv, 2025.

11. Albert Lin, Daniel Witvliet, Luis Hernandez-Nunez, Scott W. Linderman, Aravinthan D. T. Samuel, and Vivek Venkatachalam. Imaging whole-brain activity to understand behaviour. Nature Reviews Physics, 4(5):292–305, 2022. ISSN 2522-5820.

12. Eviatar Yemini, Albert Lin, Amin Nejatbakhsh, Erdem Varol, Ruoxi Sun, Gonzalo E. Mena, Aravinthan D. T. Samuel, Liam Paninski, Vivek Venkatachalam, and Oliver Hobert. Neuropal: A multicolor atlas for whole-brain neuronal identification in *C. elegans*. Cell, 184(1):272–288.e11, 2021. ISSN 0092-8674.

13. J. M. Gray, J. J. Hill, and C. I. Bargmann. A circuit for navigation in *Caenorhabditis elegans*. Proceedings of the National Academy of Sciences, 102(9):3184–3191, 2005.

14. Ephraim L. Tsalik and Oliver Hobert. Functional mapping of neurons that control locomotory behavior in *Caenorhabditis elegans*. Journal of Neurobiology, 56(2):178–197, 2003.

15. Sreekanth H. Chalasani, Nikos Chronis, Makoto Tsunozaki, Jesse M. Gray, Daniel Ramot, Miriam B. Goodman, and Cornelia I. Bargmann. Dissecting a circuit for olfactory behaviour in caenorhabditis elegans. Nature, 450(7166):63–70, 2007. ISSN 0028-0836.

16. Andrew Gordus, Navin Pokala, Sagi Levy, Steven W. Flavell, and Cornelia I. Bargmann. Feedback from network states generates variability in a probabilistic olfactory circuit. Cell, 161(2):215–227, 2015. ISSN 0092-8674.

17. Michael Hendricks, Heonick Ha, Nicolas Maffey, and Yun Zhang. Compartmentalized calcium dynamics in a *C. elegans* interneuron encode head movement. Nature, 487(7405):99–103, 2012. ISSN 0028-0836.

18. Amanda Ray and Andrew Gordus. Nonlinear integration of sensory inputs and behavioral state by a single neuron in *C. elegans*. Current Biology, 35(22):5534–5548.e4, 2025.

19. Ni Ji, Vivek Venkatachalam, Hillary Denise Rodgers, Wesley Hung, Taizo Kawano, Christopher M. Clark, Maria Lim, Mark J. Alkema, Mei Zhen, and Aravinthan D. T. Samuel. Corollary discharge promotes a sustained motor state in a neural circuit for navigation. eLife, 10:e68848, 2021.

20. Daniel Witvliet, Ben Mulcahy, James K. Mitchell, Yaron Meirovitch, Daniel R. Berger, Yuelong Wu, Yufang Liu, Wan Xian Koh, Rajeev Parvathala, Douglas Holmyard, Richard L. Schalek, Nir Shavit, Andrew D. Chisholm, Jeff W. Lichtman, Aravinthan D. T. Samuel, and Mei Zhen. Connectomes across development reveal principles of brain maturation. Nature, 596(7871):257–261, 2021. ISSN 0028-0836.

21. Emily Fryer, Sujay Guha, Lucero E. Rogel-Hernandez, Theresa Logan-Garbisch, Hodan Farah, Ehsan Rezaei, Iris N. Mollhoff, Adam L. Nekimken, Angela Xu, Lara Selin Seyahi, Sylvia Fechner, Shaul Druckmann, Thomas R. Clandinin, Seung Y. Rhee, and Miriam B. Goodman. A high-throughput behavioral screening platform for measuring chemotaxis by *C. elegans*. PLoS Biology, 22(6):e3002672, 2024.

22. Nikos Chronis, Manuel Zimmer, and Cornelia I. Bargmann. Microfluidics for in vivo imaging of neuronal and behavioral activity in *Caenorhabditis elegans*. Nature Methods, 4(9):727–731, 2007. ISSN 1548-7091.

23. Albert Lin, Shanshan Qin, Helena Casademunt, Min Wu, Wesley Hung, Gregory Cain, Nicolas Z. Tan, Raymond Valenzuela, Leila Lesanpezeshki, Vivek Venkatachalam, Cengiz Pehlevan, Mei Zhen, and Aravinthan D. T. Samuel. Functional imaging and quantification of multineuronal olfactory responses in *C. elegans*. Science Advances, 9(9):eade1249, 2023. doi: 10.1126/sciadv.ade1249.

24. Vivek Venkatachalam, Ni Ji, Xian Wang, Christopher Clark, James Kameron Mitchell, Mason Klein, Christopher J. Tabone, Jeremy Florman, Hongfei Ji, Joel Greenwood, Andrew D. Chisholm, Jagan Srinivasan, Mark Alkema, Mei Zhen, and Aravinthan D. T. Samuel. Pan-neuronal imaging in roaming *Caenorhabditis elegans*. Proceedings of the National Academy of Sciences, 113(8):E1082–E1088, 2016. ISSN 0027-8424.

25 . Vladislav Susoy, Wesley Hung, Daniel Witvliet, Joshua E. Whitener, Min Wu, Core Francisco Park, Brett J. Graham, Mei Zhen, Vivek Venkatachalam, and Aravinthan D. T. Samuel. Natural sensory context drives diverse brain-wide activity during *C. elegans* mating. Cell, 184(20):5122–5137.e17, 2021.

26. Core Francisco Park, Mahsa Barzegar-Keshteli, Kseniia Korchagina, Ariane Delrocq, Vladislav Susoy, Corinne L. Jones, Aravinthan D. T. Samuel, and Sahand Jamal Rahi. Automated neuron tracking inside moving and deforming *C. elegans* using deep learning and targeted augmentation. Nature Methods, 21(1):142–149, 2024. ISSN 1548-7091.

27. James Ryu, Amin Nejatbakhsh, Mahdi Torkashvand, Sahana Gangadharan, Maedeh Seyedolmohadesin, Jinmahn Kim, Liam Paninski, and Vivek Venkatachalam. Versatile multiple object tracking in sparse 2d/3d videos via deformable image registration. PLoS Computational Biology, 20(5):e1012075, 2024.

28. J. T. Pierce-Shimomura, T. M. Morse, and Shawn R. Lockery. The fundamental role of pirouettes in caenorhabditis elegans chemotaxis. Journal of Neuroscience, 19(21):9557–9569, November 1999.

29. Samuel Ward. Chemotaxis by the nematode caenorhabditis elegans: Identification of attractants and analysis of the response by use of mutants. Proceedings of the National Academy of Sciences, 70 (3):817–821, 1973. doi: 10.1073/pnas.70.3.817.

30. Jun Meng, Tosif Ahamed, Bin Yu, Wesley Hung, Sonia EI Mouridi, Zezhen Wang, Yongning Zhang, Quan Wen, Thomas Boulin, Shangbang Gao, and Mei Zhen. A tonically active master neuron modulates mutually exclusive motor states at two timescales. Science Advances, 10(15):eadk0002, 2024. doi: 10.1126/sciadv.adk0002.

31. M. Chalfie, J. E. Sulston, J. G. White, E. Southgate, J. N. Thomson, and S. Brenner. The neural circuit for touch sensitivity in *Caenorhabditis elegans*. Journal of Neuroscience, 5(4):956–964, 1985. ISSN 0270-6474.

32. Taizo Kawano, Michelle D. Po, Shangbang Gao, George Leung, William S. Ryu, and Mei Zhen. An imbalancing act: Gap junctions reduce the backward motor circuit activity to bias *C. elegans* for forward locomotion. Neuron, 72(4):572–586, 2011. ISSN 10974199. doi: 10.1016/j.neuron.2011.09.005.

33. Saul Kato, Harris S. Kaplan, Tina Schr”odel, Susanne Skora, Theodore H. Lindsay, Eviatar Yemini, Shawn Lockery, and Manuel Zimmer. Global brain dynamics embed the motor command sequence of Caenorhabditis elegans. Cell, 163(3):656–669, 2015. ISSN 10974172. doi: 10.1016/j.cell.2015.09.034.

34. Kevin S. Chen, Andrew M. Leifer, and Jonathan W. Pillow. State-switching navigation strategies in *C. elegans* are beneficial for chemotaxis. arXiv, 2025.

35. Ryuzo Shingai. Durations and frequencies of free locomotion in wild type and gabaergic mutants of caenorhabditis elegans. Neuroscience Research, 38(1):71–84, 2000.

36. Yuichi Iino and Kazushi Yoshida. Parallel use of two behavioral mechanisms for chemotaxis in caenorhabditis elegans. Journal of Neuroscience, 29(17):5370–5380, 2009.

37. Hiroshi Suzuki, Tod R. Thiele, Serge Faumont, Marina Ezcurra, Shawn R. Lockery, and William R. Schafer. Functional asymmetry in *Caenorhabditis elegans* taste neurons and its computational role in chemotaxis. Nature, 454(7200):114–117, July 2008. ISSN 14764687. doi: 10.1038/nature06927.

38. Dirk R. Albrecht and Cornelia I. Bargmann. High-content behavioral analysis of caenorhabditis elegans in precise spatiotemporal chemical environments. Nature Methods, 8(7):599–605, 2011.

39. Howard C. Berg and Douglas A. Brown. Chemotaxis in *Escherichia coli* analysed by three-dimensional tracking. Nature, 239(5374):500–504, 1972.

40. Yu Shen, Quan Wen, He Liu, Connie Zhong, Yuqi Qin, Gareth Harris, Taizo Kawano, Min Wu, Tianqi Xu, Aravinthan D. T. Samuel, and Yun Zhang. An extrasynaptic gabaergic signal modulates a pattern of forward movement in *C. elegans*. eLife, 5, 2016.

41. Linjiao Luo, Quan Wen, Jing Ren, Michael Hendricks, Marc Gershow, Yuqi Qin, Joel Greenwood, Edward R. Soucy, Mason Klein, Heidi K. Smith-Parker, Ana C. Calvó, Daniel A. Colón-Ramos, Aravinthan D. T. Samuel, and Yun Zhang. Dynamic encoding of perception, memory, and movement in a *C. elegans* chemotaxis circuit. Neuron, 82(5):1115–1128, 2014.

42. Beverly J. Piggott, Jie Liu, Zhaoyang Feng, Seth A. Wescott, and X. Z. Shawn Xu. The neural circuits and synaptic mechanisms underlying motor initiation in c. elegans. Cell, 147(4):922–933, 2011.

43. Joshua M. Kaplan and H. Robert Horvitz. A dual mechanosensory and chemosensory neuron in caenorhabditis elegans. Proceedings of the National Academy of Sciences, 90(6):2227–2231, 1993.

44. Katie S. Kindt, Veena Viswanath, L. J. Macpherson, Karla Quast, Hang Hu, Ardem Patapoutian, and William R. Schafer. Caenorhabditis elegans trpa-1 functions in mechanosensation. Nature Neuroscience, 10:568–577, 2007. doi: 10.1038/nn1886.

45. Miriam B. Goodman. Mechanosensation. WormBook, 2006. doi: 10.1895/wormbook.1.62.1. Review.

46. Elissa A. Hallem and Paul W. Sternberg. Acute carbon dioxide avoidance in caenorhabditis elegans. Proceedings of the National Academy of Sciences, 105(23):8038–8043, 2008.

47. Manuel Zimmer, Jesse M. Gray, Navin Pokala, Andy J. Chang, David S. Karow, Michael A. Marletta, Martin L. Hudson, David B. Morton, Nikos Chronis, and Cornelia I. Bargmann. Neurons detect increases and decreases in oxygen levels using distinct guanylate cyclases. Neuron, 61(6):865–879, 2009.

48. Julia Riedl, Charles Fieseler, and Manuel Zimmer. Tyraminergic corollary discharge filters reafferent perception in a chemosensory neuron. Current Biology, 32(14):3048–3058.e6, 2022.

49. Sarah G. Leinwand and Sreekanth H. Chalasani. Neuropeptide signaling remodels chemosensory circuit composition in *Caenorhabditis elegans*. Nature Neuroscience, 16(10):1461–1467, 2013. ISSN 1097-6256.

50. Sarah G. Leinwand, Claire J. Yang, Daphne Bazopoulou, Nikos Chronis, Jagan Srinivasan, and Sreekanth H. Chalasani. Circuit mechanisms encoding odors and driving aging-associated behavioral declines in *Caenorhabditis elegans*. Elife, 4(September 2015):1–26, 2015. ISSN 2050084X. doi: 10.7554/eLife.10181.

51. Cornelia I. Bargmann, Erika Hartwieg, and H. Robert Horvitz. Odorant-selective genes and neurons mediate olfaction in *C. elegans*. Cell, 74(3):515–527, August 1993. ISSN 00928674. doi: 10.1016/0092-8674(93)80053-H.

52. Alon Zaslaver, Idan Liani, Oshrat Shtangel, Shira Ginzburg, Lisa Yee, and Paul W. Sternberg. Hierarchical sparse coding in the sensory system of *Caenorhabditis elegans*. Proceedings of the National Academy of Sciences, 112(13):E1688–E1689, 2015.

53. Johannes Larsch, Donovan Ventimiglia, Cornelia I. Bargmann, and Dirk R. Albrecht. High-throughput imaging of neuronal activity in *Caenorhabditis elegans*. Proceedings of the National Academy of Sciences, 110(45), November 2013. ISSN 00278424. doi: 10.1073/pnas.1318325110.

54. Sagi Levy and Cornelia I. Bargmann. An adaptive-threshold mechanism for odor sensation and animal navigation. Neuron, 105(3):534–548.e13, 2020.

55. John Graham White, Eileen Southgate, J. N. Thomson, and Sydney Brenner. The structure of the nervous system of the nematode *Caenorhabditis elegans*. *Philosophical Transactions of the Royal Society of London. Series B*, Biological Sciences, 314(1165):1–340, 1986.

56. Anne C. Hart, Shannon Sims, and Joshua M. Kaplan. Synaptic code for sensory modalities revealed by c. elegans glr-1 glutamate receptor. Nature, 378:82–85, 1995. doi: 10.1038/378082a0.

57. Jarred Sanders, Stanislav Nagy, Graham Fetterman, Charles Wright, Millet Treinin, and David Biron. The Caenorhabditis elegans interneuron ALA is (also) a high-threshold mechanosensor. BMC neuroscience, 14(1):156–156, 2013.

58. Andres V. Maricq, Erin Peckol, Monica Driscoll, and Cornelia I. Bargmann. Mechanosensory signalling in *C. elegans* mediated by the glr-1 glutamate receptor. Nature, 378(6552):78–81, 1995.

59. Maria A. Lim, Jyothsna Chitturi, Valeriya Laskova, Jun Meng, Daniel Findeis, Anne Wiekenberg, Ben Mulcahy, Linjiao Luo, Yan Li, Yangning Lu, Wesley Hung, Yixin Qu, Chi-Yip Ho, Douglas Holmyard, Ni Ji, Rebecca McWhirter, Aravinthan D. T. Samuel, David M. Miller, Ralf Schnabel, John A. Calarco, and Mei Zhen. Neuroendocrine modulation sustains the *C. elegans* forward motor state. eLife, 5, 2016.

60. Yuan Wang, Xiaoqian Zhang, Qi Xin, Wesley Hung, Jeremy Florman, Jing Huo, Tianqi Xu, Yu Xie, Mark J. Alkema, Mei Zhen, and Quan Wen. Flexible motor sequence generation during stereotyped escape responses. eLife, 9, 2020.

61. Lav R Varshney, Beth L Chen, Eric Paniagua, David H Hall, and Dmitri B Chklovskii. Structural properties of the *Caenorhabditis elegans* neuronal network. PLoS computational biology, 7(2):e1001066, 2011.

62. Kevin S. Chen, Rui Wu, Marc H. Gershow, and Andrew M. Leifer. Continuous odor profile monitoring to study olfactory navigation in small animals. eLife, 12:e85910, 2023. ISSN 2050-084X. doi: 10.7554/eLife.85910.

63. S. Faumont and Shawn R. Lockery. The awake behaving worm: Simultaneous imaging of neuronal activity and behavior in intact animals at millimeter scale. Journal of Neurophysiology, 95(3):1976–1981, 2006.

64. Francesco Randi and Andrew M. Leifer. Measuring and modeling whole-brain neural dynamics in *Caenorhabditis elegans*. Current Opinion in Neurobiology, 65:167–175, 2020. ISSN 0959-4388.

65. Jeffrey P. Nguyen, Frederick B. Shipley, Ashley N. Linder, George S. Plummer, Mochi Liu, Sagar U. Setru, Joshua W. Shaevitz, and Andrew M. Leifer. Whole-brain calcium imaging with cellular resolution in freely behaving *Caenorhabditis elegans*. Proceedings of the National Academy of Sciences, 113(8):E1074–E1081, 2016.

66. Charles Fieseler, Itamar Lev, Jalaja Madhusudhanan, Zihao Zhai, Siegfried Schwartz, and Manuel Zimmer. Barlowtrack: A self-supervised framework for zero-shot multi-object cell tracking. bioRxiv, 2025. doi: 10.1101/2025.11.10.687595.

67. Eduard Bokman, Christian O. Pritz, Rotem Ruach, Eyal Itskovits, Hadar Sharvit, and Alon Zaslaver. Intricate response dynamics enhances stimulus discrimination in the resource-limited *C. elegans* chemosensory system. BMC Biology, 22(1), 2024.

68. John White. Clues to basis of exploratory behaviour of the *C. elegans* snout from head somatotropy. Philosophical Transactions of the Royal Society B, 373(1758):20170367, 2018.

69. Shay Stern, Christoph Kirst, and Cornelia I. Bargmann. Neuromodulatory Control of Long-Term Behavioral Patterns and Individuality across Development. Cell, 171(7):1649–1662.e10, 2017.

70. Jayeeta Basu and Katherine Nagel. Neural circuits for goal-directed navigation across species. Trends in Neurosciences, 47(11):904–917, 2024.

71. Edvard I. Moser, Emilio Kropff, and May-Britt Moser. Place Cells, Grid Cells, and the Brain’s Spatial Representation System. Annual Review of Neuroscience, 31(1):69–89, 2008.

72. Guangwei Si, Jessleen K Kanwal, Yu Hu, Christopher J Tabone, Jacob Baron, Matthew Berck, Gaetan Vignoud, and Aravinthan DT Samuel. Structured odorant response patterns across a complete olfactory receptor neuron population. Neuron, 101(5):950–962, 2019.

73. Matthew S. Creamer, Kevin S. Chen, Andrew M. Leifer, and Jonathan W. Pillow. Correcting motion induced fluorescence artifacts in two-channel neural imaging. PLoS Computational Biology, 18(9): e1010421–, 2022.

74. Raymond YN Lee, Elizabeth R Sawin, Martin Chalfie, H Robert Horvitz, and Leon Avery. Eat-4, a homolog of a mammalian sodium-dependent inorganic phosphate cotransporter, is necessary for glutamatergic neurotransmission in *Caenorhabditis elegans*. Journal of Neuroscience, 19(1):159–167, 1999.

75. Aixa Alfonso, Kiely Grundahl, Janet S Duerr, He-Ping Han, and James B Rand. The *Caenorhabditis elegans* unc-17 gene: a putative vesicular acetylcholine transporter. Science, 261(5121):617–619, 1993.

76. Esther Serrano-Saiz, Richard J Poole, Terry Felton, Feifan Zhang, Estanisla Daniel De La Cruz, and Oliver Hobert. Modular control of glutamatergic neuronal identity in c. elegans by distinct homeodomain proteins. Cell, 155(3):659–673, 2013.

77. Laura Pereira, Paschalis Kratsios, Esther Serrano-Saiz, Hila Sheftel, Avi E Mayo, David H Hall, John G White, Brigitte LeBoeuf, L Rene Garcia, Uri Alon, and Oliver Hobert. A cellular and regulatory map of the cholinergic nervous system of c. elegans. Elife, 4:e12432, 2015.

78. Guangshuo Ou, Makato Koga, Oliver E Blacque, Takashi Murayama, Yasumi Ohshima, Jenny C Schafer, Chunmei Li, Bradley K Yoder, Michel R Leroux, and Jonathan M Scholey. Sensory ciliogenesis in *Caenorhabditis elegans*: assignment of ift components into distinct modules based on transport and phenotypic profiles. Molecular biology of the cell, 18(5):1554–1569, 2007.

79. Seth R Taylor, Gabriel Santpere, Alexis Weinreb, Alec Barrett, Molly B Reilly, Chuan Xu, Erdem Varol, Panos Oikonomou, Lori Glenwinkel, Rebecca McWhirter, Abigail Poff, Manasa Basavaraju, Ibnul Rafi, Eviatar Yemini, Steven J Cook, Alexander Abrams, Berta Vidal, Cyril Cros, Saeed Tavazoie, Nenad Sestan, Marc Hammarlund, Oliver Hobert, and David M Miller, III. Molecular topography of an entire nervous system. Cell, 184(16):4329–4347, 2021.

80. Juliette Ben Arous, Yoshinori Tanizawa, Ithai Rabinowitch, Didier Chatenay, and William R Schafer. Automated imaging of neuronal activity in freely behaving *Caenorhabditis elegans*. Journal of neuroscience methods, 187(2):229–234, 2010.

81. Serge Faumont, Gary Rondeau, Tod R Thiele, Kristy J Lawton, Kathryn E McCormick, Matthew Sottile, Oliver Griesbeck, Ellie S Heckscher, William M Roberts, Chris Q Doe, and Shawn R Lockery. An image-free opto-mechanical system for creating virtual environments and imaging neuronal activity in freely moving *Caenorhabditis elegans*. PloS one, 6(9):e24666, 2011.

82. F. Pedregosa, G. Varoquaux, A. Gramfort, V. Michel, B. Thirion, O. Grisel, M. Blondel, P. Prettenhofer, R. Weiss, V. Dubourg, J. Vanderplas, A. Passos, D. Cournapeau, M. Brucher, M. Perrot, and E. Duchesnay. Scikit-learn: Machine learning in Python. Journal of Machine Learning Research, 12:2825–2830, 2011.

